# Composition and functionality of bacterioplankton communities in marine coastal zones adjacent to finfish aquaculture

**DOI:** 10.1101/2022.03.07.483349

**Authors:** R.R.P. Da Silva, C.A. White, J.P. Bowman, D.J. Ross

## Abstract

Finfish aquaculture is one of the fastest-growing primary industries globally and is increasingly common in coastal ecosystems. Bacterioplankton is ubiquitous in marine environment and respond rapidly to environmental changes. However, little is known about the effect of the aquaculture in the bacterioplankton community. This study aims to examine aquaculture effects in the composition and functional profiles of the bacterioplankton community using amplicon sequencing along a distance gradient from two finfish leases in a marine embayment. Our results revealed natural stratification in bacterioplankton strongly associated to NOx, conductivity, salinity, temperature and PO_4_. Among the differentially abundant bacteria in leases, we found members associated with nutrient enrichment and aquaculture activities. Abundant predicted functions near leases were assigned to organic matter degradation, fermentation, and antibiotic resistance. This study provides a first effort to describe changes in the bacterioplankton community composition and function due to finfish aquaculture in a semi-enclosed and highly stratified embayment.

## 1. Introduction

Aquaculture is one of the fastest-growing primary industries worldwide and is increasingly common in coastal ecosystems. Aquaculture of carnivorous fish species, such as Atlantic salmon (*Salmo salar*) is feed-additive, with a significant portion of feed flowing to the environment as metabolic waste products, faeces and waste feed (Navarro et al., 2008, Burridge et al., 2010, Zhang et al., 2015). As industries expand there is increasing concern regarding potential effects that this waste can cause to surrounding environments. Key to sustainable aquaculture operations in marine coastal systems is a better understanding around how pelagic environments assimilate and mitigate additional nutrient loadings.(Asami et al., 2005, á Norði et al., 2011, Belias et al., 2003, Elizondo-Patrone et al., 2015). There are two main pathways of enrichment in pelagic environments, being the solid and dissolved waste streams (Bouwman et al., 2013, Tsukamoto et al., 2008, Wang et al., 2012, Amirkolaie, 2011). The solid waste stream consists of waste feed and faecal material and is available to pelagic primary consumers as the solid debris settles out of the water column through sedimentation (Macleod et al. 2004). In contrast, metabolic waste products, in particular bioactive inorganic nitrogen compounds such as ammonia and nitrate, are released into the water column as dissolved nutrients and available for assimilation by planktonic organisms (Wang et al. 2012).

Bacterial communities are known to be ubiquitous in the marine environment making up a large fraction of the planktonic ecosystem. Bacterioplankton communities (i.e., bacterial communities of the water column) play an important role in nutrient cycling and biogeochemical processes (Azam et al., 1983, Pomeroy et al., 2007, Fuhrman and Steele, 2008). Bacterioplankton are highly dynamic, responding to natural or anthropogenic changes (Newell et al., 2013, Bowman and Ducklow, 2015, Yu et al., 2014, Reji et al., 2020). Previous studies on sediments beneath aquaculture operations have identified changes in the bacterial community due to nutrient enrichment (McCaig et al., 1999, Bissett et al., 2006, Dowle et al., 2015, Rubio-Portillo et al., 2019). However, bacterioplankton community change across enrichment gradients caused by aquaculture activities have yet to be fully defined. Olsen et al. (2017) studying the response of the bacterial community to nutrient input in surface waters adjacent to fish cages in southern Chile found an impact on community structure, although changes in diversity were not observed. While this suggests stability and functional resilience in bacterioplankton communities, there are very few other studies that examined response, particularly across different receiving environments (Yoshikawa and Eguchi, 2013, Elizondo-Patrone et al., 2015, Girvan et al., 2005). Aquatic ecosystems are complex, with cumulative or synergetic effects frequently acting in a multifactorial way. Natural variation of the system and management of aquaculture inputs are key factors to consider when assessing the impact of waste on the pelagic environment (Wu, 1995, Buschmann et al., 2006). Therefore, it is important to study the responses of bacterial community to environmental variations to enhance our ability to understand changes caused by both natural and anthropogenic environmental changes.

The advancement of molecular-based techniques has greatly advanced the field of microbial ecology (Pace, 1997, Gilbert et al., 2010, Caporaso et al., 2011, Yarza et al., 2014, Martínez-Porchas and Vargas-Albores, 2017, DeLong et al., 1993). The high-throughput sequencing of 16S rRNA has been used as a powerful tool to study the composition of bacterial community from a variety of habitats (Caporaso et al., 2011, Logares et al., 2014, Morelan, 2019, Han et al., 2017). Additionally, the availability of genomic data now allows inference of functional capabilities based on taxon identification and relative abundances (Langille et al., 2013, Sevigny et al., 2019, Asshauer et al., 2015, Nagpal et al., 2016, Ortiz-Estrada et al., 2019). Previous studies using of 16S rRNA gene amplicon sequencing have showed a response of the microbial community and function to aquaculture activities in various aquatic systems (Keeley et al., 2018, Kolda et al., 2020, Ortiz-Estrada et al., 2019, Martins et al., 2018, Moncada et al., 2019, Frühe et al., 2021). Cordier (2020) could detect bacterial compositional and functional changes in the sediments close to aquaculture, where molecular-based techniques performed similarly to traditional microscopy-based techniques in predicting organic enrichment. However, natural local characteristics such as water column stratification and natural variation in environmental factors have major effects on the bacterial community structure and function (Labbate et al., 2016, Martiny et al., 2006, Zorz et al., 2019, Da Silva et al., 2021). The strong influence of the environment can make comparisons between studies undertaken in different locations difficult. However, the more we understand the natural variability across different marine systems, the better placed we are to understand common responses in bacteria community different perturbations, both natural and anthropogenic.

Macquarie Harbour is highly stratified fjord-like system with restricted water exchange with the ocean. The water column is characterized by a brackish tannin-rich water layer on top of more saline marine waters. The major freshwater source flows from the southern end carrying high amounts of organic matter with mean monthly flows varying according to the season (< 100 m^3^ sec^™1^ in summer and early autumn, and 500 m^3^ sec^™1^ late autumn, winter, and spring) (Hartstein et al., 2019, Carpenter et al., 1991, Teasdale et al., 2003). Marine water enters through a narrow and shallow inlet in the north, allowing the renewal of the dissolved oxygen (DO) in the deeper waters. Marine incursions into the Harbour are likely driven by forces such as tide, wind, atmospheric pressure and freshwater inputs (Cresswell et al., 1989, Hartstein et al., 2019). The Harbour has a history of anthropogenic impacts including mining, damming and aquaculture (King and Tyler, 1982, Crawford et al., 2003, Koehnken, 2005, Kirkpatrick et al., 2019). Pelagic DO concentrations naturally drop with increasing depth reaching around 4 to 6 mg O_2_ L^-1^ at bottom layers. However, a recent report observed a considerable decline, with levels now 2 to 4 mg O_2_ L^-1^ or lower below 25 m in depth (Ross et al., 2021). This decline is thought to be related to a reduction in the frequency and scalability of DO recharge along with a high biological oxygen demand due to input of organic matter from fish farms. Additionally, previous studies have suggested that pelagic microbial oxygen is likely to be an important determinant of the decline in pelagic oxygen within the Harbour (Revill et al., 2016, Ross et al., 2021, Da Silva et al., 2021). However, the impact of fish farm inputs on the bacterioplankton community remain unclear.

For this reason, the main goal of this study is to better characterize the relationship between the bacterioplankton community and proximity to aquaculture within Macquarie Harbour. This will provide a starting point for further studies to examine the effects of aquaculture in the bacterioplankton community, especially in a region with restricted water exchange. The study aims (i) to examine the composition and distribution of bacterioplankton community and (ii) to assess the influence of aquaculture activity on microbial community structure and distribution. Based on previous studies and taking into account the hydrodynamics of the Harbour, we hypothesize that the effects of aquaculture in the bacterioplankton community will be localized.

## 2. Material and methods

### Location and sample collection

In December of 2019, sampling was conducted at two fish farm leases (“Lease 1” and “Lease 2”) and control stations (“Control”) in Macquarie Harbour (42°18’13.8”S; 145°22’22.7”E), Tasmania, Australia (Fig. 1). At each of the leases, samples were collected at sites on replicate transects radiating from a fully stocked Atlantic salmon cage (*Salmo salar*) fed a pellet diet using 10 L Niskin bottles (Tables S1 and S2). Seawater was collected from the surface (0-4 m), middle waters (10 – 15 m) and bottom waters (20 – 37 m) at each site except for the site inside the cage. Each transect (Lease 1: n = 2; Lease 2: n=3) included 4 distances (inside the cage + 3 distances) at the surface, and 3 distances at middle and bottom layers. Sampling over distance was carried out by letting the boat drift along the direction of the current. Three control stations (>1500 m away from cages) were sampled at the three depths mentioned above. Genetic material was collected on 0.22 µM polyethersulfone Sterivex^™^ filter cartridges (Millipore, Darmstadt, Germany) using a multichannel peristaltic pump. Each sample was stored at −80°C until DNA extraction. Haversine’s formula was used to calculate the distance between sample points by use of latitude and longitude data (Sinnott, 1984).

**Fig 1.**
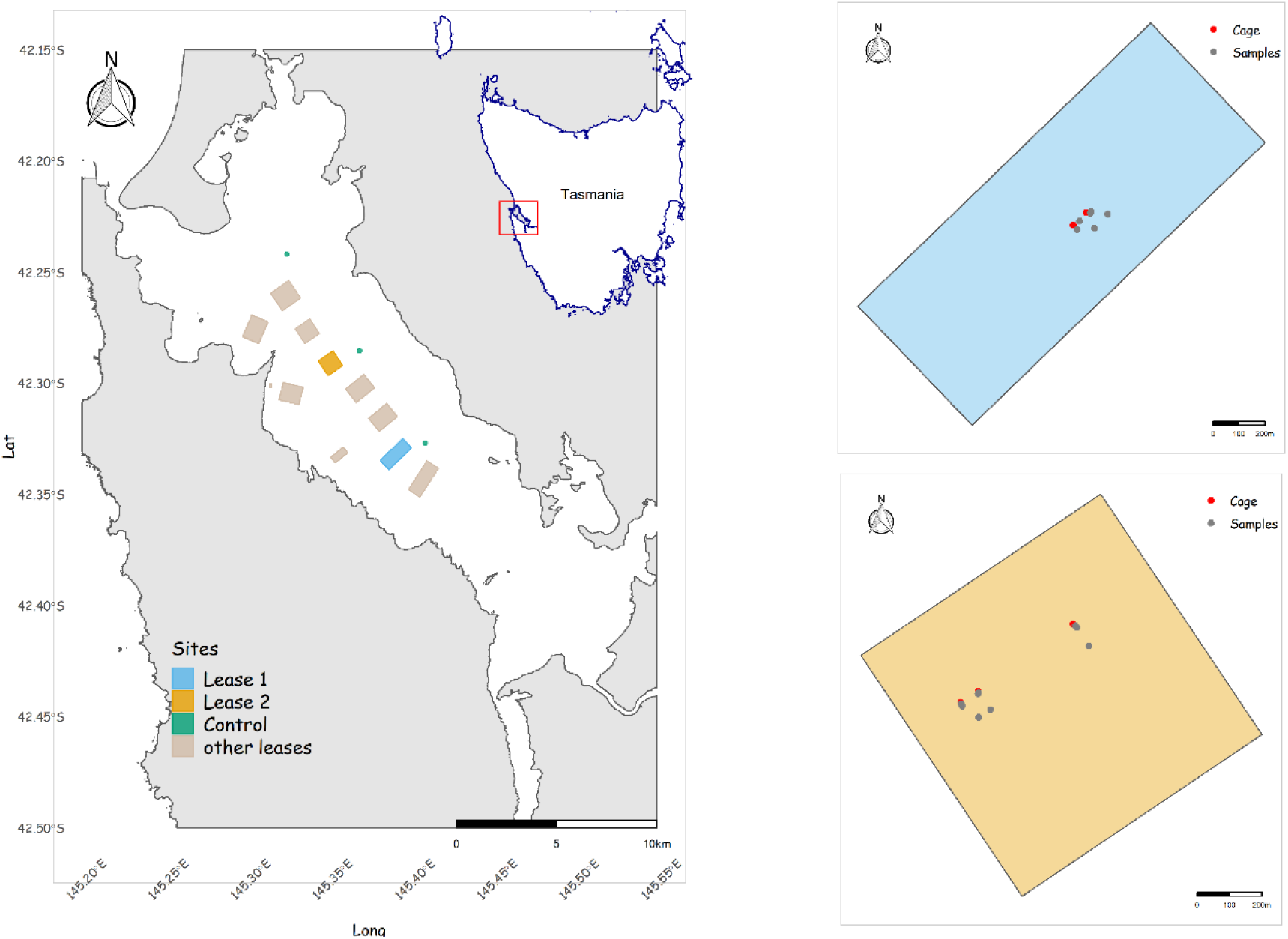
Map on the left displays the locations of sites sampled within this study. Green points are control sites and squares are leases located in the Macquarie Harbour. Sampled leases are the blue (Lease 1) and orange (Lease 2) ones. The two maps on the right show the sampled leases where the red points are the cages and the grey points are the samples collected along a distance gradient.

### Environmental variables

Physiochemical parameters including temperature (°C), dissolved oxygen (DO, mg L-1), turbidity (NTU) and salinity were measured *in situ* at each site using a YSI 6600 V2 Multi Parameter Water Quality Sonde with a YSI 650 MDS logger. Nutrient analyses were assayed using a Bran + Luebbe AA3 HR segmented flow analyzer following standard spectrophotometric methods (Grasshoff et al., 2009). Detection limits for NOx (NO _3_^-^+ NO _2_^-^), nitrite (NO _2_^-^) were 0.015 μmol/L, for phosphate (PO _4_^-3^) to 0.01 μmol/L, and for ammonium (NH_4_^+^) to 0.004 μmol/L.

### DNA extraction and sequencing

DNA was extracted using a modified phenol:chloroform:isoamyl based DNA extraction protocol of the DNeasy PowerWater SterivexTM Kit (MoBio-Qiagen, vHilden, Germany) as described in Appleyard et al. (2013). Amplicon sequencing targeted the V1 to V3 region of the bacterial 16S rRNA gene (27F–519R) (Lane, 1991, Lane et al., 1985). Libraries were separately generated and sequenced for each sample at the Ramaciotti Centre for Genomics (University of New South Wales, Sydney) using 300 bp paired-end sequencing on the MiSeq platform (Illumina Inc., USA). Raw sequence reads and metadata are publicly available at the Sequence Read Archive under the BioProject accession PRJNA803985 (https://www.ncbi.nlm.nih.gov/sra/PRJNA803985).

### Bioinformatic Pipeline

Paired end sequences of the 16S rRNA gene amplicons were analyzed through the DADA2 package pipeline, version 1.20.0 (Callahan et al., 2016) in R 4.1.0 environment (R Core Team, 2021). Reads were filtered with the following parameters: truncLen = c(290, 225), truncQ = 2, maxEE = c(2, 2). Merging of the forward and reverse reads was done with the mergePairs function using the default parameters (minOverlap = 12, maxMismatch = 0). Chimeras were removed using *removeBimeraDenovo* with default parameters. Taxonomic classification was performed using the *assignTaxonomy* function using the parameters (tryRC = FALSE, minBoot = 50) with the SILVA v138 database (Yilmaz et al., 2014) using the naive Bayesian classifier method described in Wang et al. (2007). Mitochondria-derived sequences were removed from the original dataset, obtaining 24575 ASVs. ASV abundance tables, taxonomy and environmental data were imported into the phyloseq R package version 1.36.0 (McMurdie and Holmes, 2013). To reduce the data complexity (Cao et al., 2021), bacterial ASVs were filtered to keep taxa observed at least 2 times in at least 2 samples resulting in 3260 ASVs for further analyses (2.7 % of total reads removed).

### Functional Annotations

Functional composition were predicted using PICRUSt2 version 2.4.1 (Douglas et al., 2020). First, the nearest sequenced taxon index (NSTI) values are obtained as a metric to determine the degree to which ASVs in a sample are related to available reference genomes and environmentally sampled organisms. Then, gene content inference per ASV was performed using hidden-state prediction (HSP) approaches. Before prediction, data was normalized by their 16S rRNA gene copy number abundance. Finally, metagenome inference was used to generate MetaCyc pathway predictions (Caspi et al., 2018). MetaCyc pathways were linked to their respective secondary superclass levels using the tools available at https://metacyc.org.

### Data Analysis and Statistics

All data analysis and statistical tests were conducted in R 4.1.0 environment (R Core Team, 2021). Environmental variables were explored using the tidyverse package v.1.3.1 (Wickham et al., 2019). Bacterial and functional communities were centered log-ratio (CLR) transformed to address the negative correlation bias intrinsic to compositional data (Gloor et al., 2017) using CoDaSeq v.0.99.6 (Gloor, 2016) and zCompositions v.1.3.4 (Palarea-Albaladejo and Martín-Fernández, 2015) packages. To visualize differences among samples based on environmental and community (ASVs composition and functional annotations) data, Principal Composition Analysis (PCA) followed by K-means clustering and distance-based Redundancy Analysis (db-RDA) on Aitchison distance (Aitchison, 1986) were performed, respectively. PCA were implemented using tidymodels v.0.1.3 (Kuhn and Wickham, 2020), K-mean clustering and validation methods using factoextra v.1.0.7 (Kassambara and Mundt, 2020), and db-RDA using vegan v.2.5.7 (Oksanen et al., 2018) packages. Environmental variables which could explain variation found on db-RDA ordinations were fitted using both the *envfit*, and *ordistep* functions, the latter with forward and backward stepwise selections. Significant correlation between environmental variables and ordination (p-value < 0.01 in both analyses) were plotted as vectors (arrows) whose direction and length indicate the environmental variable gradient and the correlation, respectively. Prior to both ordination methods, autocorrelated environmental variables (i.e., r > 0.7, p-value ≤ 0.05) were combined into one for downstream analyses.

To verify the magnitude of variation along a gradient of environmental changes between taxonomic and functional profiles, a correlation between bacterial community and functional annotation distance matrices was calculated using Mantel tests (Spearman rank correlation and 9999 permutations). To evaluate the difference in community composition between leases and control sites, ASVs and functional annotations were subjected to two widely used statistical methods: ALDEx2 (Fernandes et al., 2013) and DESeq2 (Love et al., 2014) packages. The combination of these two methods aims to overcome the drawbacks found in differential abundance methods, such as low power and high False Discovery Rate (FDR) (Hawinkel et al., 2019). For ALDEx2, clr-transformed abundance table was used as the input for the analyses, whereas DESeq2 used variance-stabilized data to perform the evaluation. ASVs and pathways were considered differentially abundant if they satisfied the following assumptions: an effect size > |1.5| in ALDEx2 and an adjusted P value (FDR) < 0.05 and a log2foldchange > |2| in DESeq2 analysis. The package ggtern v.3.3.5 (Hamilton and Ferry, 2018) was used for data visualization. Then, the differentially abundant ASVs and pathways were used as model input data (clr-transformed count table) to verify the relationship with the environmental variables used in the previous ordination analyses (i.e., NH_4_, NOx, turbidity, and oxygen) as described below.

Random forest regression modelling was implemented through tidymodels v.0.1.3 (Kuhn and Wickham, 2020) and randomForest v.4.6.14 (Liaw and Wiener, 2002) packages to test the ability of the differentially abundant ASVs to predict estimates of the variability of the aforementioned environmental variables. For this process normalized clr-transformed data was split into training and testing sets in a proportion of ¾. To tune the hyperparameters (mtry and min_n), a k-fold cross validation scheme (k = 10 with 3 repeats) with 501 trees of the training set was used to estimate the best model based on root-mean-square error (RMSE) metric. The model was finalized using the *finalize_workflow* and *select_best* functions. After that, the model performance was tested in the training set and the metrics RMSE and R2 were collected using the *collect_metrics* function. Variable importance was measured by the residual sum of squares. Model predictions and variable importance were extracted with the *collect_predictions* function and vip v.0.3.2 package (Greenwell and Boehmke, 2020), respectively and visualized using tidyverse package v.1.3.1 (Wickham et al., 2019). Code used for data analyses in this study can be found in the associated GitHub repository (https://github.com/ricrocha82/MH_2019_16s).

## 3. Results

### Environmental Variables

The samples from leases and control sites showed a uniform trend in most environmental variables when distances from cages were considered, however for NH_4_, NO_2_ and NOx, concentrations were highly variable at both leases (Fig. 2; Tables S2, S3 and S4). Despite this variation, NH_4_ level was observed to decrease as distance from cage increased at the surface of Lease 2 (2.84 ± 0.47 μM to 0.1 ± 0.03 μM) and NOx decreased in concentration with distance in the middle waters of Lease 1 (6.38 ± 1.22 μM to 1.29 ± 0.46 μM). Also, at Lease 1, NO_2_ showed an increase in concentration as distance increased after ∼18 m (0.04 ± 0.05 μM to 0.28 ± 0.04 μM).

**Fig 2.**
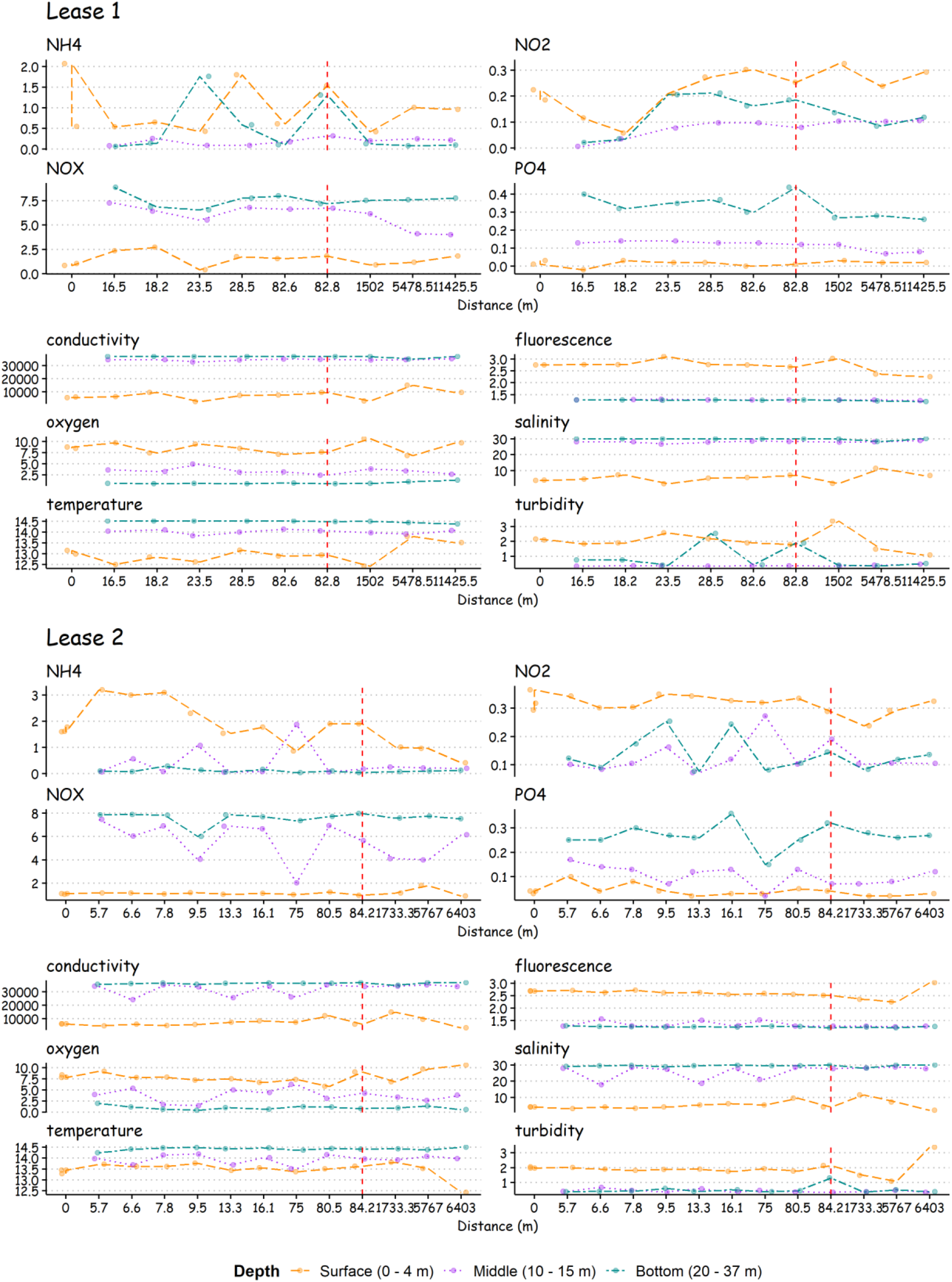
Environmental variables along a distance gradient in each lease area. Each line in the graph displays environmental variable measurements in each depth. Red dashed line shows the farthest distance from the cage. (i.e., end of cage influenced zone). After this line cage site distances are displayed. Distance 0 m indicates sample from inside the cages (2 cages in Lease 1 and 3 cages in Lease 2). Dissolved nutrients (μmol l^-1^), conductivity, fluorescence, oxygen (mg/l), temperature (°C), turbidity (NTU)

When depth is considered, stratification is clear for all environmental variables at all leases and control sites (Fig. 3 – A; Tables S1-S4). Despite the variation observed in nutrient concentrations, NH_4_ (1.53 ± 0.85 μM to 0.3 ± 0.48 μM) and NO_2_ (0.28 ± 0.08 μM to 0.14 ± 0.07 μM) decreased with depth whereas NOx (1.29 ± 0.51 μM to 7.57 ± 0.63 μM) and PO_4_ (0.03 ± 0.02 μM to 0.3 ± 0.07 μM) displayed the opposite trend. Similarly to NOx and PO_4_, conductivity (6914.73 ± 2781.28 to 36573.31 ± 548.15), salinity (5 ± 2.26 to 29.7 ± 0.5) and temperature (13.28 ± 0.41 °C to 14.45 ± 0.07 °C) were greater at depth, whereas fluorescence (2.68 ± 0.2 to 1.26 ± 0.022 °C), oxygen (8.10 ± 1.14 mg/l to 0.86 ± 0.4 mg/l), PAR (0.62 ± 1 to 0.003 ± 0.001) and turbidity (2 ± 0.4 NTU to 0.72 ± 0.6 NTU) were all lower at depth as seen for NH_4_ and NO_2_ concentrations. NOx was highly correlated to conductivity, salinity, temperature and PO_4_, whereas NH_4_ was correlated to NO_2_ and oxygen to PAR and fluorescence (Fig.S2). Therefore, NOx, NH_4_ and oxygen were used as proxies of their highly correlated peers.

**Fig 3.**
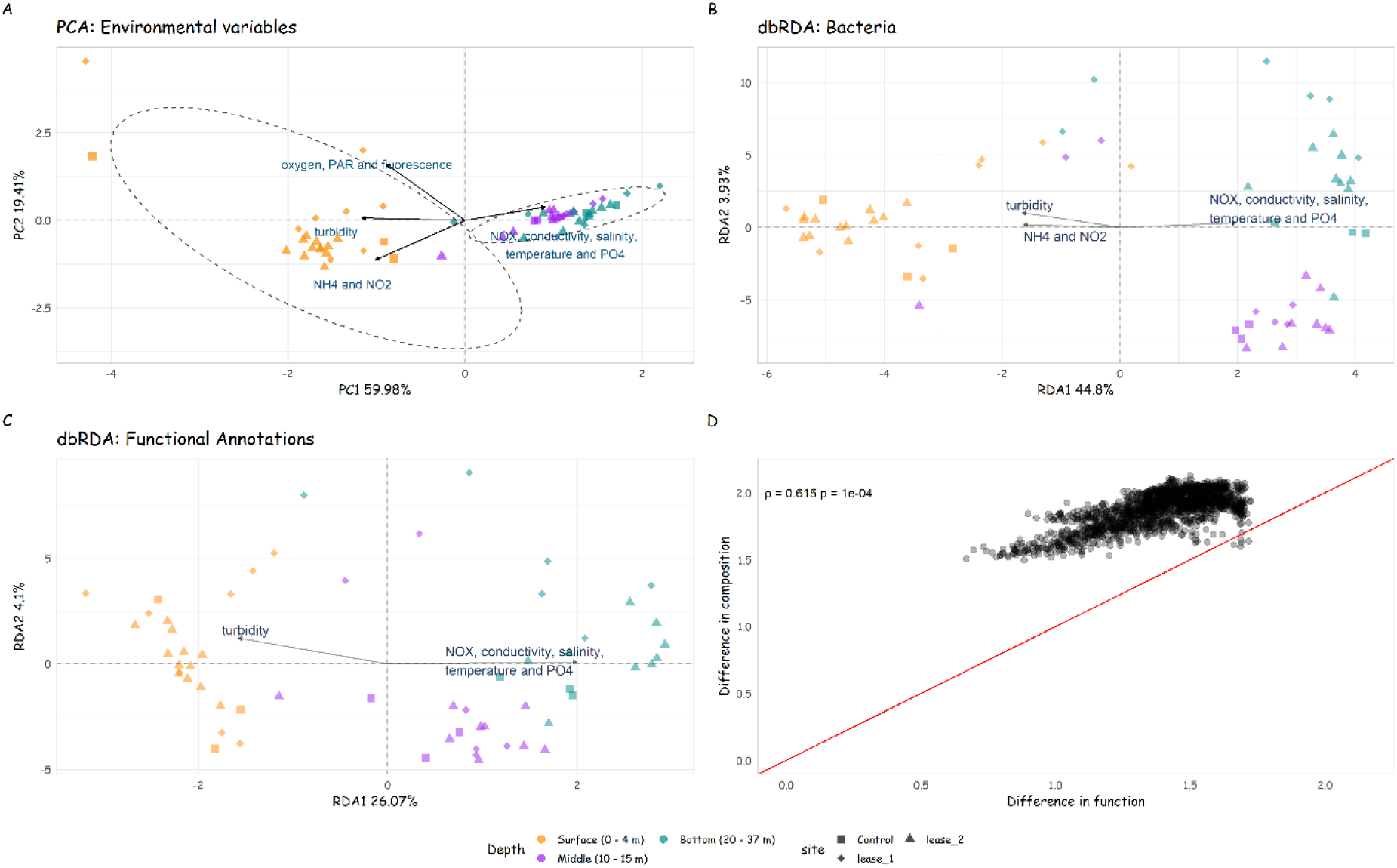
Ordination plots of the samples and vectors (arrows). (**A**) Principal Component Analysis (PCA) of environmental data. Dashed lines highlight the results of K-mean clustering analyses. (**B-C**) Distance-based Redundancy Analysis (db-RDA) of the composition differences of bacterial and functional communities. To avoid multicollinearity, NOx included conductivity, salinity, temperature and PO_4_, and NH_4_ included NO_2_. Vectors indicate a significant correlation between environmental variables and ordination (p < 0.01) using *envfit* function of vegan package. (**D**) Differences in ASV bacterial and functional compositions. Equal changes are shown by the red line. Mantel test shows a significant correlation between the two distances matrices (Spearman rho = 0.6154).

### Bacterial and Functional Communities Structure

At phylum and class levels, bacterial community at all sites was dominated by *Proteobacteria* (Control: 58.4 %; Lease 1: 55.3 %; Lease 2: 57.9 %) and *Gammaproteobacteria* (Control: 30.4 %; Lease 1: 30.7 %; Lease 2: 31.7 %), respectively. *Proteobacteria* was followed by *Bacteroidota* (Control: 15.6 %; Lease 1: 17 %; Lease 2: 18.1 %) at all sites, *Actinobacteriota* (5.6 %) at Control sites and *Campylobacterota* (Lease 1: 7.63 %; Lease 2: 5.45 %) at lease sites. *Gammaproteobacteria* was followed by *Alphaproteobacteria* (Control: 28.1 %; Lease 1: 24.6 %; Lease 2: 21.6 %) and *Bacteroidia* (Control: 15.2 %; Lease 1: 16.7 %; Lease 2: 17.9 %) at all sites. ASVs encompassing the class *Actinobacteria* (Control: 4.3 %; Lease 1: 1.93 %; Lease 2: 1.90 %) were more abundant at control sites whereas *Campylobacteria* ASVs were more abundant at lease sites (Control: 3.3 %; Lease 1: 7.63 %; Lease 2: 5.45 %) (Fig. S3).

Functional annotations were uniformly distributed among leases and control sites. Amino-acid biosynthesis was the most abundant pathway (Control: 17 %; Lease 1: 16.8 %; Lease 2: 16.9 %), followed by nucleotide-biosynthesis (Control: 16.2 %; Lease 1: 16.3 %; Lease 2: 16.3 %) and vitamin-biosynthesis (Control: 9.93 %; Lease 1: 10.2 %; Lease 2: 10.1 %) (Fig. S4). Despite little variation in both communities among sites, depth stratification was evident when ordination analyses were performed (Fig. 3 - B and C). db-RDA explained 49.47% and 31.18% of the variation in bacterial and functional communities, respectively. The ordination of bacterial community distribution was significantly correlated to NOx (r^2^ = 0.76; p < 0.01), NH_4_ (r^2^ = 0.49; p < 0.01) and turbidity (r^2^ = 0.36; p < 0.01) whereas functional profiles were significantly related to NOx (r^2^ = 0.74; p < 0.01) and turbidity (r^2^ = 0.41; p < 0.01) (Table 1).

**Table 1.** Result of redundancy analysis (*ordistep* and *envif* functions) and random forest (rf) of bacterial and functional communities in the Macquarie Harbour. Regarding the db-RDA, we considered significant, if the environmental variable had a p-value < 0.01 in both analyses. To avoid multicollinearity, NOx included conductivity, salinity, temperature and PO_4_. NH_4_ included NO_2_.

| Community | env variable | p-value<br>(ordistep) | p-value<br>(envfit) | r2<br>(envfit) | RMSE (rf) | R2 (rf) |
| --- | --- | --- | --- | --- | --- | --- |
| Bacteria | NO<sub>x</sub>* | 0.005 | 0.001 | 0.76 | 1.14 | 0.87 |
|  | turbidity | 0.005 | 0.001 | 0.36 | 0.66 | 0.44 |
|  | NH<sub>4</sub>** | 0.005 | 0.001 | 0.49 | 0.70 | 0.53 |
| Functional<br>annotation | NO<sub>x</sub>* | 0.005 | 0.001 | 0.74 | 1.23 | 0.82 |
|  | turbidity | 0.005 | 0.001 | 0.41 | 0.53 | 0.59 |

\*NO<sub>x</sub> included conductivity, salinity, temperature and PO<sub>4</sub>.
\*\*NH<sub>4</sub> included NO<sub>2</sub>.

The results indicate that environmental parameters and both bacterial and functional communities are more affected by depth stratification than by aquaculture activity. Moreover, the correlation between the distance matrices of bacterial ASVs and functional compositions suggested a significant relationship (rho: 0.62; p < 0.05). In other words, the differences in the taxonomic profile between samples co-vary with the differences in the functional profile between samples. However, bacterial community showed a more prominent overall shift than the associated functional data (Fig. 3 - D).

### Differential Abundance Among Sites by Depth

To detect key members in bacterial and functional communities at lease and control sites, differential abundance analyses were performed considering the effect of depth stratification observed in the community structure analyses. Therefore, the analyses were conducted by executing pairwise comparisons between sites nested by depth (i.e., surface, middle and bottom water layers). ALDEx2 and DESeq2 found a total of 90 and 167 bacterial ASVs and 82 and 61 functional annotations differentially abundant, respectively. At all depths, ASVs and functional annotations selected by both methods were more abundant in both lease influential zones (i.e., relative abundance > 60%) compared to control site. The difference in bacterial community was pronounced at the bottom and surface layers (Fig. 4). Most of the enriched ASVs found in Lease 1 area inhabited the bottom layers (Fig. S5 and Table S5). The taxon with the highest number of differentially abundant ASVs were those belonging to the genus *Psychromonas* (family *Psychromonadaceae*) followed by *Spirochaeta* 2 (family *Spirochaetaceae*), unknown *Arcobacteraceae* (order *Campylobacterales*), unknown *Bacteroidales, Hypnocyclicus* and an unknown genus (both assigned to the family *Leptotrichiaceae*), *Draconibacterium* and three unknown genera related to the order *Bacteroidales, Sulfurimonas* (family *Sulfurimonadaceae*), *Desulfovibrio* (family *Desulfovibrionaceae*), *Pseudomonas* (family *Pseudomonadaceae*), and *Woeseia* (family *Steroidobacterales*). In contrast, at the surface of Lease 2, *Aliivibrio* (family: *Vibrionaceae*) had the highest number of differentially abundant ASVs (Fig. S6 and Table S6). In the control, taxa related to *Nitrospina* and *Gammaproteobacteria* were more abundant in the bottom layers (Fig. S7 and Table S7).

**Fig 4.**
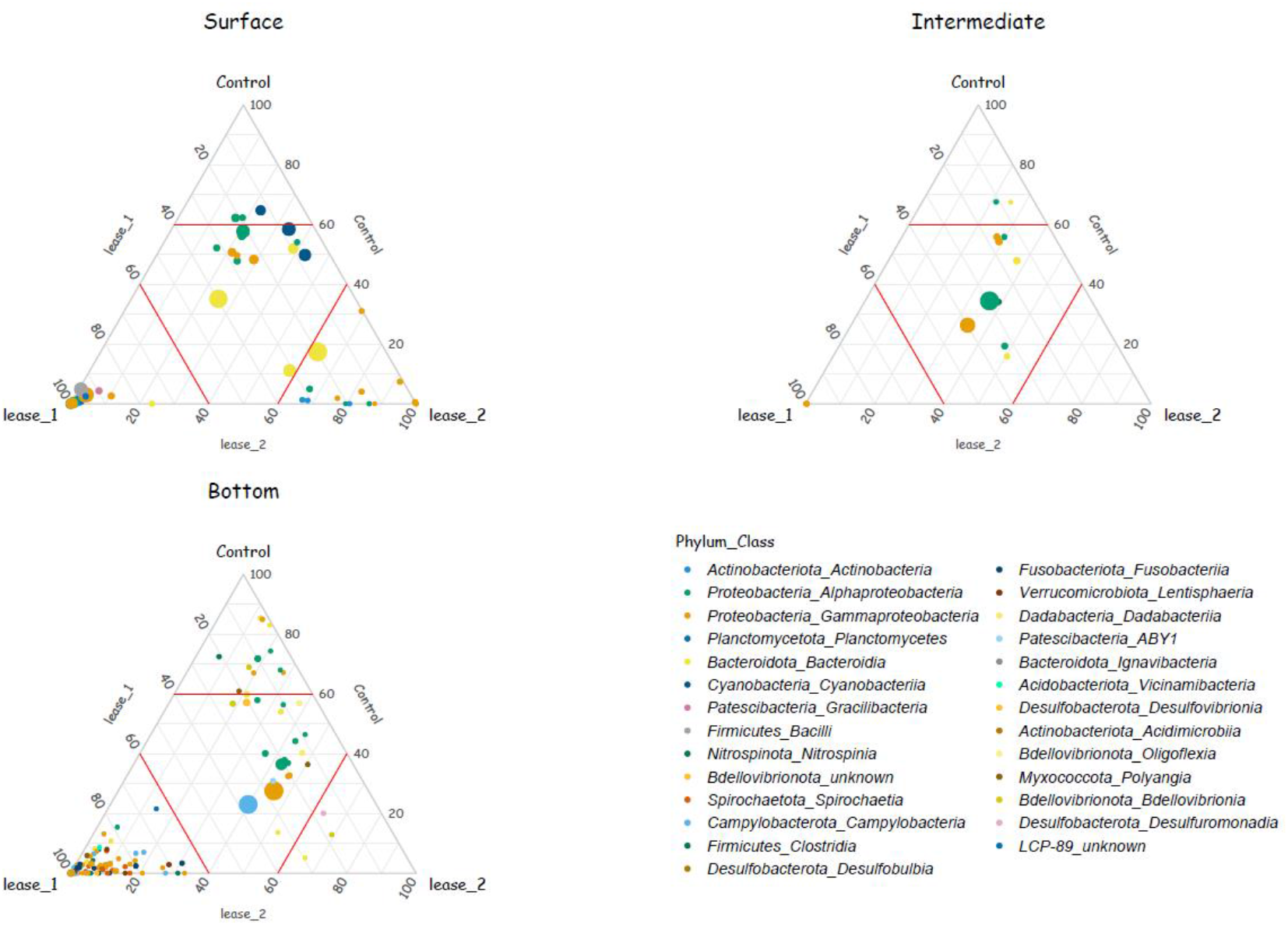
Ternary plots of per-depth relative abundances of differentially abundant ASVs. Each point represents an ASV, and its position indicates the proportion of its relative abundance at each site. Points closer to the ternary plot corners indicate that a greater proportion of the total relative abundance of this ASV was found in this site. Point colors indicate the Phylum and Class of the ASV. The size of each point indicates the mean relative abundance of the ASV in all samples. The red lines inside the ternary plot indicate the 60% level of relative abundance of each of the sites.

Regarding the functional community, the number of enriched functional annotation was the highest at Lease 1, followed by Lease 2 and control sites (Fig. 5, S8 and S9; Tables S8 and S9). In the bottom layer at Lease 1, the analysis detected pathways related with organic matter degradation such as fermentation, carbohydrates, amino-acids, aromatic compounds, and alcohol degradations, as well as those associated with trace metal and macro-nutrient limitations (secondary metabolites, cofactor, vitamins, and amino acids biosynthesis), methanogenesis and cell metabolism. Pathways associated with vitamin degradation and cell metabolism (cell-structure/cell-wall biosynthesis) and those related to alcohol degradation and vitamin biosynthesis were more abundant at the surface and middle layers, respectively. At the control site, lipid biosynthesis and secondary metabolite biosynthesis were differentially expressed at the middle and bottom layers, respectively. These results suggest that the differences observed in bacterioplankton community (16S rRNA gene sequencing) and functional profile (functional annotation: PICRUSt2) at each depth between the lease and control stations are likely to reflect changes in the local environment due to aquaculture activities.

**Fig 5.**
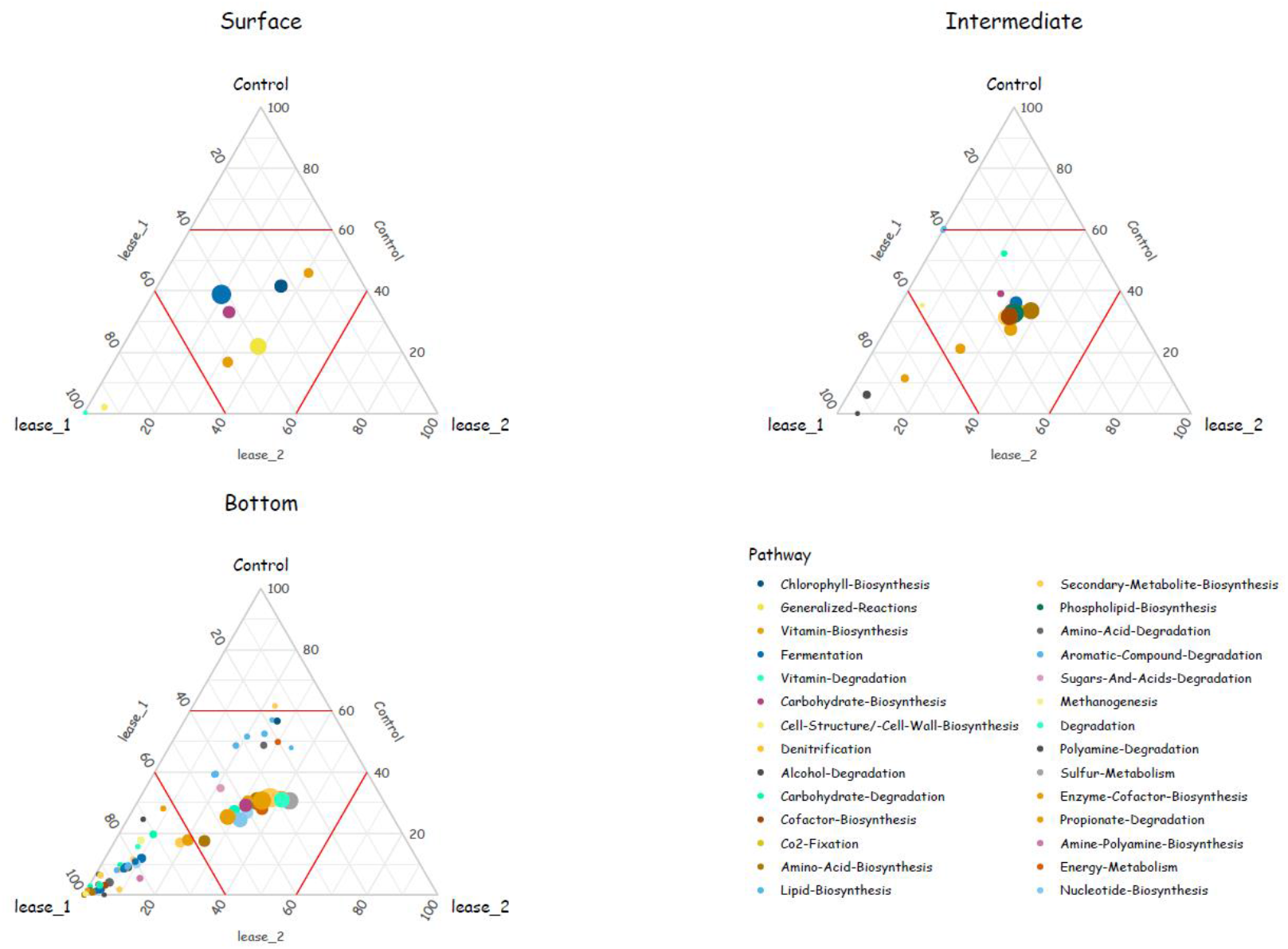
Ternary plots of per-depth relative abundances of differentially abundant functions. Point colors indicate the pathway of the function according to https://metacyc.org. The size of each point indicates the mean relative abundance of the function in all samples. The red lines inside the ternary plot indicate the 60% level of relative abundance of each of the sites.

### Predictive Performance of the Selected ASVs

To measure the importance of these differentially expressed features in predicting the environmental drivers found in the db-RDA analysis (Fig. 3 and Table 1), supervised machine learning was used to estimate the relationship between feature distribution and environmental variable measurements. Then, RF regression models were trained with 257 ASVs and 143 functional annotations. Only NOx (included conductivity, salinity, temperature and PO_4_) concentration prediction models yielded desirable performance for both communities (Table 1). The most important ASV and functional annotation in reducing variance and uncertainty to NOx values were assigned to *Oligoflexaceae* (Fig. 6 and S10; Table S8) and to vitamin biosynthesis, respectively (Fig. 7 and S11; Table S9). Additionally, the most important features in both communities were uniformly distributed among sites, except for some ASVs and functional annotations. For instance, ASVs that could predict NOx concentration, OM27 clade (*Bdellovibrionaceae*) and *Planctomycetes*, were found more abundant (> 60% relative abundance) at the control and Lease 1 samples, respectively. And two functional annotations assigned to methanogenesis and vitamin biosynthesis, both important in NOx prediction, were more enriched in Lease 1 samples.

**Fig 6.**
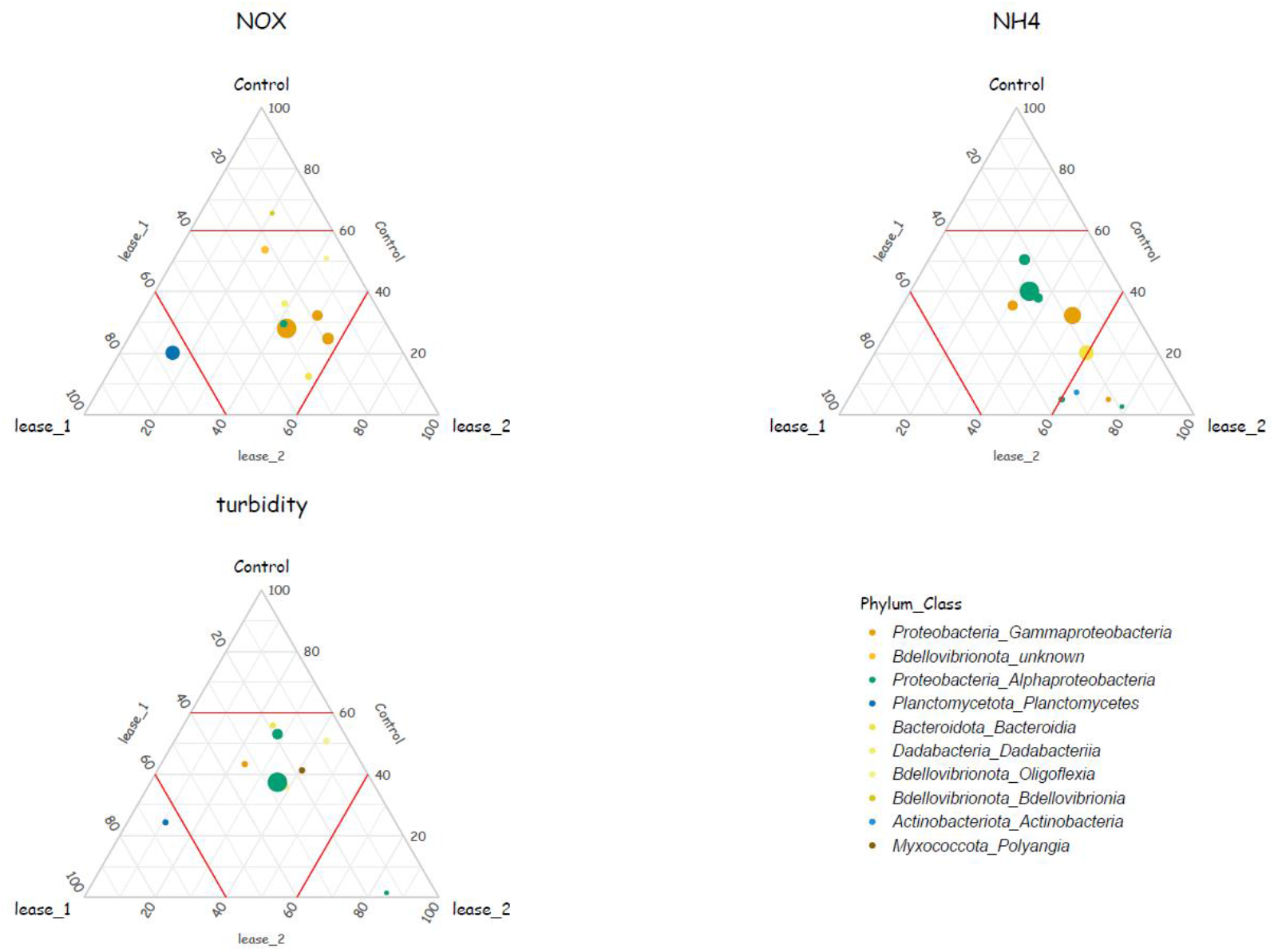
Ternary plot showing the distribution of the 10 most important ASVs based on the residuals sum of square in regression-based prediction model of the environmental drivers of the bacterial community (NH_4_, NOx and turbidity). The size of each point indicates the mean relative abundance of the ASV in all samples. The red lines inside the ternary plot indicate the 60% level of relative abundance of each of the sites. To avoid multicollinearity, NOx included conductivity, salinity, temperature and PO_4_; and NH_4_ included NO_2_.

**Fig 7.**
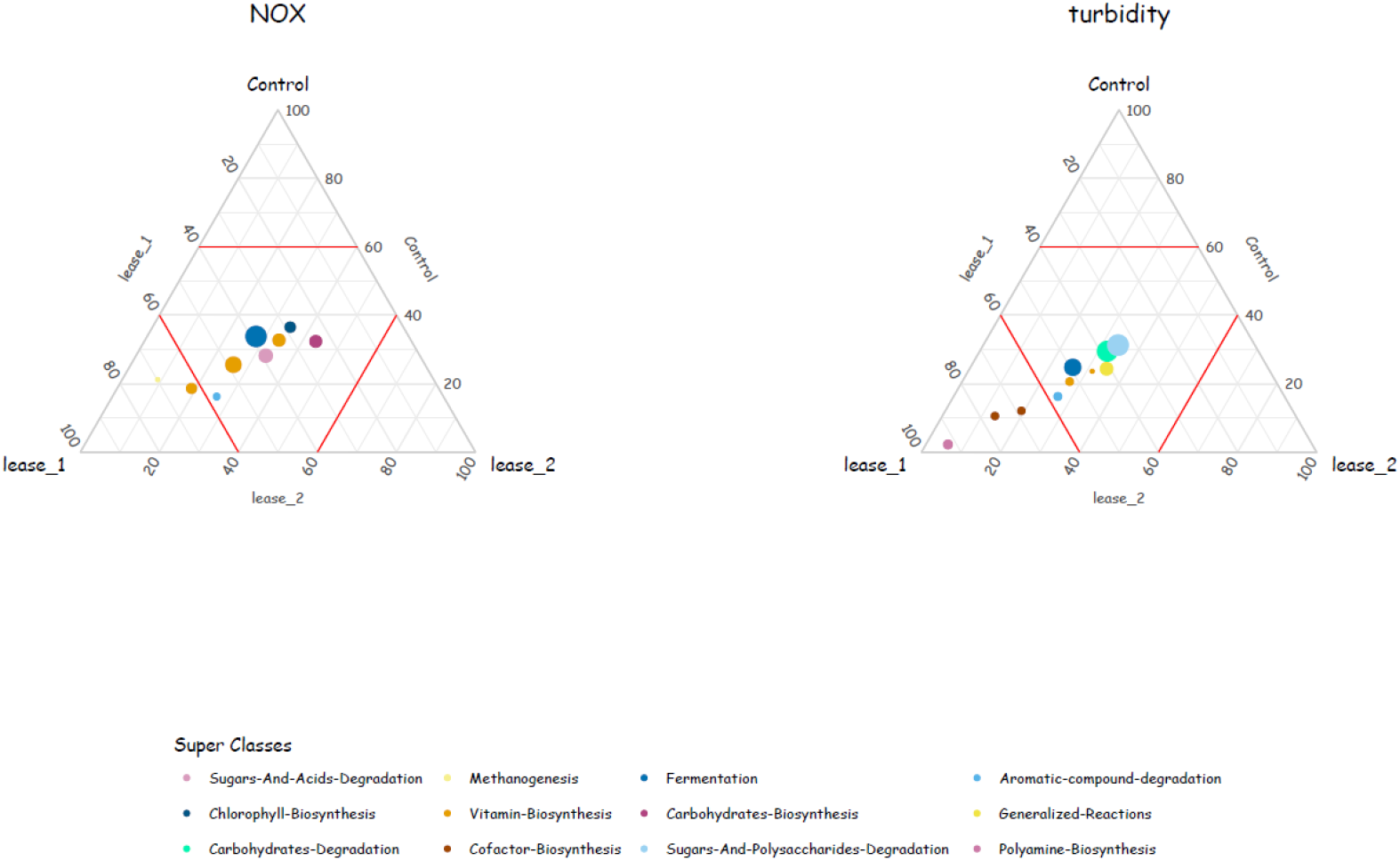
Ternary plot showing the distribution of the 10 most important pathways based on the residuals sum of square in regression-based prediction model of the environmental drivers of the functional community (NOx and turbidity). The size of each point indicates the mean relative abundance of the function in all samples. To avoid multicollinearity, NOx included conductivity, salinity, temperature and PO_4_.

## 4. Discussion

In this study, we used taxonomic and functional profiles to characterize the bacterioplankton community of a coastal embayment containing finfish aquaculture operations in Tasmania (Australia). Using unsupervised and supervised analysis techniques of 16S rRNA bacterial community composition, we observed that the effects of depth stratification on bacterioplankton dominates over aquaculture effects. Despite this natural influence, we also detected differentially abundant ASVs and functional annotations between both marine farm leases (< 85 m) and control sites (> 1500 m), that could be used as good bioindicators for aquaculture activity. According to the random forest regression model, these features could only predict NOx concentrations along with its highly correlated peers (i.e., conductivity, salinity, temperature and PO_4_). Given the environmental heterogeneity and complexity of Macquarie Harbour, it is not possible to draw comprehensive conclusions regarding the impact of salmon aquaculture in the Harbour. Rather, our study provides initial insight into the interaction between salmon aquaculture and the bacterioplankton community, as discussed further below.

### 4.1 Community variation and depth stratification

The main driver of variation in the environmental variables, bacterial community and functional annotations was water column stratification. Macquarie Harbour is considered a highly stratified marine embayment with distinct communities inhabiting each layer in the water column (Da Silva et al., 2021). The higher turbidity and NH_4_ concentration observed at the surface could be related to the tannin-rich waters from Gordon River affecting light penetration which can favor nitrification and organic matter degradation within the water column (Terry, 2001, Hartstein et al., 2019, Bartl et al., 2018, Hou et al., 2018). This could explain the increasing NOx and decreasing DO and NH_4_ concentrations with increasing depth. Additionally, the stratified distribution of both communities (ASVs and functional annotations) could be explained by the variation in the environmental variables, especially those which increased with depth. Most of the differentially abundant ASVs and functional annotation used in the regression analysis to predict NOx concentration were enriched at the bottom layers. Thus, it is unsurprising that the vertical differences in both community distributions observed in this study were strongly correlated to NOx concentration along with conductivity, salinity, temperature and PO_4_ that followed the same vertical profile.

### 4.2 Effects of aquaculture on bacterial community composition

In this study, differential abundance analysis revealed an effect on the taxonomic and functional profiles at both the surface and the near-bottom water layers in lease areas. Most previous aquaculture-environmental impact studies focus on sediments and associated effects to the benthic microbial community (Borja et al., 2009, Fodelianakis et al., 2014, Dowle et al., 2015, Aylagas et al., 2017, Stoeck et al., 2018, Cordier et al., 2018, Keeley et al., 2021). The pelagic environment is less studied and as such the information regarding bacterioplankton response to aquaculture operation is limited, despite their known sensitivity to environmental changes (Wei et al., 2016, Lindh and Pinhassi, 2018, Kolda et al., 2020). Aquaculture activities are known to increase nutrient loading and change trophic status near the fish cages from waste, by-products, fish feces or feed products inputs (Belias et al., 2003, Buschmann et al., 2006, Amirkolaie, 2011). Various microbial groups have been associated with high level of nutrient inputs in the water column (Haro-Moreno et al., 2020, Yang et al., 2019), sediments (Verhoeven et al., 2018, Fruhe et al., 2021, Zhang et al., 2020, Asami et al., 2005, Dully et al., 2021) and land-based aquaculture operations (Mahmoud and Magdy, 2021, Kandel et al., 2014, Aalto et al., 2021, Blancheton et al., 2013) and some have been reported as potential aquaculture impact biomarkers. (Haro-Moreno et al., 2020, Mahmoud and Magdy, 2021, Kandel et al., 2014, Verhoeven et al., 2018, Frühe et al., 2021, Zhang et al., 2020, Aalto et al., 2021, Blancheton et al., 2013, Yang et al., 2019, Rasigraf et al., 2017, Dully et al., 2021). In accordance with these studies, we found differentially abundant ASVs in lease areas that could be related to higher levels of organic matter and the presence of farmed fish. These include members of the taxa *Psychromonadaceae* (*Psychromonas*), *Fusobacteriales (Fusobacteriaceae), Vibrionaceae, Desulfovibrionaceae (Desulfovibrio), Sulfurimonadaceae (Sulfurimonas), Sulfurovaceae (Sulfurovum), Fusibacteraceae (Fusibacter), Rhizobiales, Sphingomonadaceae, Methylotenera, Hyphomonadaceae, Spirosomaceae, Flavobacteriaceae* (NS3a marine group) and *Bacteriovoracaceae*. The abundant ASVs detected around leases in this study correlate well with previous studies conducted in different environments. It suggests that bacterioplankton community from the Macquarie Harbour had similar response to aquaculture activities. Therefore, the techniques applied here could be potential candidates for biomonitoring in coastal salmon aquaculture.

The abundance of members linked to sulphur cycling (i.e., *Desulfovibrionaceae* and *Fusibacter*) have been found in a range of impacted aquatic systems (Blancheton et al., 2013, Aranda et al., 2015, Frühe et al., 2021, Verhoeven et al., 2018, Won et al., 2017). They contribute to organic matter and sulphur-cycling in environments with high levels of organic matter. These findings support previous studies linking nutrient enrichment and sulphur cycle leading to oxygen depletion and the increased production of sulphide (Barton and Hamilton, 2007, Kwon et al., 2016). Moreover, other ASVs detected in higher relative abundance in proximity to aquaculture were those often related to nitrogen and methane cycling. The presence of *Thioglobaceae* (SUP05 cluster), *Beijerinckiaceae* (*Methylocystis*) and *Methylophilaceae* (*Methylotenera*) have been previously reported in both pelagic (Haro-Moreno et al., 2020) and benthic (Fan et al., 2019) aquaculture sites as well as in Atlantic salmon hatcheries (Lokesh et al., 2019, Minich et al., 2020) and could be linked to oxygen concentration and feed inputs (Fan et al., 2019, Hernandez et al., 2015). Interestingly, the predatory *Bacteriovoracaceae* were also found differentially abundant in around leases and could be related to the increase in the number of prey items such as *Vibrio* (Chen et al., 2011, Pineiro et al., 2007). This supports a previous study that detected *Bacteriovoracaceae* as a potential indicator for aquaculture activity (Frühe et al., 2021). It is plausible that these enriched bacterial taxa are most likely related to organic enrichment caused by fish faeces or feed inputs. These distinctive groups of microorganisms represent potential biomarkers and could be tested in further studies to trace and monitor changes caused by aquaculture.

*Psychromonadaceae* (*Psychromonas*), *Fusobacteriales (Fusobacteriaceae), Spirochaetaceae*, and *Leptotrichiaceae, Enterobacterales, Pseudomonadales* (*Pseudomonas*) and *Vibrionaceae* (*Vibrio* and *Aliivibrio*) were also found in higher differential abundance in lease influenced zones compared with control sites. Those ASVs have been reported as members of the marine and freshwater fish gut microbiome (Algammal et al., 2020, Fogarty et al., 2019) often found in bacterial profiling studies using molecular techniques in sediments (Verhoeven et al., 2016, Fruhe et al., 2021, Verhoeven et al., 2018, Kolda et al., 2020), water column (Kopprio et al., 2021, Haro-Moreno et al., 2020), and other aquaculture operation (Martins et al., 2013). Their presence in leases and their absence or low abundance in the control sites suggests an influence of fish faeces on the bacterioplankton community at the aquaculture sites. Additionally, some of these ASVs have been reported as potential pathogens such as *Aliivibrio, Pseudomonas*, and *Arcobacteraceae* (Zarkasi et al., 2016, Bozzi et al., 2021, Wakabayashi, 1980, Loch et al., 2012, Ho et al., 2008) and should be considered in further studies to analyse the role of fish farms as potential reservoirs for pathogens.

### 4.3 The influence of aquaculture activity on functional structure of bacterioplankton

Taxonomic composition of bacterial communities through the use of 16S rRNA gene profiling is useful in showing differences in community composition between sites, as well as providing understanding regarding taxonomic response to the presence of fish cages in the Harbour. However, evidence on potential functional composition can provide important insights into the interaction between metabolic pathways and environmental stress (Mukherjee et al., 2017, Wu et al., 2019, Laroche et al., 2021, Raes et al., 2021). PICRUSt2 is a computational approach to predict the functional composition of a metagenome from the 16S rRNA sequence (Douglas et al., 2020), and has been used widely in previous studies on aquaculture environmental impact (Qu et al., 2021, Vargas-Albores et al., 2019, Ortiz-Estrada et al., 2019, Laroche et al., 2021, Hornick and Buschmann, 2017). By applying PICRUSt2, we are able to explore how the functional community changes with the presence of aquaculture activities.

In this study, PICRUSt2 inferred a methanogenesis pathway from bacterial 16S rRNA amplicon sequencing data. It is well known that methanogenesis is only seen in the archaeal kingdom. Although only bacterial sequences were used as input, this inference could be related to an incorrect clustering with reference archaeal sequences. For these reasons, results from functional prediction tools are useful and insightful but should be interpreted with caution (Douglas et al., 2020). The differences were more pronounced in the near-bottom water layers of leases areas where enriched pathways were associated with organic compounds and vitamin degradation, fermentation, methanogenesis. These pathways are likely related to environment change caused by the presence of fish, including organic enrichment and oxygen depletion, and is consistent with studies on functional alteration along pollution gradients (Galand et al., 2016, Cordier et al., 2019). Interestingly, antibiotic resistance pathway was also found enriched closed to the finfish farms supporting previous studies on aquaculture impact (Tamminen et al., 2011, Hornick and Buschmann, 2017, Chen et al., 2017). The application of antimicrobials can lead to selective pressures in which resistant taxa thrive to the detriment of others (Miranda and Zemelman, 2002). While outputs from functional prediction tools need to be interpreted within their limitations, results from this study suggest that an accumulative effect of organic enrichment, oxygen depletion and antibiotic usage is likely to affect the functional profile of the bacterial community in Macquarie Harbour. Further studies are necessary to test this hypothesis.

Overall, we found that the relative stability of the functional community was greater than the taxonomic community between lease and control sites. The stability detected in this study may be related to the concept of functional redundancy among prokaryotes (Hornick and Buschmann, 2017, Cordier, 2020, Louca et al., 2016, Louca et al., 2018, Bissett et al., 2006, Bissett et al., 2007). This same relationship was observed in previous studies that assessed bacterial community and functional prediction as indicators of anthropogenic pressures such as aquaculture and oil and gas activities (Cordier, 2020, Laroche et al., 2018). Both authors suggested that functional redundancy along with 16S rRNA sequencing could be a useful proxy to assess ecosystem resilience, since deep metagenomic sequencing across many samples is economically prohibitive for many studies. (Wemheuer et al., 2020, Sun et al., 2021, Teichert et al., 2018). Therefore, this functional trait-based approach should be considered as a potential tool for assessing the impacts of aquaculture. That said, metagenome prediction tools can lead to incorrect predictions as a result of gene transfer rate or phylogenetic variability. Furthermore, the performance of these tools for inference on genes associated with “housekeeping” drops significantly in environmental samples (Sun et al., 2020, Langille et al., 2013). Therefore, it is important to highlight that the improvement of currently available reference databases used by these inference tools is crucial for environmental impact assessment, since most of the genomes available are associated to human health and biotechnology use (Sun et al., 2020).

### 4.4 Implications for environmental variability

It is important to note a greater number of differentially abundant ASVs associated with the Lease 1 local area, particularly in the near-bottom waters, compared to Lease 2 and control locations. The difference could be related to the production (e.g., stock density, faeces), management practices (e.g., feed input rate), hydrodynamics and location (e.g., shallow or deeper waters) of that specific lease (Belias et al., 2003, Sarà et al., 2007, Tsagaraki et al., 2011, Jansen et al., 2016, Sanderson et al., 2008). Hydrodynamics and location of both leases could be important factors explaining this difference. While Lease 1 is located near the southern end of Macquarie Harbour, Lease 2 is further north, near the deepest basin with a depth of approximately 52 m. Macquarie Harbour is a fjord-like system with periods of deep-water renewal entering by a narrow and shallow entrance located at the Harbour’s northern end. Due to its geological formation, and freshwater, seasonal or weather-related influences, the distribution of the marine water intrusions are heterogeneously distributed leading to a less frequent and less pronounced deep water renewal events in the southern Harbour (Hartstein et al., 2019, Ross et al., 2021). Also, Teasdale et al. (2003) revealed increasing organic carbon and sulphide concentrations from north to south which was influenced by riverine inputs. This north-south environmental gradient may affect the oxygen and organic matter concentration in the water column in the Lease 1 area, causing changes in the environmental conditions as we observed in our study. Although our findings suggest bacterioplankton as a potential indicator for aquaculture activities, more work is needed to confirm its reliability over time in the context of natural variation. Understanding seasonal change in the bacterial community across the Harbour will be particularly important to improve our capacity to detect disturbances caused by both natural and anthropogenic environmental changes (Allison and Martiny, 2008, Magurran et al., 2010, Lindh et al., 2015).

## 5. Conclusions

In summary, investigation of bacterioplankton communities and predicted functions in water column revealed localised changes in the presence of salmon cages. Local enriched taxa were related to biochemical processes (i.e., sulphur, nitrogen, methane, and carbon cycling), potential pathogens or members of the fish gut microbiome, while the relative abundance of function annotations was associated with organic compounds and vitamin degradation, fermentation, methanogenesis and antibiotic resistance. Our study demonstrates that molecular-based methods could be a valuable tool in monitoring aquaculture effects. Trends in nutrient concentrations associated with aquaculture were likely be masked by the natural environmental variation in the water column, strongly influencing the distribution of the bacterial taxonomic and functional communities. The robustness of the aquaculture-associated indicators suggested by this study could be further validated through sampling across both season and farm production cycle to examine temporal variation in taxonomic and functional bacterioplankton communities. Overall, the study showed that the 16S rRNA sequencing and functional annotation of the bacterioplankton communities have the potential to become effective tools in understanding the interaction between finfish aquaculture and the pelagic environment.

## 6. Author contributions

**RRPS, CW and JR**: Conceptualization. **RRPS**: DNA extraction, Formal analysis, Writing-Original draft preparation. **CW, JR and JB**: Supervision, Writing-Reviewing and Editing.

## 7. Declaration of competing interest

We declare that the submitted work was conducted in the absence of any commercial or financial relationships that could potentially be construed as a conflict of interest.

## 8. Acknowledgments

This study was supported by the Fisheries Research & Development Council grant 2016-067 and the Sustainable Marine Research Collaborative Agreement between the Institute for Marine and Antarctic Studies and the State Government of Tasmania. The authors would like to thank Samuel Kruimink, Patricia Peinado Fuentes, Ruth Eriksen, Andy Revill, Kirsten Karsh, Lev Bodrossy and Eric Raes for their assistance in obtaining field samples and laboratory data. We appreciate the help and comments of the anonymous reviewers, that improved our manuscript notably.

## Notes

### Competing Interest Statement

The authors have declared no competing interest.

## References

áNorði, G., Glud, R. N., Gaard, E. and Simonsen, K. (2011) ‘Environmental impacts of coastal fish farming: carbon and nitrogen budgets for trout farming in Kaldbaksfjør7%%FONT_ERR%%ur (Faroe Islands)’, Marine Ecology Progress Series, 431, pp. 223–241. doi: 10.3354/meps09113

Aalto, S. L., Suurnakki, S., von Ahnen, M., Tiirola, M. and Pedersen, P. B. (2021) ‘Microbial communities in full-scale woodchip bioreactors treating aquaculture effluents’, J Environ Manage, 301, pp. 113852. doi: 10.1016/j.jenvman.2021.113852

Aitchison, J. (1986) The Statistical Analysis of Compositional Data. London; New York: Chapman and Hall.

Algammal, A. M., Mabrok, M., Sivaramasamy, E., Youssef, F. M., Atwa, M. H., El-Kholy, A. W., Hetta, H. F. and Hozzein, W. N. (2020) ‘Emerging MDR-Pseudomonas aeruginosa in fish commonly harbor oprL and toxA virulence genes and blaTEM, blaCTX-M, and tetA antibiotic-resistance genes’, Sci Rep, 10(1), pp. 15961. doi: 10.1038/s41598-020-72264-4

Allison, S. D. and Martiny, J. B. (2008) ‘Resistance, resilience, and redundancy in microbial communities’, Proceedings of the National Academy of Sciences, 105(Supplement 1), pp. 11512–11519. doi:

Amirkolaie, A. K. (2011) ‘Reduction in the environmental impact of waste discharged by fish farms through feed and feeding’, Reviews in Aquaculture, 3(1), pp. 19–26. doi: 10.1111/j.1753-5131.2010.01040.x

Appleyard, S., Abell, G. and Watson, R. (2013) Tackling microbial related issues in cultured shellfish via integrated molecular and water chemistry approaches. Seafood CRC Final Report (2011/729) (pp. 89). Available at: http://frdc.com.au/Archived-Reports/FRDC%20Projects/2011-729-DLD.pdf (Accessed: 20 August 2019).

Aranda, C. P., Valenzuela, C., Matamala, Y., Godoy, F. A. and Aranda, N. (2015) ‘Sulphur-cycling bacteria and ciliated protozoans in a Beggiatoaceae mat covering organically enriched sediments beneath a salmon farm in a southern Chilean fjord’, Mar Pollut Bull, 100(1), pp. 270–278. doi: 10.1016/j.marpolbul.2015.08.040

Asami, H., Aida, M. and Watanabe, K. (2005) ‘Accelerated sulfur cycle in coastal marine sediment beneath areas of intensive shellfish aquaculture’, Appl Environ Microbiol, 71(6), pp. 2925–33. doi: 10.1128/AEM.71.6.2925-2933.2005

Asshauer, K. P., Wemheuer, B., Daniel, R. and Meinicke, P. (2015) ‘Tax4Fun: predicting functional profiles from metagenomic 16S rRNA data’, Bioinformatics, 31(17), pp. 2882–4. doi: 10.1093/bioinformatics/btv287

Aylagas, E., Borja, A., Tangherlini, M., Dell’Anno, A., Corinaldesi, C., Michell, C. T., Irigoien, X., Danovaro, R. and Rodriguez-Ezpeleta, N. (2017) ‘A bacterial community-based index to assess the ecological status of estuarine and coastal environments’, Mar Pollut Bull, 114(2), pp. 679–688. doi: 10.1016/j.marpolbul.2016.10.050

Azam, F., Fenchel, T., Field, J., Grey, J., Meyer-Reil, L. and Thingstad, F. (1983) ‘The ecological role of water-column microbes’, Mar. ecol. Prog. ser, 10, pp. 257–263. doi:

Bartl, I., Liskow, I., Schulz, K., Umlauf, L. and Voss, M. (2018) ‘River plume and bottom boundary layer– Hotspots for nitrification in a coastal bay?’, Estuarine, Coastal and Shelf Science, 208, pp. 70–82. doi:

Barton, L. L. and Hamilton, W. A. (2007) Sulphate-reducing bacteria: environmental and engineered systems. Cambridge University Press.

Belias, C. V., Bikas, V. G., Dassenakis, M. J. and Scoullos, M. J. (2003) ‘Environmental impacts of coastal aquaculture in eastern mediterranean bays the case of astakos gulf, Greece’, Environmental Science and Pollution Research, 10(5), pp. 287. doi: 10.1065/espr2003.06.159

Bissett, A., Bowman, J. and Burke, C. (2006) ‘Bacterial diversity in organically-enriched fish farm sediments’, FEMS Microbiol Ecol, 55(1), pp. 48–56. doi: 10.1111/j.1574-6941.2005.00012.x

Bissett, A., Burke, C., Cook, P. L. and Bowman, J. P. (2007) ‘Bacterial community shifts in organically perturbed sediments’, Environ Microbiol, 9(1), pp. 46–60. doi: 10.1111/j.1462-2920.2006.01110.x

Blancheton, J. P., Attramadal, K. J. K., Michaud, L., d’Orbcastel, E. R. and Vadstein, O. (2013) ‘Insight into bacterial population in aquaculture systems and its implication’, Aquacultural Engineering, 53, pp. 30–39. doi: 10.1016/j.aquaeng.2012.11.009

Borja, Á., Rodríguez, J. G., Black, K., Bodoy, A., Emblow, C., Fernandes, T. F., Forte, J., Karakassis, I., Muxika, I., Nickell, T. D., Papageorgiou, N., Pranovi, F., Sevastou, K., Tomassetti, P. and Angel, D. (2009) ‘Assessing the suitability of a range of benthic indices in the evaluation of environmental impact of fin and shellfish aquaculture located in sites across Europe’, Aquaculture, 293(3-4), pp. 231–240. doi: 10.1016/j.aquaculture.2009.04.037

Bouwman, L., Beusen, A., Glibert, P. M., Overbeek, C., Pawlowski, M., Herrera, J., Mulsow, S., Yu, R. and Zhou, M. (2013) ‘Mariculture: significant and expanding cause of coastal nutrient enrichment’, Environmental Research Letters, 8(4). doi: 10.1088/1748-9326/8/4/044026

Bowman, J. S. and Ducklow, H. W. (2015) ‘Microbial Communities Can Be Described by Metabolic Structure: A General Framework and Application to a Seasonally Variable, Depth-Stratified Microbial Community from the Coastal West Antarctic Peninsula’, PLoS One, 10(8), pp. e0135868. doi: 10.1371/journal.pone.0135868

Bozzi, D., Rasmussen, J. A., Carøe, C., Sveier, H., Nordøy, K., Gilbert, M. T. P. and Limborg, M. T. (2021) ‘Salmon gut microbiota correlates with disease infection status: potential for monitoring health in farmed animals’, Animal microbiome, 3(1), pp. 1–17. doi:

Burridge, L., Weis, J. S., Cabello, F., Pizarro, J. and Bostick, K. (2010) ‘Chemical use in salmon aquaculture: A review of current practices and possible environmental effects’, Aquaculture, 306(1), pp. 7–23. doi: https://doi.org/10.1016/j.aquaculture.2010.05.020

Buschmann, A., Riquelme, V., Hernandezgonzalez, M., Varela, D., Jimenez, J., Henriquez, L., Vergara, P., Guinez, R. and Filun, L. (2006) ‘A review of the impacts of salmonid farming on marine coastal ecosystems in the southeast Pacific’, ICES Journal of Marine Science, 63(7), pp. 1338–1345. doi: 10.1016/j.icesjms.2006.04.021

Callahan, B. J., McMurdie, P. J., Rosen, M. J., Han, A. W., Johnson, A. J. and Holmes, S. P. (2016) ‘DADA2: High-resolution sample inference from Illumina amplicon data’, Nat Methods, 13(7), pp. 581–3. doi: 10.1038/nmeth.3869

Cao, Q., Sun, X., Rajesh, K., Chalasani, N., Gelow, K., Katz, B., Shah, V. H., Sanyal, A. J. and Smirnova, E. (2021) ‘Effects of Rare Microbiome Taxa Filtering on Statistical Analysis’, Frontiers in Microbiology, 11. doi: 10.3389/fmicb.2020.607325

Caporaso, J. G., Lauber, C. L., Walters, W. A., Berg-Lyons, D., Lozupone, C. A., Turnbaugh, P. J., Fierer, N. and Knight, R. (2011) ‘Global patterns of 16S rRNA diversity at a depth of millions of sequences per sample’, Proc Natl Acad Sci U S A, 108 Suppl 1, pp. 4516–22. doi: 10.1073/pnas.1000080107

Carpenter, P. D., Butler, E. C. V., Higgins, H. W., Mackey, D. J. and Nichols, P. D. (1991) ‘Chemistry of trace elements, humic substances and sedimentary organic matter in Macquarie Harbour, Tasmania’, Marine and Freshwater Research, 42(6). doi: 10.1071/mf9910625

Caspi, R., Billington, R., Fulcher, C. A., Keseler, I. M., Kothari, A., Krummenacker, M., Latendresse, M., Midford, P. E., Ong, Q. and Ong, W. K. (2018) ‘The MetaCyc database of metabolic pathways and enzymes’, Nucleic acids research, 46(D1), pp. D633–D639. doi:

Chen, C., Zheng, L., Zhou, J. and Zhao, H. (2017) ‘Persistence and risk of antibiotic residues and antibiotic resistance genes in major mariculture sites in Southeast China’, Science of the Total Environment, 580, pp. 1175–1184. doi:

Chen, H., Athar, R., Zheng, G. and Williams, H. N. (2011) ‘Prey bacteria shape the community structure of their predators’, ISME J, 5(8), pp. 1314–22. doi: 10.1038/ismej.2011.4

Cordier, T. (2020) ‘Bacterial communities’ taxonomic and functional turnovers both accurately predict marine benthic ecological quality status’, Environmental DNA, 2(2), pp. 175–183. doi: 10.1002/edn3.55

Cordier, T., Forster, D., Dufresne, Y., Martins, C. I. M., Stoeck, T. and Pawlowski, J. (2018) ‘Supervised machine learning outperforms taxonomy-based environmental DNA metabarcoding applied to biomonitoring’, Molecular Ecology Resources, 18(6), pp. 1381–1391. doi: 10.1111/1755-0998.12926

Cordier, T., Frontalini, F., Cermakova, K., Apotheloz-Perret-Gentil, L., Treglia, M., Scantamburlo, E., Bonamin, V. and Pawlowski, J. (2019) ‘Multi-marker eDNA metabarcoding survey to assess the environmental impact of three offshore gas platforms in the North Adriatic Sea (Italy)’, Mar Environ Res. doi: 10.1016/j.marenvres.2018.12.009

Crawford, C. M., Macleod, C. K. A. and Mitchell, I. M. (2003) ‘Effects of shellfish farming on the benthic environment’, Aquaculture, 224(1-4), pp. 117–140. doi: 10.1016/s0044-8486(03)00210-2

Cresswell, G., Edwards, R. and Barker, B. ‘Macquarie Harbour, Tasmania-seasonal oceanographic surveys in 1985’. Papers and Proceedings of the Royal Society of Tasmania, 63–66.

Da Silva, R. R. P., White, C. A., Bowman, J. P., Raes, E., Bisset, A., Chapman, C., Bodrossy, L. and Ross, D. J. (2021) ‘Environmental influences shaping microbial communities in a low oxygen, highly stratified marine embayment’, Aquatic Microbial Ecology, 87, pp. 185–203. doi: 10.3354/ame01978

DeLong, E. F., Franks, D. G. and Alldredge, A. L. (1993) ‘Phylogenetic diversity of aggregateÁattached vs. freeÁliving marine bacterial assemblages’, Limnology and oceanography, 38(5), pp. 924–934. doi:

Douglas, G. M., Maffei, V. J., Zaneveld, J. R., Yurgel, S. N., Brown, J. R., Taylor, C. M., Huttenhower, C. and Langille, M. G. (2020) ‘PICRUSt2 for prediction of metagenome functions’, Nature biotechnology, 38(6), pp. 685–688. doi:

Dowle, E., Pochon, X., Keeley, N. and Wood, S. A. (2015) ‘Assessing the effects of salmon farming seabed enrichment using bacterial community diversity and high-throughput sequencing’, FEMS microbiology ecology, 91(8), pp. fiv089. doi. Available at: https://watermark.silverchair.com/fiv089.pdf?token=AQECAHi208BE49Ooan9kkhW_Ercy7Dm3ZL_9Cf3qfKAc485ysgAAAkYwggJCBgkqhkiG9w0BBwagggIzMIICLwIBADCCAigGCSqGSIb3DQEHATAeBglghkgBZQMEAS4wEQQMQWvb8ISYQTRAZ8wwAgEQgIIB-fmA6b-VKmxRIKm_00Vh6ojIDonSb5oQGpfAQhoAK7igi1-yzZ-2H5wf9AwmnCPN9mmtNwSHu1U_sKBJ9DUlyT_9xnAYifzcC4yMxRzWZgbKMhZ8s4RrZFNl496VoXcuokjiJPTNitieX4ec1Ek9-BwSn-pHuX2QmqAW1LHrtagPHJeWv55y3148eIz_fZTo9J2wpimBscNkCnSi0r1irXrpljF2GlBgJmPFpShLs3b3kg6y3nXICUHXucM-1TxqhYkasn0KE4-3M8q0tewG8pM823-9bjLCkff61kwjFjR0GHs15jfmg6cGq85Y-3suRr9Q_ec0GGZtVzTyOP-yx9w_49Yj-N147JH4X72e0EilBqgzws1h6KdSTv1T8BGWECVCZSKGy7dZK5RD_v1kDwLJZprNL-hlNQbmPUDomKoSxRXaIZV_Rb5qP6scjFz_K1y2fjove3wKvuO_sAJ5NTLu8Yvx3PtvY5fsaYVPgranqdas5IacMVyOPydOSb48ZMbNXzXjkDvDboUi2SfJ5NljVB1IoAO64B9LRlKuJ-tlecpjpcd74CpzZAsUKZyCi3SlpZC2MqKa7tqkFSG1jK9K_B_hbmVrhOFrKWZKY06E_eEtfjlgiHAwegrbj9Z4y4bIWGdoaXm0BK9kpJHeJLOHwrHN3gvu_3Y

Dully, V., Balliet, H., Frühe, L., Däumer, M., Thielen, A., Gallie, S., Berrill, I. and Stoeck, T. (2021) ‘Robustness, sensitivity and reproducibility of eDNA metabarcoding as an environmental biomonitoring tool in coastal salmon aquaculture – An inter-laboratory study’, Ecological Indicators, 121. doi: 10.1016/j.ecolind.2020.107049

Elizondo-Patrone, C., Hernández, K., Yannicelli, B., Olsen, L. M. and Molina, V. (2015) ‘The response of nitrifying microbial assemblages to ammonium (NH4+) enrichment from salmon farm activities in a northern Chilean Fjord’, Estuarine, Coastal and Shelf Science, 166, pp. 131–142. doi: 10.1016/j.ecss.2015.03.021

Fan, L., Qiu, L., Song, C., Meng, S., Zheng, Y., Li, D., Hu, G. and Chen, J. (2019) ‘Effects of feed input and planting of submerged aquatic vegetation on methanotrophic communities in the surface sediments of aquaculture ponds’, Applied soil ecology, 143, pp. 10–16. doi:

Fernandes, A. D., Macklaim, J. M., Linn, T. G., Reid, G. and Gloor, G. B. (2013) ‘ANOVA-like differential expression (ALDEx) analysis for mixed population RNA-Seq’, PLoS One, 8(7), pp. e67019. doi: 10.1371/journal.pone.0067019

Fodelianakis, S., Papageorgiou, N., Karakassis, I. and Ladoukakis, E. D. (2014) ‘Community structure changes in sediment bacterial communities along an organic enrichment gradient associated with fish farming’, Annals of Microbiology, 65(1), pp. 331–338. doi: 10.1007/s13213-014-0865-4

Fogarty, C., Burgess, C. M., Cotter, P. D., Cabrera-Rubio, R., Whyte, P., Smyth, C. and Bolton, D. J. (2019) ‘Diversity and composition of the gut microbiota of Atlantic salmon (Salmo salar) farmed in Irish waters’, J Appl Microbiol, 127(3), pp. 648–657. doi: 10.1111/jam.14291

Frühe, L., Dully, V., Forster, D., Keeley, N. B., Laroche, O., Pochon, X., Robinson, S., Wilding, T. and Stoeck, T. (2021) ‘Global trends of benthic bacterial diversity and community composition along organic enrichment gradients of salmon farms’, Frontiers in Microbiology, 12, pp. 853. doi:

Fruhe, L., Dully, V., Forster, D., Keeley, N. B., Laroche, O., Pochon, X., Robinson, S., Wilding, T. A. and Stoeck, T. (2021) ‘Global Trends of Benthic Bacterial Diversity and Community Composition Along Organic Enrichment Gradients of Salmon Farms’, Front Microbiol, 12, pp. 637811. doi: 10.3389/fmicb.2021.637811

Fuhrman, J. A. and Steele, J. A. (2008) ‘Community structure of marine bacterioplankton: patterns, networks, and relationships to function’, Aquatic Microbial Ecology, 53, pp. 69–81. doi: 10.3354/ame01222

Galand, P. E., Lucas, S., Fagervold, S. K., Peru, E., Pruski, A. M., Vetion, G., Dupuy, C. and Guizien, K. (2016) ‘Disturbance Increases Microbial Community Diversity and Production in Marine Sediments’, Front Microbiol, 7, pp. 1950. doi: 10.3389/fmicb.2016.01950

Gilbert, J. A., Field, D., Swift, P., Thomas, S., Cummings, D., Temperton, B., Weynberg, K., Huse, S., Hughes, M. and Joint, I. (2010) ‘The taxonomic and functional diversity of microbes at a temperate coastal site: a ‘multi-omic’study of seasonal and diel temporal variation’, PloS one, 5(11), pp. e15545. doi. Available at: https://www.ncbi.nlm.nih.gov/pmc/articles/PMC2993967/pdf/pone.0015545.pdf

Girvan, M., Campbell, C., Killham, K., Prosser, J. I. and Glover, L. A. (2005) ‘Bacterial diversity promotes community stability and functional resilience after perturbation’, Environmental microbiology, 7(3), pp. 301–313. doi:

Gloor, G. (2016) ‘CoDaSeq: Analyzing HTS using compositional data analysis’, F1000Research, 5. doi:

Gloor, G. B., Macklaim, J. M., Pawlowsky-Glahn, V. and Egozcue, J. J. (2017) ‘Microbiome Datasets Are Compositional: And This Is Not Optional’, Front Microbiol, 8, pp. 2224. doi: 10.3389/fmicb.2017.02224

Grasshoff, K., Kremling, K. and Ehrhardt, M. (2009) Methods of seawater analysis. John Wiley & Sons.

Greenwell, B. M. and Boehmke, B. C. (2020) ‘Variable Importance Plots—An Introduction to the vip Package. ‘, The R Journal, 12(1), pp. 343--366. doi: https://doi.org/10.32614/RJ-2020-013.

Hamilton, N. E. and Ferry, M. (2018) ‘ggtern: Ternary diagrams using ggplot2’, Journal of Statistical Software, 87(1), pp. 1–17. doi:

Han, P., Klumper, U., Wong, A., Li, M., Lin, J. G., Quan, Z., Denecke, M. and Gu, J. D. (2017) ‘Assessment of molecular detection of anaerobic ammonium-oxidizing (anammox) bacteria in different environmental samples using PCR primers based on 16S rRNA and functional genes’, Appl Microbiol Biotechnol, 101(20), pp. 7689–7702. doi: 10.1007/s00253-017-8502-3

Haro-Moreno, J. M., Coutinho, F. H., Zaragoza-Solas, A., Picazo, A., Almagro-Moreno, S. and Lopez-Perez, M. (2020) ‘Dysbiosis in marine aquaculture revealed through microbiome analysis: reverse ecology for environmental sustainability’, FEMS Microbiol Ecol, 96(12). doi: 10.1093/femsec/fiaa218

Hartstein, N. D., Maxey, J. D., Loo, J. C. H. and Then, A. Y.-H. (2019) ‘Drivers of deep water renewal in Macquarie Harbour, Tasmania’, Journal of Marine Systems, 199(103226). doi: 10.1016/j.jmarsys.2019.103226

Hawinkel, S., Mattiello, F., Bijnens, L. and Thas, O. (2019) ‘A broken promise: microbiome differential abundance methods do not control the false discovery rate’, Brief Bioinform, 20(1), pp. 210–221. doi: 10.1093/bib/bbx104

Hernandez, M. E., Beck, D. A., Lidstrom, M. E. and Chistoserdova, L. (2015) ‘Oxygen availability is a major factor in determining the composition of microbial communities involved in methane oxidation’, PeerJ, 3, pp. e801. doi:

Ho, H., Lipman, L. and Gaastra, W. (2008) ‘Is Arcobacter a Food Related Pathogen Causing an Emerging Disease?’, International Journal of Infectious Diseases, 12, pp. e325–e326. doi:

Hornick, K. M. and Buschmann, A. H. (2017) ‘Insights into the diversity and metabolic function of bacterial communities in sediments from Chilean salmon aquaculture sites’, Annals of Microbiology, 68(2), pp. 63–77. doi: 10.1007/s13213-017-1317-8

Hou, L., Xie, X., Wan, X., Kao, S.-J., Jiao, N. and Zhang, Y. (2018) ‘Niche differentiation of ammonia and nitrite oxidizers along a salinity gradient from the Pearl River estuary to the South China Sea’, Biogeosciences, 15(16), pp. 5169–5187. doi: 10.5194/bg-15-5169-2018

Jansen, H. M., Reid, G. K., Bannister, R. J., Husa, V., Robinson, S. M. C., Cooper, J. A., Quinton, C. and Strand, Ø. (2016) ‘Discrete water quality sampling at open-water aquaculture sites: limitations and strategies’, Aquaculture Environment Interactions, 8, pp. 463–480. doi: 10.3354/aei00192

Kandel, P. P., Pasternak, Z., van Rijn, J., Nahum, O. and Jurkevitch, E. (2014) ‘Abundance, diversity and seasonal dynamics of predatory bacteria in aquaculture zero discharge systems’, FEMS Microbiol Ecol, 89(1), pp. 149–61. doi: 10.1111/1574-6941.12342

Author (2020) factoextra: Extract and Visualize the Results of Multivariate Data Analyses. R package version 1.0.7. https://CRAN.R-project.org/package=factoextra.

Keeley, N., Laroche, O., Birch, M. and Pochon, X. (2021) ‘A Substrate-Independent Benthic Sampler (SIBS) for Hard and Mixed-Bottom Marine Habitats: A Proof-of-Concept Study’, Frontiers in Marine Science, 8. doi: 10.3389/fmars.2021.627687

Keeley, N., Wood, S. A. and Pochon, X. (2018) ‘Development and preliminary validation of a multi-trophic metabarcoding biotic index for monitoring benthic organic enrichment’, Ecological Indicators, 85, pp. 1044–1057. doi: 10.1016/j.ecolind.2017.11.014

King, R. D. and Tyler, P. A. (1982) ‘Downstream effects of the Gorden River Power Development, south-west Tasmania’, Marine and Freshwater Research, 33(3), pp. 431–442. doi:

Kirkpatrick, J. B., Kriwoken, L. K. and Styger, J. (2019) ‘The reverse precautionary principle: science, the environment and the salmon aquaculture industry in Macquarie Harbour, Tasmania, Australia’, Pacific Conservation Biology, 25(1). doi: 10.1071/pc17014

Koehnken, L. (2005) ‘Overview of Water Quality in Macquarie Harbour and Assessment of Risks due to Copper Levels’. doi:

Kolda, A., Gavrilović, A., Jug-Dujaković, J., Ljubešić, Z., El-Matbouli, M., Lillehaug, A., Lončarević, S., Perić, L., Knežević, D., Vukić Lušić, D. and Kapetanović, D. (2020) ‘Profiling of bacterial assemblages in the marine cage farm environment, with implications on fish, human and ecosystem health’, Ecological Indicators, 118. doi: 10.1016/j.ecolind.2020.106785

Kopprio, G. A., Cuong, L. H., Luyen, N. D., Duc, T. M., Ha, T. H., Huong, L. M. and Gärdes, A. (2021) ‘Carrageenophyte-attached and planktonic bacterial communities in two distinct bays of Vietnam: Eutrophication indicators and insights on ice-ice disease’, Ecological Indicators, 121. doi: 10.1016/j.ecolind.2020.107067

Author (2020) Tidymodels: a collection of packages for modeling and machine learning using tidyverse principles. https://www.tidymodels.org.

Kwon, M. J., O’Loughlin, E. J., Boyanov, M. I., Brulc, J. M., Johnston, E. R., Kemner, K. M. and Antonopoulos, D. A. (2016) ‘Impact of organic carbon electron donors on microbial community development under iron-and sulfate-reducing conditions’, PloS one, 11(1), pp. e0146689. doi:

Labbate, M., Seymour, J. R., Lauro, F. and Brown, M. V. (2016) ‘Editorial: Anthropogenic Impacts on the Microbial Ecology and Function of Aquatic Environments’, Front Microbiol, 7, pp. 1044. doi: 10.3389/fmicb.2016.01044

Lane, D. (1991) ‘16S/23S rRNA sequencing’, Nucleic acid techniques in bacterial systematics, pp. 115–175. doi:

Lane, D. J., Pace, B., Olsen, G. J., Stahl, D. A., Sogin, M. L. and Pace, N. R. (1985) ‘Rapid determination of 16S ribosomal RNA sequences for phylogenetic analyses’, Proceedings of the National Academy of Sciences, 82(20), pp. 6955–6959. doi:

Langille, M. G., Zaneveld, J., Caporaso, J. G., McDonald, D., Knights, D., Reyes, J. A., Clemente, J. C., Burkepile, D. E., Vega Thurber, R. L., Knight, R., Beiko, R. G. and Huttenhower, C. (2013) ‘Predictive functional profiling of microbial communities using 16S rRNA marker gene sequences’, Nat Biotechnol, 31(9), pp. 814–21. doi: 10.1038/nbt.2676

Laroche, O., Meier, S., Mjøs, S. A. and Keeley, N. (2021) ‘Effects of fish farm activities on the sponge Weberella bursa, and its associated microbiota’, Ecological Indicators, 129. doi: 10.1016/j.ecolind.2021.107879

Laroche, O., Pochon, X., Tremblay, L. A., Ellis, J. I., Lear, G. and Wood, S. A. (2018) ‘Incorporating molecular-based functional and co-occurrence network properties into benthic marine impact assessments’, FEMS Microbiol Ecol, 94(11). doi: 10.1093/femsec/fiy167

Liaw, A. and Wiener, M. (2002) ‘Classification and regression by randomForest’, R news, 2(3), pp. 18–22. doi:

Lindh, M. V., Lefebure, R., Degerman, R., Lundin, D., Andersson, A. and Pinhassi, J. (2015) ‘Consequences of increased terrestrial dissolved organic matter and temperature on bacterioplankton community composition during a Baltic Sea mesocosm experiment’, Ambio, 44 Suppl 3, pp. 402–12. doi: 10.1007/s13280-015-0659-3

Lindh, M. V. and Pinhassi, J. (2018) ‘Sensitivity of Bacterioplankton to Environmental Disturbance: A Review of Baltic Sea Field Studies and Experiments’, Frontiers in Marine Science, 5. doi: 10.3389/fmars.2018.00361

Loch, T., Scribner, K., Tempelman, R., Whelan, G. and Faisal, M. (2012) ‘Bacterial infections of Chinook salmon, Oncorhynchus tshawytscha (Walbaum), returning to gamete collecting weirs in Michigan’, Journal of Fish Diseases, 35(1), pp. 39–50. doi:

Logares, R., Sunagawa, S., Salazar, G., Cornejo-Castillo, F. M., Ferrera, I., Sarmento, H., Hingamp, P., Ogata, H., de Vargas, C., Lima-Mendez, G., Raes, J., Poulain, J., Jaillon, O., Wincker, P., Kandels-Lewis, S., Karsenti, E., Bork, P. and Acinas, S. G. (2014) ‘Metagenomic 16S rDNA Illumina tags are a powerful alternative to amplicon sequencing to explore diversity and structure of microbial communities’, Environ Microbiol, 16(9), pp. 2659–71. doi: 10.1111/1462-2920.12250

Lokesh, J., Kiron, V., Sipkema, D., Fernandes, J. M. and Moum, T. (2019) ‘Succession of embryonic and the intestinal bacterial communities of Atlantic salmon (Salmo salar) reveals stageÁspecific microbial signatures’, MicrobiologyOpen, 8(4), pp. e00672. doi:

Louca, S., Parfrey, L. W. and Doebeli, M. (2016) ‘Decoupling function and taxonomy in the global ocean microbiome’, Science, 353(6305), pp. 1272-1277. doi. Available at: http://science.sciencemag.org/content/353/6305/1272.long

Louca, S., Polz, M. F., Mazel, F., Albright, M. B. N., Huber, J. A., O’Connor, M. I., Ackermann, M., Hahn, A. S., Srivastava, D. S., Crowe, S. A., Doebeli, M. and Parfrey, L. W. (2018) ‘Function and functional redundancy in microbial systems’, Nat Ecol Evol, 2(6), pp. 936–943. doi: 10.1038/s41559-018-0519-1

Love, M. I., Huber, W. and Anders, S. (2014) ‘Moderated estimation of fold change and dispersion for RNA-seq data with DESeq2’, Genome Biol, 15(12), pp. 550. doi: 10.1186/s13059-014-0550-8

Magurran, A. E., Baillie, S. R., Buckland, S. T., Dick, J. M., Elston, D. A., Scott, E. M., Smith, R. I., Somerfield, P. J. and Watt, A. D. (2010) ‘Long-term datasets in biodiversity research and monitoring: assessing change in ecological communities through time’, Trends Ecol Evol, 25(10), pp. 574–82. doi: 10.1016/j.tree.2010.06.016

Mahmoud, M. A. A. and Magdy, M. (2021) ‘Metabarcoding profiling of microbial diversity associated with trout fish farming’, Sci Rep, 11(1), pp. 421. doi: 10.1038/s41598-020-80236-x

Martínez-Porchas, M. and Vargas-Albores, F. (2017) ‘Microbial metagenomics in aquaculture: a potential tool for a deeper insight into the activity’, Reviews in Aquaculture, 9(1), pp. 42–56. doi: 10.1111/raq.12102

Martins, P., Cleary, D. F., Pires, A. C., Rodrigues, A. M., Quintino, V., Calado, R. and Gomes, N. C. (2013) ‘Molecular analysis of bacterial communities and detection of potential pathogens in a recirculating aquaculture system for Scophthalmus maximus and Solea senegalensis’, PLoS One, 8(11), pp. e80847. doi: 10.1371/journal.pone.0080847

Martins, P., Coelho, F. J. R. C., Cleary, D. F. R., Pires, A. C. C., Marques, B., Rodrigues, A. M., Quintino, V. and Gomes, N. C. M. (2018) ‘Seasonal patterns of bacterioplankton composition in a semi-intensive European seabass (Dicentrarchus labrax) aquaculture system’, Aquaculture, 490, pp. 240–250. doi: 10.1016/j.aquaculture.2018.02.038

Martiny, J. B., Bohannan, B. J., Brown, J. H., Colwell, R. K., Fuhrman, J. A., Green, J. L., Horner-Devine, M. C., Kane, M., Krumins, J. A., Kuske, C. R., Morin, P. J., Naeem, S., Ovreas, L., Reysenbach, A. L., Smith, V. H. and Staley, J. T. (2006) ‘Microbial biogeography: putting microorganisms on the map’, Nat Rev Microbiol, 4(2), pp. 102–12. doi: 10.1038/nrmicro1341

McCaig, A. E., Phillips, C. J., Stephen, J. R., Kowalchuk, G. A., Harvey, S. M., Herbert, R. A., Embley, T. M. and Prosser, J. I. (1999) ‘Nitrogen cycling and community structure of proteobacterial β-subgroup ammonia-oxidizing bacteria within polluted marine fish farm sediments’, Applied and Environmental Microbiology, 65(1), pp. 213-220. doi. Available at: https://www.ncbi.nlm.nih.gov/pmc/articles/PMC91005/pdf/am000213.pdf

McMurdie, P. J. and Holmes, S. (2013) ‘phyloseq: an R package for reproducible interactive analysis and graphics of microbiome census data’, PloS one, 8(4), pp. e61217. doi:

Minich, J. J., Poore, G. D., Jantawongsri, K., Johnston, C., Bowie, K., Bowman, J., Knight, R., Nowak, B. and Allen, E. E. (2020) ‘Microbial ecology of Atlantic salmon (Salmo salar) hatcheries: impacts of the built environment on fish mucosal microbiota’, Applied and environmental microbiology, 86(12), pp. e00411–20. doi:

Miranda, C. D. and Zemelman, R. (2002) ‘Bacterial resistance to oxytetracycline in Chilean salmon farming’, Aquaculture, 212(1-4), pp. 31–47. doi:

Moncada, C., Hassenruck, C., Gardes, A. and Conaco, C. (2019) ‘Microbial community composition of sediments influenced by intensive mariculture activity’, FEMS Microbiol Ecol, 95(2). doi: 10.1093/femsec/fiz006

Morelan, I. A. (2019) ‘16S rRNA Gene Amplicon Sequencing Reveals Trends in Marine Bacterial Diversity and Taxonomic Composition in Natural and Human-built Systems’. doi:

Mukherjee, A., Chettri, B., Langpoklakpam, J. S., Basak, P., Prasad, A., Mukherjee, A. K., Bhattacharyya, M., Singh, A. K. and Chattopadhyay, D. (2017) ‘Bioinformatic approaches including predictive metagenomic profiling reveal characteristics of bacterial response to petroleum hydrocarbon contamination in diverse environments’, Scientific reports, 7(1), pp. 1–22. doi:

Nagpal, S., Haque, M. M. and Mande, S. S. (2016) ‘Vikodak--A Modular Framework for Inferring Functional Potential of Microbial Communities from 16S Metagenomic Datasets’, PLoS One, 11(2), pp. e0148347. doi: 10.1371/journal.pone.0148347

Navarro, N., Leakey, R. J. G. and Black, K. D. (2008) ‘Effect of salmon cage aquaculture on the pelagic environment of temperate coastal waters: seasonal changes in nutrients and microbial community’, Marine Ecology Progress Series, 361, pp. 47–58. doi. Available at: https://www.int-res.com/abstracts/meps/v361/p47-58/

Newell, S. E., Fawcett, S. E. and Ward, B. B. (2013) ‘Depth distribution of ammonia oxidation rates and ammonia-oxidizer community composition in the Sargasso Sea’, Limnology and Oceanography, 58(4), pp. 1491–1500. doi: 10.4319/lo.2013.58.4.1491

Oksanen, J. F., Blanchet, G., Friendly, M., Kindt, R., Legendre, P., McGlinn, D., Peter, R. and Minchin, R. (2018) ‘O’Hara; Gavin, L.; et al’, The Vegan Community Ecology Package. Available online: https://cran.r-project.org/web/packages/vegan/(accessed on 1 March 2019). doi:

Olsen, L. M., Hernández, K. L., Ardelan, M. V., Iriarte, J. L., Bizsel, K. C. and Olsen, Y. (2017) ‘Responses in bacterial community structure to waste nutrients from aquaculture: an in situ microcosm experiment in a Chilean fjord’, Aquaculture Environment Interactions, 9, pp. 21–32. doi. Available at: https://www.int-res.com/abstracts/aei/v9/p21-32/

Ortiz-Estrada, Á.M., Gollas-Galván, T., Martínez-Córdova, L. R. and Martínez-Porchas, M. (2019) ‘Predictive functional profiles using metagenomic 16S rRNA data: a novel approach to understanding the microbial ecology of aquaculture systems’, Reviews in Aquaculture, 11(1), pp. 234–245. doi: 10.1111/raq.12237

Pace, N. R. (1997) ‘A Molecular View of Microbial Diversity and the Biosphere’, Science, 276(5313), pp. 734–740. doi: 10.1126/science.276.5313.734

Palarea-Albaladejo, J. and Martín-Fernández, J. A. (2015) ‘zCompositions—R package for multivariate imputation of left-censored data under a compositional approach’, Chemometrics and Intelligent Laboratory Systems, 143, pp. 85–96. doi:

Pineiro, S. A., Stine, O. C., Chauhan, A., Steyert, S. R., Smith, R. and Williams, H. N. (2007) ‘Global survey of diversity among environmental saltwater Bacteriovoracaceae’, Environ Microbiol, 9(10), pp. 2441–50. doi: 10.1111/j.1462-2920.2007.01362.x

Pomeroy, L. R., le B. Williams, P.J., Azam, F. and Hobbie, J. E. (2007) ‘The microbial loop’, Oceanography, 20(2), pp. 28–33. doi:

Qu, J., Yang, H., Liu, Y., Qi, H., Wang, Y. and Zhang, Q. (2021) ‘The study of natural biofilm formation and microbial community structure for recirculating aquaculture system’, IOP Conference Series: Earth and Environmental Science, 742(1). doi: 10.1088/1755-1315/742/1/012018

R Core Team 2021. R: A language and environment for statistical computing. R Foundation for Statistical Computing, Vienna, Austria. URL https://www.R-project.org/.

Raes, E. J., Karsh, K., Sow, S. L. S., Ostrowski, M., Brown, M. V., van de Kamp, J., Franco-Santos, R. M., Bodrossy, L. and Waite, A. M. (2021) ‘Metabolic pathways inferred from a bacterial marker gene illuminate ecological changes across South Pacific frontal boundaries’, Nat Commun, 12(1), pp. 2213. doi: 10.1038/s41467-021-22409-4

Rasigraf, O., Schmitt, J., Jetten, M. S. M. and Luke, C. (2017) ‘Metagenomic potential for and diversity of N-cycle driving microorganisms in the Bothnian Sea sediment’, Microbiologyopen, 6(4). doi: 10.1002/mbo3.475

Reji, L., Tolar, B. B., Chavez, F. P. and Francis, C. A. (2020) ‘Depth-Differentiation and Seasonality of Planktonic Microbial Assemblages in the Monterey Bay Upwelling System’, Front Microbiol, 11, pp. 1075. doi: 10.3389/fmicb.2020.01075

Revill, A. T., Ross, J. a. and Thompson, P. A. (2016) Investigating Dissolved Oxygen Drawdown in Macquarie Harbour. CSIRO, Australia. Report to Huon Aquaculture: Institute for Marine and Antarctic Studies (IMAS), Hobart. Available at: https://www.huonaqua.com.au/wp-content/uploads/2015/08/Appendix-4-part-4.pdf (Accessed: 16 September 2019).

Ross, J., Beard, J., Wild-Allen, K., Andrewartha Moreno, D., Semmens, J., Davey, A., Hortle, J., Pender Quigley B., Andrewartha, J., Stehfest, K., Durand, A. and Macleod, C. a. (2021) Understanding oxygen dynamics and the importance for benthic recovery in Macquarie Harbour. Hobart, Australia CC BY 3.0: Institute for Marine and Antarctic Studies, University of Tasmania.

Rubio-Portillo, E., Villamor, A., Fernandez-Gonzalez, V., Antón, J. and Sanchez-Jerez, P. (2019) ‘Exploring changes in bacterial communities to assess the influence of fish farming on marine sediments’, Aquaculture, 506, pp. 459–464. doi: 10.1016/j.aquaculture.2019.03.051

Sanderson, J. C., Cromey, C. J., Dring, M. J. and Kelly, M. S. (2008) ‘Distribution of nutrients for seaweed cultivation around salmon cages at farm sites in north–west Scotland’, Aquaculture, 278(1-4), pp. 60–68. doi: 10.1016/j.aquaculture.2008.03.027

Sarà, G., Lo Martire, M., Buffa, G., Mannino, A. M. and Badalamenti, F. (2007) ‘The fouling community as an indicator of fish farming impact in Mediterranean’, Aquaculture Research, 38(1), pp. 66–75. doi: 10.1111/j.1365-2109.2006.01632.x

Sevigny, J. L., Rothenheber, D., Diaz, K. S., Zhang, Y., Agustsson, K., Bergeron, R. D. and Thomas, W. K. (2019) ‘Marker genes as predictors of shared genomic function’, BMC Genomics, 20(1), pp. 268. doi: 10.1186/s12864-019-5641-1

Sinnott, R. W. (1984) ‘Virtues of the Haversine’, S&T, 68(2), pp. 158. doi:

Stoeck, T., Fruhe, L., Forster, D., Cordier, T., Martins, C. I. M. and Pawlowski, J. (2018) ‘Environmental DNA metabarcoding of benthic bacterial communities indicates the benthic footprint of salmon aquaculture’, Mar Pollut Bull, 127, pp. 139–149. doi: 10.1016/j.marpolbul.2017.11.065

Sun, S., Jones, R. B. and Fodor, A. A. (2020) ‘Inference-based accuracy of metagenome prediction tools varies across sample types and functional categories’, Microbiome, 8(1), pp. 46. doi: 10.1186/s40168-020-00815-y

Sun, X., Dong, J., Hu, C., Zhang, Y., Chen, Y. and Zhang, X. (2021) ‘Use of macrofaunal assemblage indices and biological trait analysis to assess the ecological impacts of coastal bivalve aquaculture’, Ecological Indicators, 127, pp. 107713. doi:

Tamminen, M., Karkman, A., Corander, J., Paulin, L. and Virta, M. (2011) ‘Differences in bacterial community composition in Baltic Sea sediment in response to fish farming’, Aquaculture, 313(1-4), pp. 15–23. doi: 10.1016/j.aquaculture.2011.01.020

Teasdale, P. R., Apte, S. C., Ford, P. W., Batley, G. E. and Koehnken, L. (2003) ‘Geochemical cycling and speciation of copper in waters and sediments of Macquarie Harbour, Western Tasmania’, Estuarine, Coastal and Shelf Science, 57(3), pp. 475–487. doi: 10.1016/s0272-7714(02)00381-5

Teichert, N., Lepage, M. and Lobry, J. (2018) ‘Beyond classic ecological assessment: the use of functional indices to indicate fish assemblages sensitivity to human disturbance in estuaries’, Science of the Total Environment, 639, pp. 465–475. doi:

Terry, C. (2001) ‘Numerical Modelling of Macquarie Harbour, Tasmania’, In: Conference on Hydraulics in Civil Engineering (6th : 2001 : Hobart, Tas.). 6th Conference on Hydraulics in Civil Engineering: The State of Hydraulics; Proceedings. Barton, A.C.T.: Institution of Engineers, Australia, 2001: 345–354.

Tsagaraki, T. M., Petihakis, G., Tsiaras, K., Triantafyllou, G., Tsapakis, M., Korres, G., Kakagiannis, G., Frangoulis, C. and Karakassis, I. (2011) ‘Beyond the cage: ecosystem modelling for impact evaluation in aquaculture’, Ecological modelling, 222(14), pp. 2512–2523. doi:

Tsukamoto, K., Kawamura, T., Takeuchi, T., Beard Jr, T. and Kaiser, M. (2008) ‘Environmental Impact of Aquaculture on Coastal Planktonic Ecosystems’, Fisheries for Global Welfare and Environment, pp. 181. doi:

Vargas-Albores, F., Martínez-Córdova, L. R., Gollas-Galván, T., Garibay-Valdez, E., Emerenciano, M. G. C., Lago-Leston, A., Mazorra-Manzano, M. and Martínez-Porchas, M. (2019) ‘Inferring the functional properties of bacterial communities in shrimp-culture bioflocs produced with amaranth and wheat seeds as fouler promoters’, Aquaculture, 500, pp. 107–117. doi: 10.1016/j.aquaculture.2018.10.005

Verhoeven, J. T. P., Salvo, F., Hamoutene, D. and Dufour, S. C. (2016) ‘Bacterial community composition of flocculent matter under a salmonid aquaculture site in Newfoundland, Canada’, Aquaculture Environment Interactions, 8, pp. 637–646. doi: 10.3354/aei00204

Verhoeven, J. T. P., Salvo, F., Knight, R., Hamoutene, D. and Dufour, S. C. (2018) ‘Temporal Bacterial Surveillance of Salmon Aquaculture Sites Indicates a Long Lasting Benthic Impact With Minimal Recovery’, Front Microbiol, 9, pp. 3054. doi: 10.3389/fmicb.2018.03054

Wakabayashi, H. (1980) ‘Bacterial gill disease of salmonid fish’, Fish pathology, 14(4), pp. 185–189. doi:

Wang, Q., Garrity, G. M., Tiedje, J. M. and Cole, J. R. (2007) ‘Naive Bayesian classifier for rapid assignment of rRNA sequences into the new bacterial taxonomy’, Appl Environ Microbiol, 73(16), pp. 5261–7. doi: 10.1128/AEM.00062-07

Wang, X., Olsen, L. M., Reitan, K. I. and Olsen, Y. (2012) ‘Discharge of nutrient wastes from salmon farms: environmental effects, and potential for integrated multi-trophic aquaculture’, Aquaculture Environment Interactions, 2(3), pp. 267–283. doi: 10.3354/aei00044

Wei, G., Li, M., Li, F., Li, H. and Gao, Z. (2016) ‘Distinct distribution patterns of prokaryotes between sediment and water in the Yellow River estuary’, Appl Microbiol Biotechnol, 100(22), pp. 9683–9697. doi: 10.1007/s00253-016-7802-3

Wemheuer, F., Taylor, J. A., Daniel, R., Johnston, E., Meinicke, P., Thomas, T. and Wemheuer, B. (2020) ‘Tax4Fun2: prediction of habitat-specific functional profiles and functional redundancy based on 16S rRNA gene sequences’, Environ Microbiome, 15(1), pp. 11. doi: 10.1186/s40793-020-00358-7

Wickham, H., Averick, M., Bryan, J., Chang, W., McGowan, L., François, R., Grolemund, G., Hayes, A., Henry, L., Hester, J., Kuhn, M., Pedersen, T., Miller, E., Bache, S., Müller, K., Ooms, J., Robinson, D., Seidel, D., Spinu, V., Takahashi, K., Vaughan, D., Wilke, C., Woo, K. and Yutani, H. (2019) ‘Welcome to the Tidyverse’, Journal of Open Source Software, 4(43), pp. 1686. doi: 10.21105/joss.01686

Won, N. I., Kim, K. H., Kang, J. H., Park, S. R. and Lee, H. J. (2017) ‘Exploring the Impacts of Anthropogenic Disturbance on Seawater and Sediment Microbial Communities in Korean Coastal Waters Using Metagenomics Analysis’, Int J Environ Res Public Health, 14(2), pp. 130. doi: 10.3390/ijerph14020130

Wu, D. M., Dai, Q. P., Liu, X. Z., Fan, Y. P. and Wang, J. X. (2019) ‘Comparison of bacterial community structure and potential functions in hypoxic and non-hypoxic zones of the Changjiang Estuary’, PLoS One, 14(6), pp. e0217431. doi: 10.1371/journal.pone.0217431

Wu, R. S. S. (1995) ‘The environmental impact of marine fish culture: Towards a sustainable future’, Marine Pollution Bulletin, 31(4), pp. 159–166. doi: https://doi.org/10.1016/0025-326X(95)00100-2

Yang, Y., Gao, Y., Huang, X., Ni, P., Wu, Y., Deng, Y. and Zhan, A. (2019) ‘Adaptive shifts of bacterioplankton communities in response to nitrogen enrichment in a highly polluted river’, Environ Pollut, 245, pp. 290–299. doi: 10.1016/j.envpol.2018.11.002

Yarza, P., Yilmaz, P., Pruesse, E., Glockner, F. O., Ludwig, W., Schleifer, K. H., Whitman, W. B., Euzeby, J., Amann, R. and Rossello-Mora, R. (2014) ‘Uniting the classification of cultured and uncultured bacteria and archaea using 16S rRNA gene sequences’, Nat Rev Microbiol, 12(9), pp. 635–45. doi: 10.1038/nrmicro3330

Yilmaz, P., Parfrey, L. W., Yarza, P., Gerken, J., Pruesse, E., Quast, C., Schweer, T., Peplies, J., Ludwig, W. and Glöckner, F. O. (2014) ‘The SILVA and “all-species living tree project (LTP)” taxonomic frameworks’, Nucleic acids research, 42(D1), pp. D643–D648. doi:

Yoshikawa, T. and Eguchi, M. (2013) ‘Planktonic processes contribute significantly to the organic carbon budget of a coastal fish-culturing area’, Aquaculture Environment Interactions, 4(3), pp. 239–250. doi:

Yu, Z., Yang, J., Amalfitano, S., Yu, X. and Liu, L. (2014) ‘Effects of water stratification and mixing on microbial community structure in a subtropical deep reservoir’, Sci Rep, 4, pp. 5821. doi: 10.1038/srep05821

Zarkasi, K. Z., Taylor, R. S., Abell, G. C., Tamplin, M. L., Glencross, B. D. and Bowman, J. P. (2016) ‘Atlantic salmon (Salmo salar L.) gastrointestinal microbial community dynamics in relation to digesta properties and diet’, Microbial ecology, 71(3), pp. 589–603. doi:

Zhang, K., Zheng, X., He, Z., Yang, T., Shu, L., Xiao, F., Wu, Y., Wang, B., Li, Z., Chen, P. and Yan, Q. (2020) ‘Fish growth enhances microbial sulfur cycling in aquaculture pond sediments’, Microb Biotechnol, 13(5), pp. 1597–1610. doi: 10.1111/1751-7915.13622

Zhang, Q., Tang, F., Zhou, Y., Xu, J., Chen, H., Wang, M. and Laanbroek, H. J. (2015) ‘Shifts in the pelagic ammonia-oxidizing microbial communities along the eutrophic estuary of Yong River in Ningbo City, China’, Front Microbiol, 6, pp. 1180. doi: 10.3389/fmicb.2015.01180

Zorz, J., Willis, C., Comeau, A. M., Langille, M. G. I., Johnson, C. L., Li, W. K. W. and LaRoche, J. (2019) ‘Drivers of Regional Bacterial Community Structure and Diversity in the Northwest Atlantic Ocean’, Front Microbiol, 10, pp. 281. doi: 10.3389/fmicb.2019.00281

