## Supplementary material for "Composition and functionality of bacterioplankton communities in marine coastal zones adjacent to finfish aquaculture": sup_material_figures

### Supplementary Material – Figures

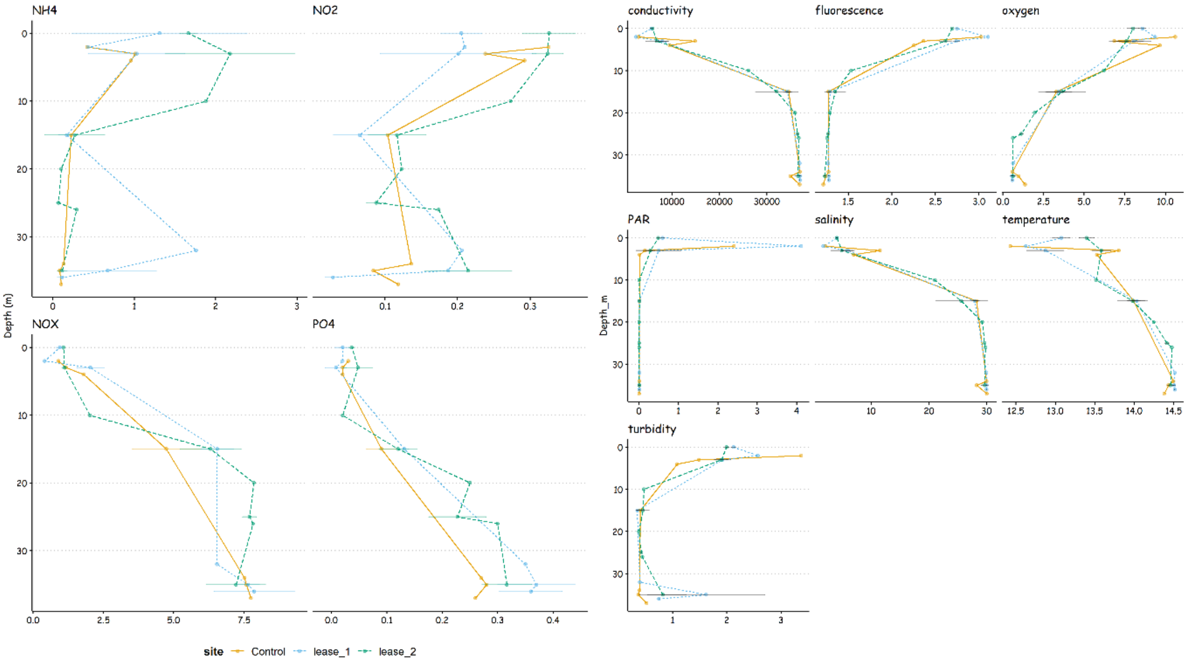


Fig. S1 – Depth profile of the environmental variables. Colors represent samples sites (both leases and control sites).

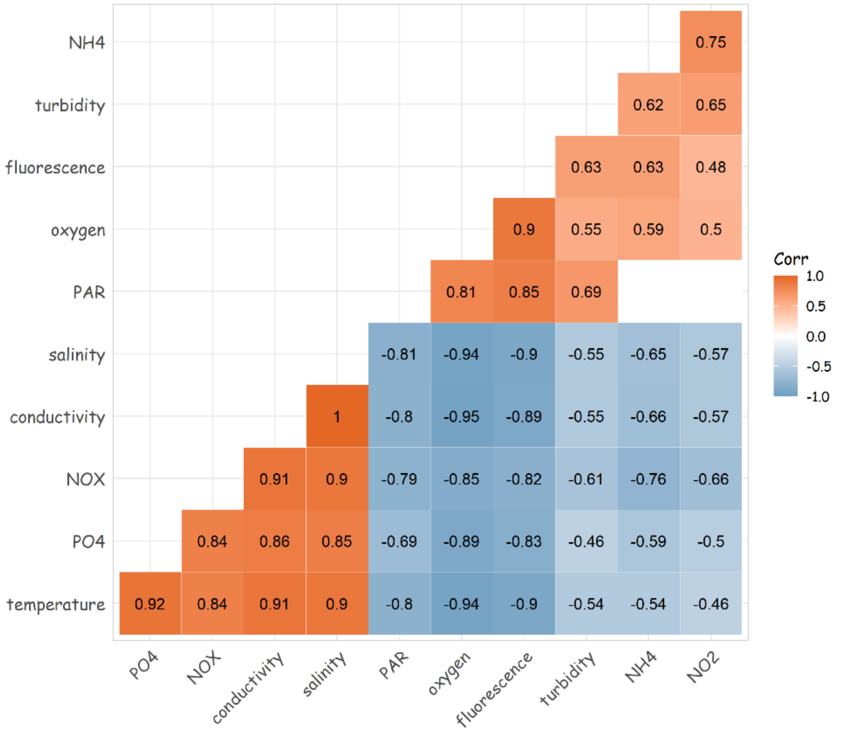


Fig. S2 – Correlation between environmental variables. Spearman rho values with p $\leq$ 0.05 are displayed. Blank squares are no significant coefficients.

Control

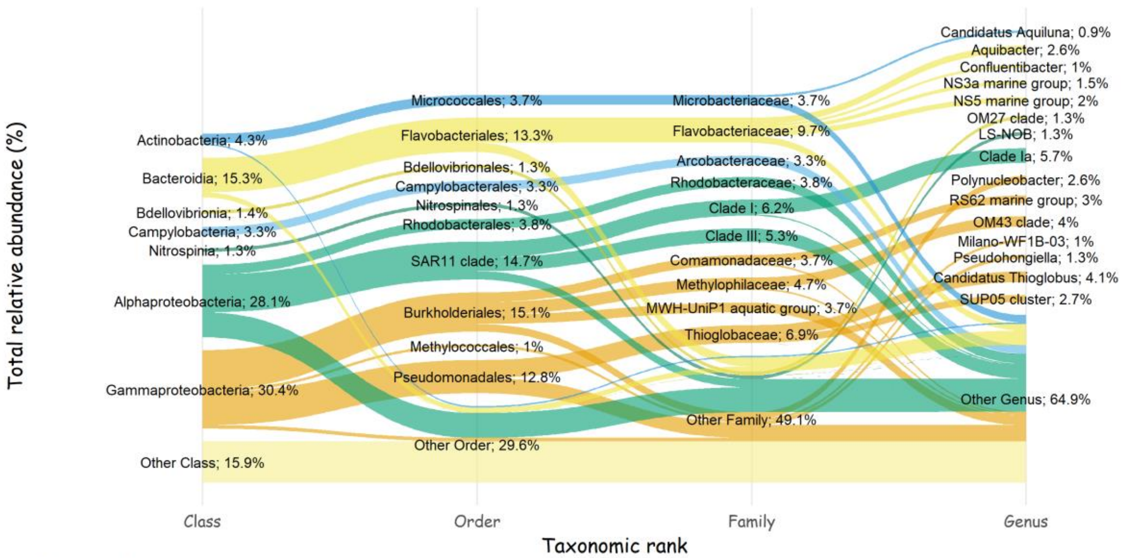


lease_1

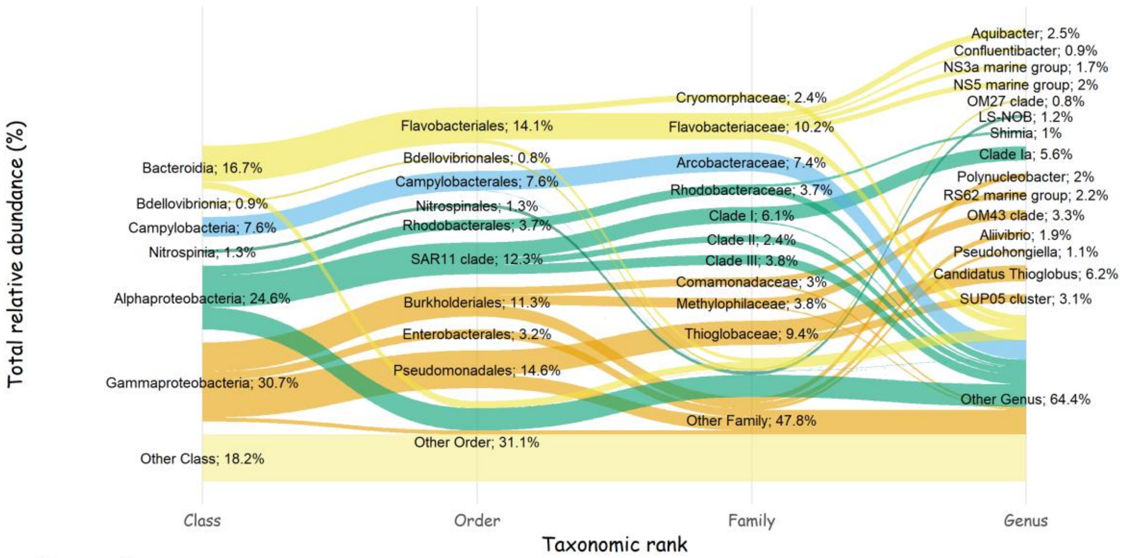


lease_2

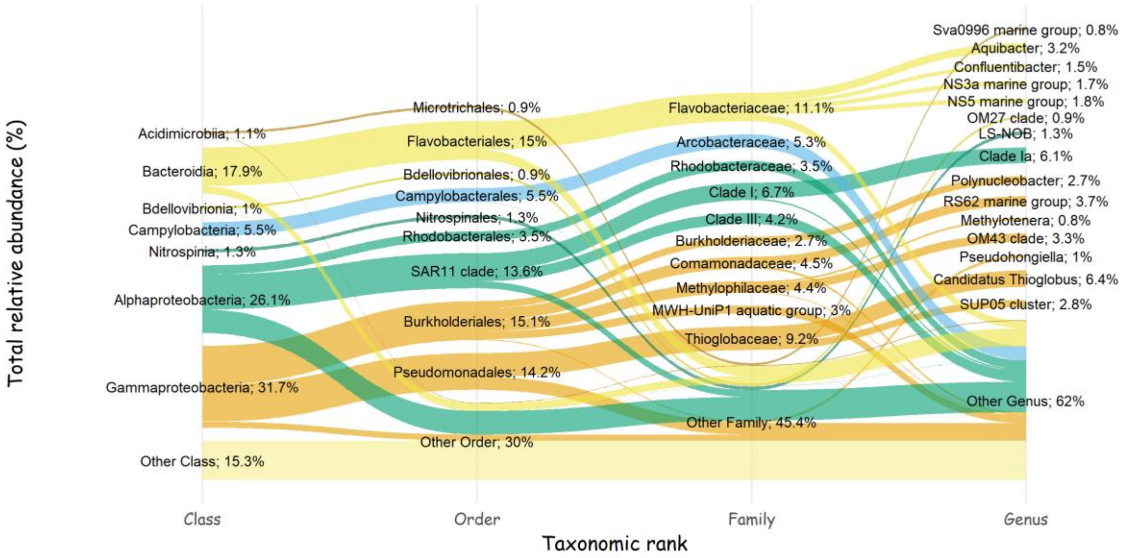

Fig. S3 – Alluvial plot exhibiting the total relative abundance distribution of bacterial community across the leases and control sites. The different taxonomic ranks (class, order, family, and genus) are displayed along with the relative abundance (%).

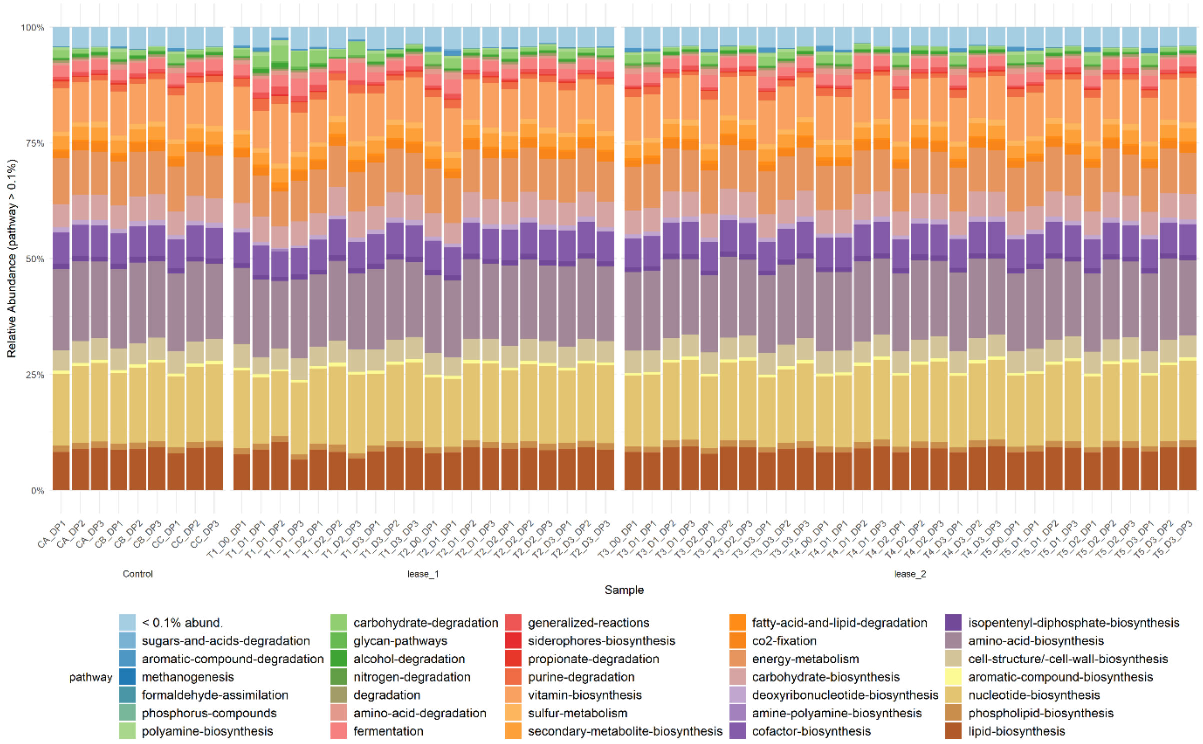


Fig. S4 – Bar plot showing the relative abundance distribution of functional community across sites. Functions with abundance less than 0.1% were agglomerate together.

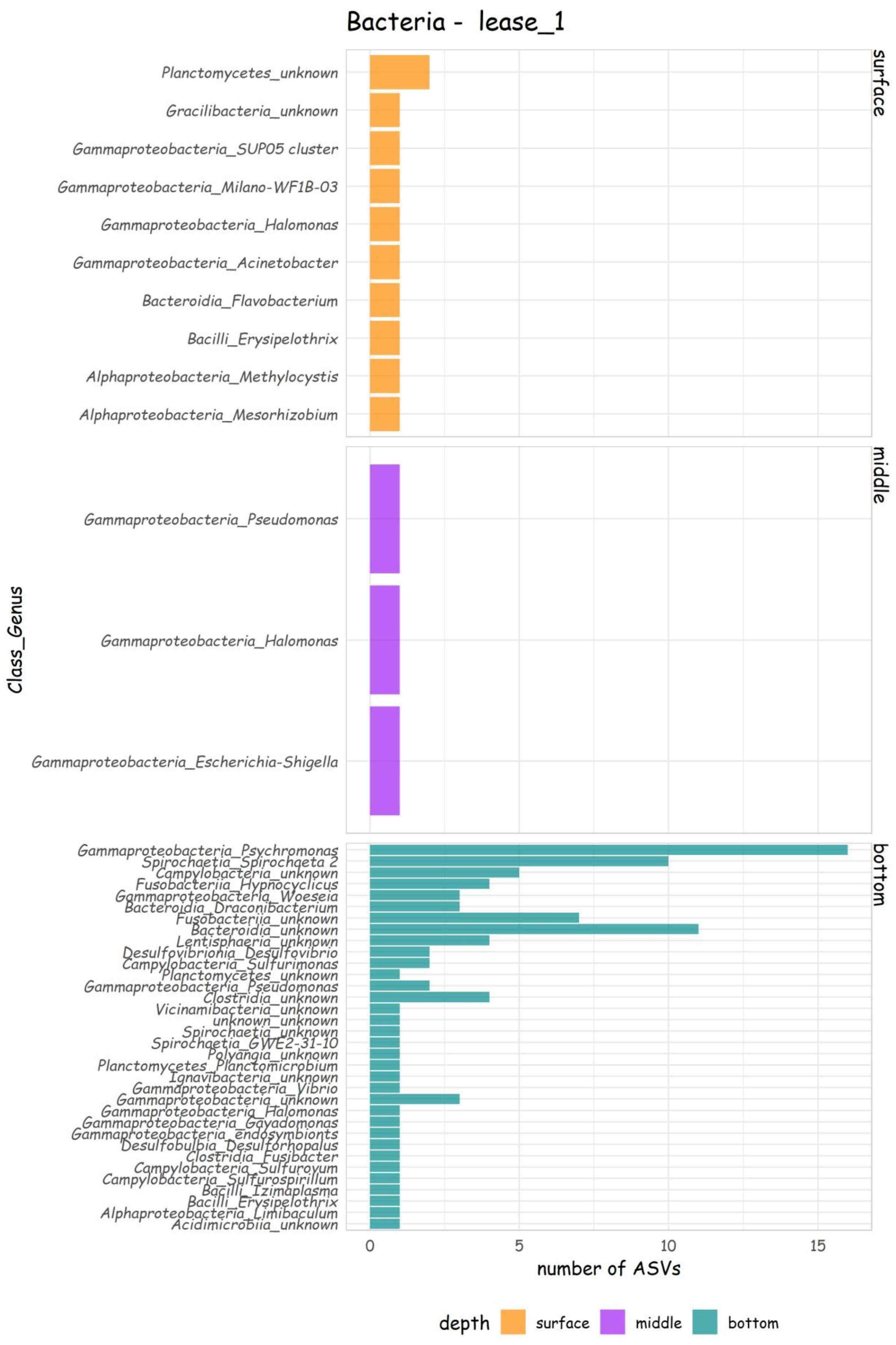
Bacteria - lease_1
surface
middle
bottom
Class_Genus
Planctomycetes_unknown
Gracilibacteria_unknown
Gammaproteobacteria_SUP05 cluster
Gammaproteobacteria_Milano-WF1B-03
Gammaproteobacteria_Halomonas
Gammaproteobacteria_Acinetobacter
Bacteroidia_Flavobacterium
Bacilli_Erysipelothrix
Alphaproteobacteria_Methylocystis
Alphaproteobacteria_Mesorhizobium
Gammaproteobacteria_Pseudomonas
Gammaproteobacteria_Halomonas
Gammaproteobacteria_Escherichia-Shigella
Gammaproteobacteria_Psychromonas
Spirochaetia_Spirochaeta 2
Campylobacteria_unknown
Fusobacteriia_Hypnocyclicus
Gammaproteobacteria_Woeseia
Bacteroidia_Draconibacterium
Fusobacteriia_unknown
Bacteroidia_unknown
Lentisphaeria_unknown
Desulfovibrionia_Desulfovibrio
Campylobacteria_Sulfurimonas
Planctomycetes_unknown
Gammaproteobacteria_Pseudomonas
Clostridia_unknown
Vicinamibacteria_unknown
unknown_unknown
Spirochaetia_unknown
Spirochaetia_GWE2-31-10
Polyangia_unknown
Planctomycetes_Planctomicrobium
Ignavibacteria_unknown
Gammaproteobacteria_Vibrio
Gammaproteobacteria_unknown
Gammaproteobacteria_Halomonas
Gammaproteobacteria_Gayadomonas
Gammaproteobacteria_endosymbionts
Desulfobulbia_Desulforhopalus
Clostridia_Fusibacter
Campylobacteria_Sulfurovum
Campylobacteria_Sulfurospirillum
Bacilli_Izimaplasma
Bacilli_Erysipelothrix
Alphaproteobacteria_Limibaculum
Acidimicrobiia_unknown
0
5
10
15
number of ASVs
depth
surface
middle
bottom

Fig. S5 – Number of ASVs found in ALDEx2 and DESeq2 analyses with relative abundance ≥ 60% in Lease 1.

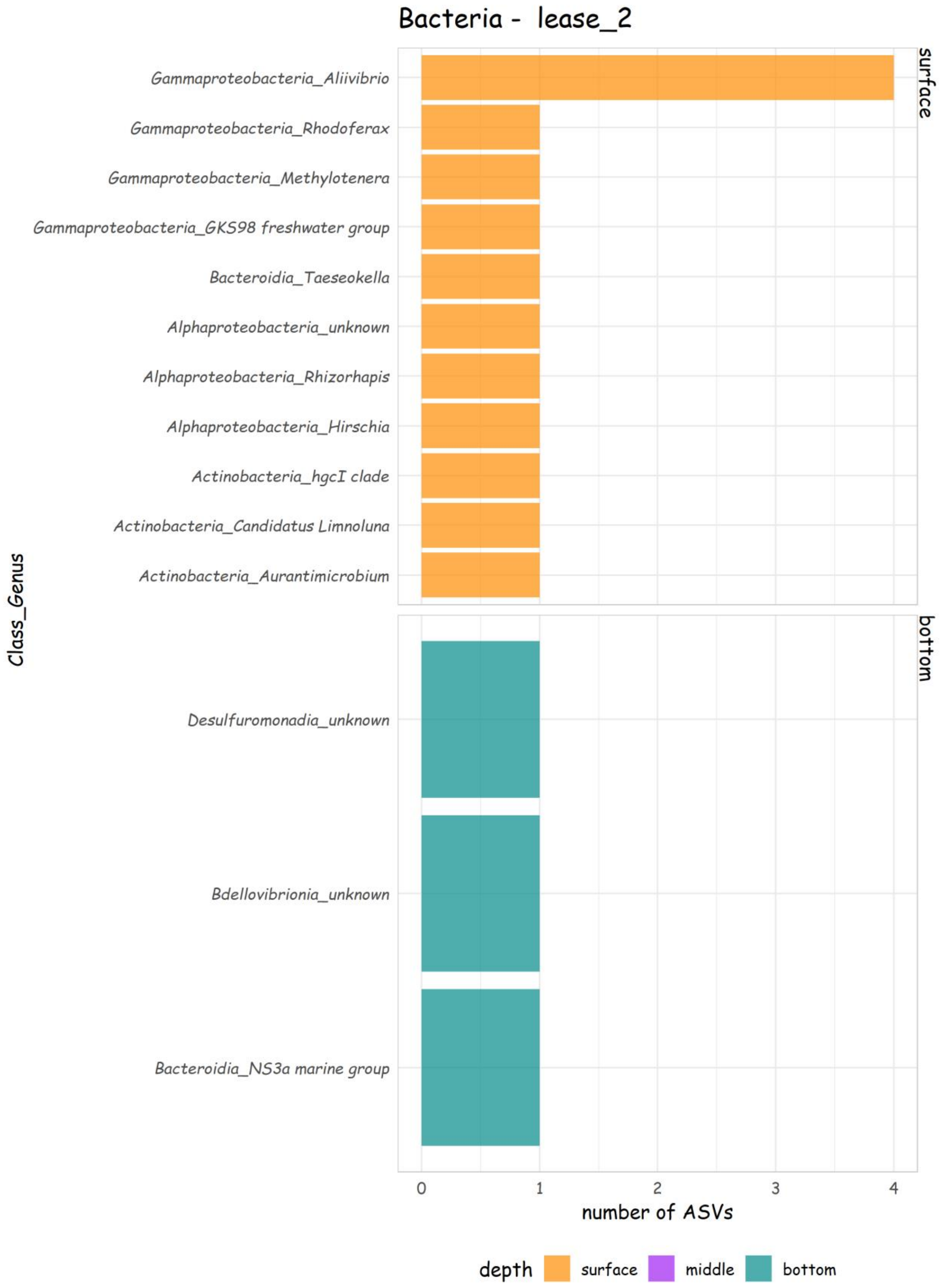


Fig. S6 – Number of ASVs found in ALDEx2 and DESeq2 analyses with relative abundance ≥ 60% in Lease 2.

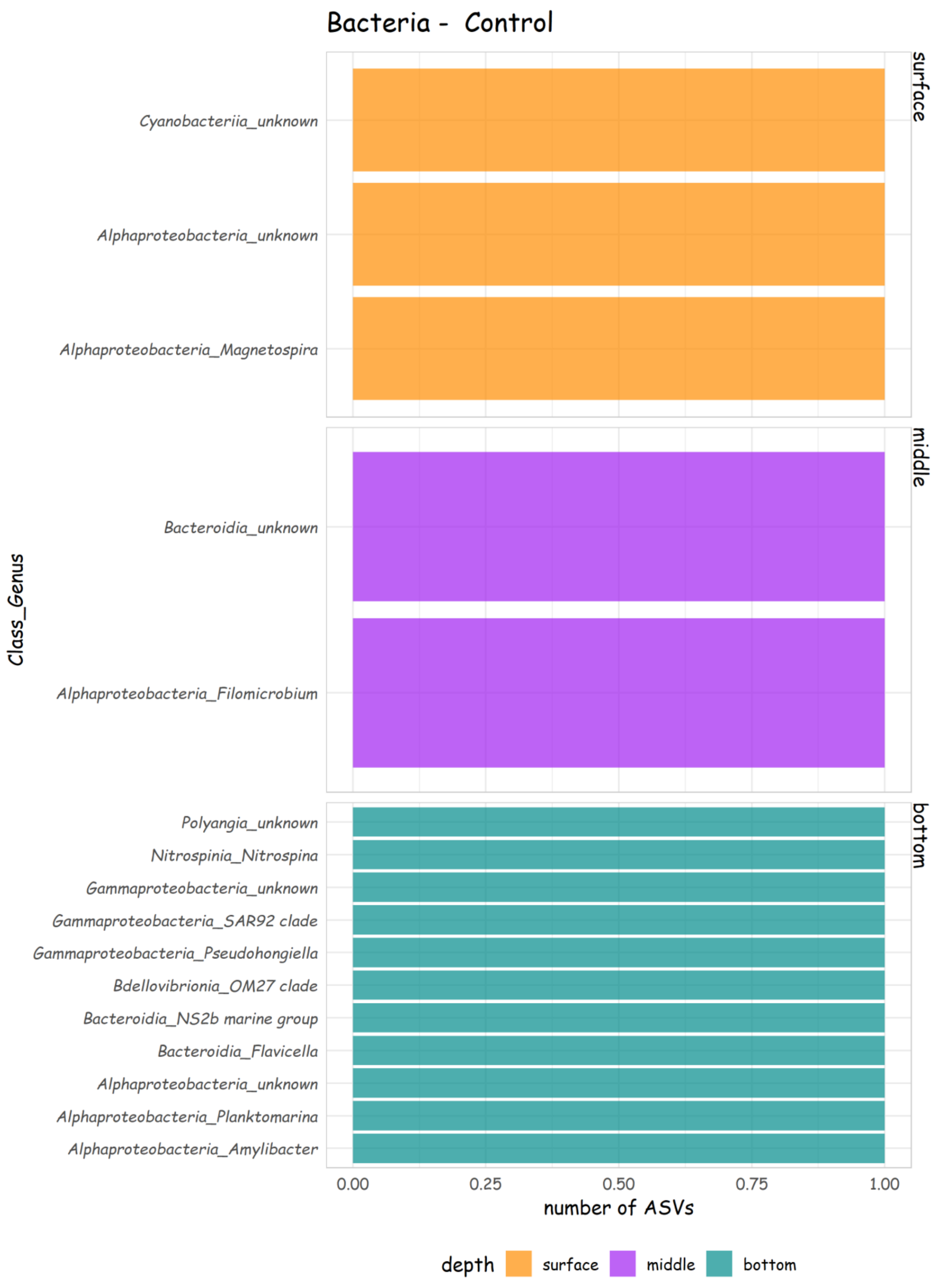


Fig. S7 – Number of ASVs found in ALDEx2 and DESeq2 analyses with relative abundance ≥ 60% in control sites.

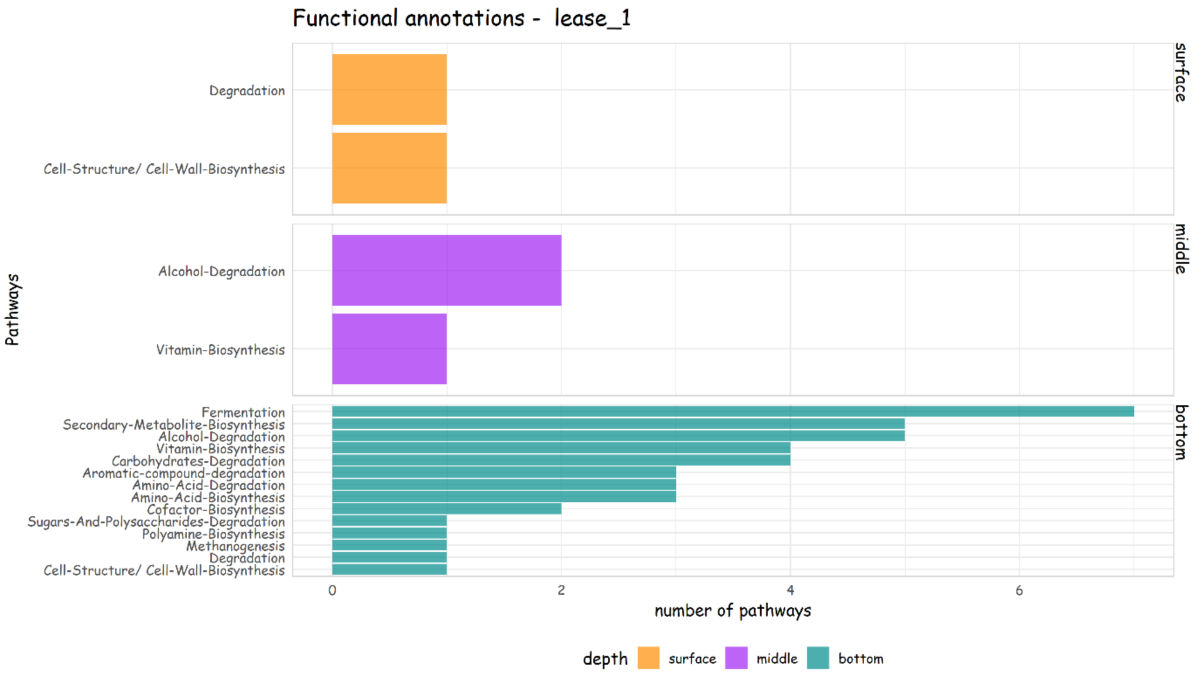


Fig. S8 – Number of functional annotations found in ALDEx2 and DESeq2 analyses with relative abundance $\geq$ 60% in Lease 1.

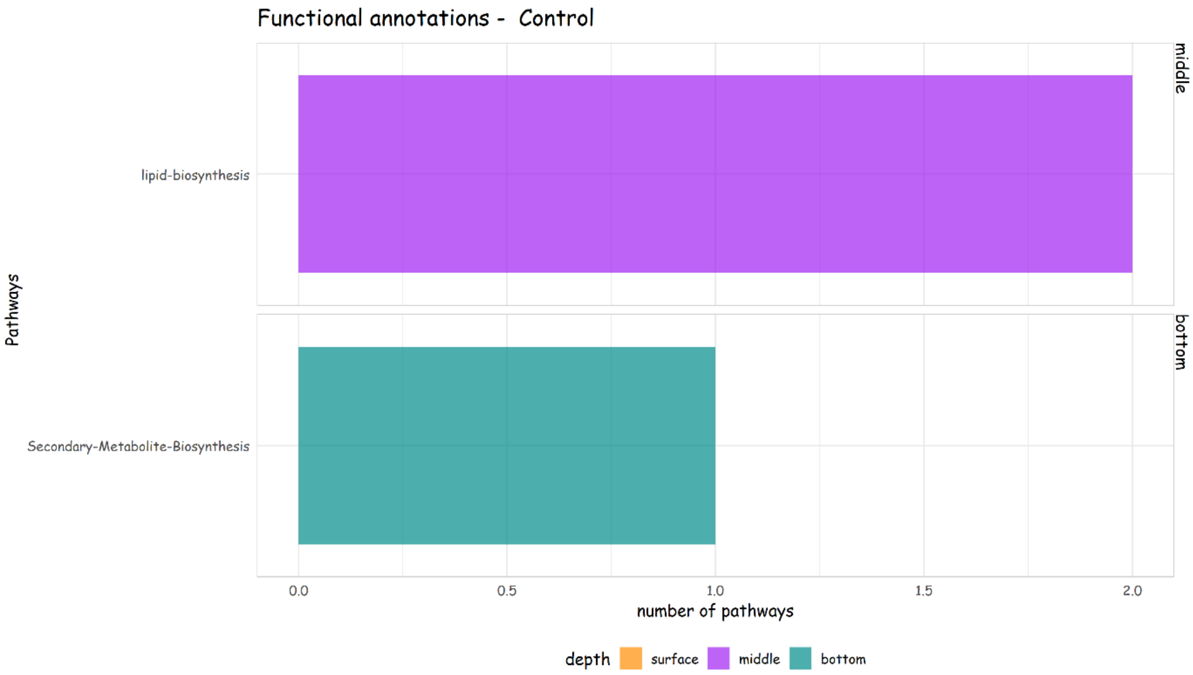


Fig. S9 – Number of functional annotations found in ALDEx2 and DESeq2 analyses with relative abundance $\geq$ 60% in control sites.

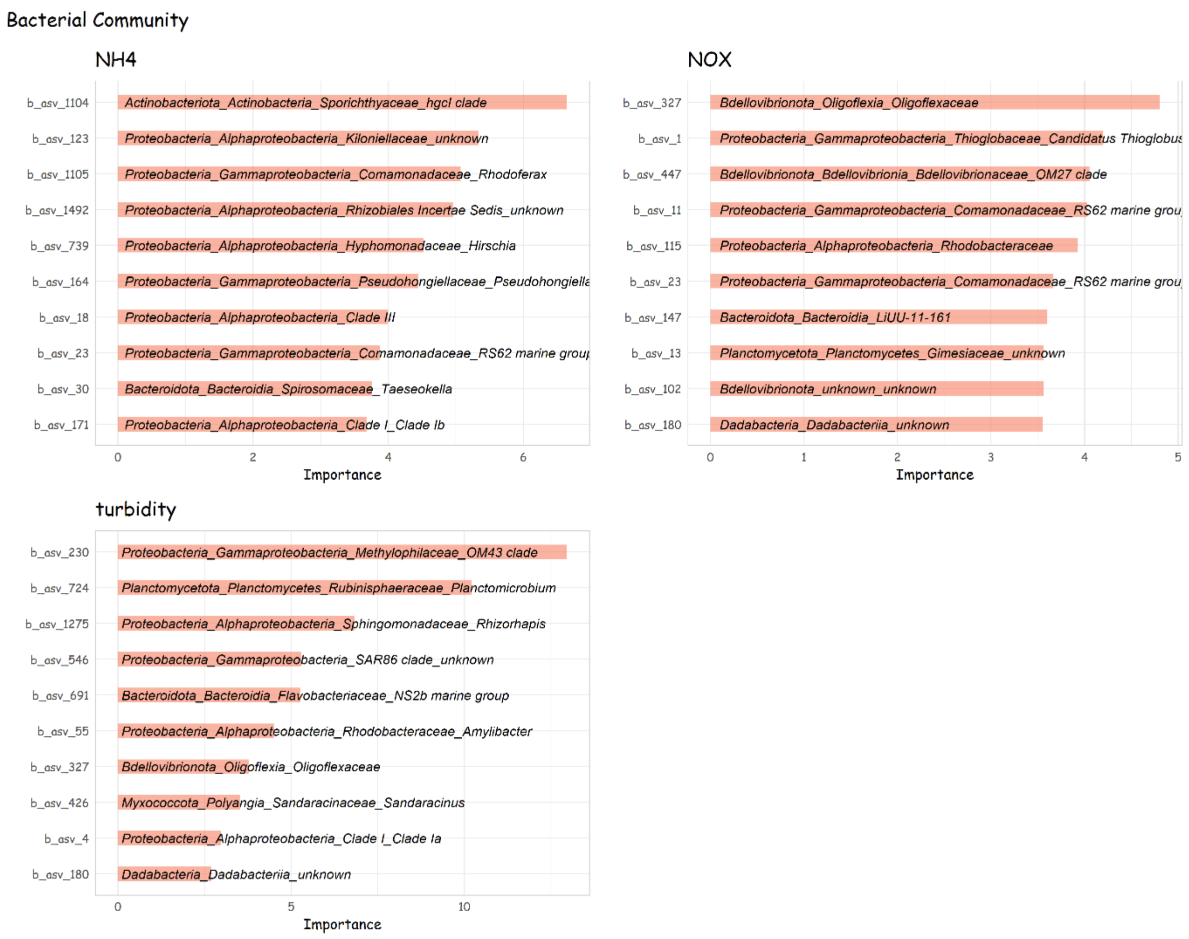


Fig. S10 – The 10 most important ASVs based on the residuals sum of square in regression-based prediction model of the environmental drivers of the bacterial community (NH4, NOx and turbidity). ASV identifications are displayed in y-axis along with the respective taxonomic levels inside of each bar (phylum_class_family_genus).

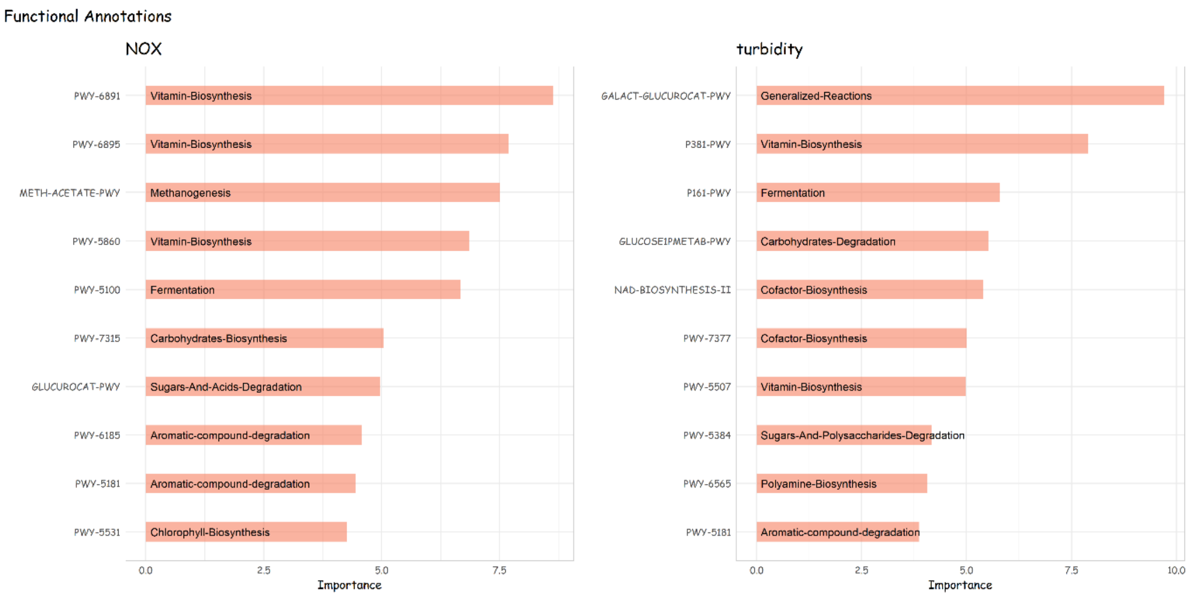


Fig. S11 – The 10 most important pathways based on the residuals sum of square in regression-based prediction model of the environmental drivers of the functional community (NOx and turbidity). Pathway id is displayed in y-axis along with the respective superclass according to MetaCyc.
