## Supplementary material for "Composition and functionality of bacterioplankton communities in marine coastal zones adjacent to finfish aquaculture": sup_tables_tables

### Supplementary Material – Tables

Table S1 – Distances (m) sampled from the cages in each lease calculated by the Haversine formula. D0 (0 m) is the water sample collected inside the cage.

| Lease | Cage | D0 | D1 | D2 | D3 | Control A | Control B | Control C |
|---|---|---|---|---|---|---|---|---|
| **1** | 1 | 0 | 16.5 | 18.2 | 82.6 | 5465 | 11417 | 1470 |
| | 2 | 0 | 23.5 | 28.5 | 82.8 | 5492 | 11434 | 1534 |
| **2** | 1 | 0 | 6.63 | 13.26 | 75 | 1890 | 5806 | 6447 |
| | 2 | 0 | 5.7 | 7.77 | 80.54 | 1826 | 5792 | 6426 |
| | 3 | 0 | 9.47 | 16.13 | 84.25 | 1484 | 5703 | 6336 |

Table S2 Environmental variables measurements.

| sample_id | site | Volume filtered (ml) | Dist(m) | Depth(m) | lat | long | conductivity | oxygen | salinity | temperature | PAR | fluorescence | turbidity | NH4 | NOX | NO2 | PO[illegible] |
|---|---|---|---|---|---|---|---|---|---|---|---|---|---|---|---|---|---|
| **T1_D0_DP1** | lease_1 | 500 | 0 | 0 | -42.3315 | 145.3922 | 5862.10 | 8.45 | 4.08 | 12.99 | 0.57 | 2.75 | 2.10 | 0.55 | 1.05 | 0.185 | 0.0[illegible] |
| **T1_D1_DP1** | lease_1 | 800 | 16.5 | 3 | -42.3315 | 145.3924 | 6185.44 | 9.72 | 4.57 | 12.50 | 0.46 | 2.77 | 1.84 | 0.54 | 2.35 | 0.116 | -0.0[illegible] |
| **T1_D1_DP2** | lease_1 | 1300 | 16.5 | 15 | -42.3315 | 145.3924 | 34352.33 | 3.64 | 27.98 | 14.05 | 0.00 | 1.29 | 0.34 | 0.08 | 7.25 | 0.007 | 0.1[illegible] |
| **T1_D1_DP3** | lease_1 | 2000 | 16.5 | 36 | -42.3315 | 145.3924 | 36953.40 | 0.59 | 29.98 | 14.51 | 0.00 | 1.27 | 0.75 | 0.07 | 8.88 | 0.021 | 0.4[illegible] |
| **T1_D2_DP1** | lease_1 | 70 | 18.2 | 3 | -42.3314 | 145.3924 | 9411.00 | 7.41 | 7.29 | 12.85 | 1.41 | 2.76 | 1.90 | 0.65 | 2.73 | 0.058 | 0.0[illegible] |
| **T1_D2_DP2** | lease_1 | 2000 | 18.2 | 15 | -42.3314 | 145.3924 | 34550.25 | 3.29 | 28.13 | 14.10 | 0.00 | 1.29 | 0.35 | 0.26 | 6.42 | 0.035 | 0.1[illegible] |
| **T1_D2_DP3** | lease_1 | 2000 | 18.2 | 36 | -42.3314 | 145.3924 | 36956.00 | 0.56 | 29.99 | 14.51 | 0.00 | 1.28 | 0.75 | 0.15 | 6.85 | 0.036 | 0.3[illegible] |
| **T1_D3_DP1** | lease_1 | 600 | 82.6 | 3 | -42.3315 | 145.3932 | 7485.40 | 7.10 | 5.66 | 12.90 | 0.02 | 2.74 | 1.90 | 0.61 | 1.54 | 0.302 | 0 |
| **T1_D3_DP2** | lease_1 | 1800 | 82.6 | 15 | -42.3315 | 145.3932 | 35030.75 | 3.19 | 28.54 | 14.14 | 0.00 | 1.29 | 0.35 | 0.18 | 6.62 | 0.097 | 0.1[illegible] |
| **T1_D3_DP3** | lease_1 | 2000 | 82.6 | 35 | -42.3315 | 145.3932 | 36935.33 | 0.66 | 29.97 | 14.51 | 0.00 | 1.27 | 0.43 | 0.11 | 7.99 | 0.163 | 0.3[illegible] |
| **T2_D0_DP1** | lease_1 | 500 | 0 | 0 | -42.3319 | 145.3916 | 5455.67 | 8.74 | 3.94 | 13.15 | 0.62 | 2.75 | 2.15 | 2.07 | 0.85 | 0.225 | 0.0[illegible] |
| **T2_D1_DP1** | lease_1 | 900 | 23.5 | 2 | -42.3321 | 145.3918 | 2434.73 | 9.36 | 1.70 | 12.62 | 4.11 | 3.11 | 2.58 | 0.43 | 0.41 | 0.21 | 0.0[illegible] |
| **T2_D1_DP2** | lease_1 | 1400 | 23.5 | 15 | -42.3321 | 145.3918 | 32751.33 | 4.90 | 26.70 | 13.83 | 0.00 | 1.31 | 0.36 | 0.09 | 5.52 | 0.077 | 0.1[illegible] |
| **T2_D1_DP3** | lease_1 | 700 | 23.5 | 32 | -42.3321 | 145.3918 | 36889.00 | 0.63 | 29.93 | 14.51 | 0.00 | 1.27 | 0.40 | 1.76 | 6.54 | 0.206 | 0.3[illegible] |
| **T2_D2_DP1** | lease_1 | 1000 | 28.5 | 3 | -42.3318 | 145.3919 | 7128.50 | 8.46 | 5.32 | 13.17 | 0.57 | 2.76 | 2.18 | 1.8 | 1.75 | 0.274 | 0.0[illegible] |
| **T2_D2_DP2** | lease_1 | 2000 | 28.5 | 15 | -42.3318 | 145.3919 | 34202.00 | 3.14 | 27.89 | 14.00 | 0.00 | 1.28 | 0.34 | 0.09 | 6.78 | 0.097 | 0.1[illegible] |
| **T2_D2_DP3** | lease_1 | 2000 | 28.5 | 35 | -42.3318 | 145.3919 | 36967.40 | 0.52 | 29.99 | 14.51 | 0.00 | 1.28 | 2.53 | 0.59 | 7.76 | 0.212 | 0.3[illegible] |

| sample_id | site | Volume filtered (ml) | Dist(m) | Depth(m) | lat | long | conductivity | oxygen | salinity | temperature | PAR | fluorescence | turbidity | NH4 | NOX | NO2 | PO4 |
|---|---|---|---|---|---|---|---|---|---|---|---|---|---|---|---|---|---|
| **T2_D3_DP1** | lease_1 | 1100 | 82.8 | 3 | -42.332 | 145.3926 | 9358.57 | 7.63 | 7.06 | 12.95 | 0.01 | 2.67 | 1.78 | 1.54 | 1.83 | 0.254 | [illegible] |
| **T2_D3_DP2** | lease_1 | 2000 | 82.8 | 15 | -42.332 | 145.3926 | 34745.50 | 2.44 | 28.33 | 14.07 | 0.00 | 1.28 | 0.36 | 0.32 | 6.71 | 0.08 | [illegible] |
| **T2_D3_DP3** | lease_1 | 600 | 82.8 | 35 | -42.332 | 145.3926 | 36935.50 | 0.51 | 29.99 | 14.49 | 0.00 | 1.27 | 1.90 | 1.31 | 7.19 | 0.186 | [illegible] |
| **T3_D0_DP1** | lease_2 | 500 | 0 | 0 | -42.2912 | 145.3474 | 5904.89 | 8.41 | 4.12 | 13.28 | 0.49 | 2.70 | 2.05 | 1.61 | 1.04 | 0.294 | [illegible] |
| **T3_D1_DP1** | lease_2 | 700 | 6.63 | 3 | -42.2913 | 145.3475 | 5790.11 | 7.76 | 3.95 | 13.62 | 0.28 | 2.64 | 1.91 | 3 | 1.16 | 0.302 | [illegible] |
| **T3_D1_DP2** | lease_2 | 2000 | 6.63 | 15 | -42.2913 | 145.3475 | 24159.79 | 5.35 | 17.91 | 13.69 | 0.05 | 1.57 | 0.67 | 0.56 | 6.02 | 0.083 | [illegible] |
| **T3_D1_DP3** | lease_2 | 2000 | 6.63 | 25 | -42.2913 | 145.3475 | 36287.82 | 1.13 | 29.58 | 14.42 | 0.00 | 1.27 | 0.42 | 0.07 | 7.88 | 0.089 | [illegible] |
| **T3_D2_DP1** | lease_2 | 600 | 13.26 | 3 | -42.2913 | 145.3475 | 7205.33 | 7.48 | 5.33 | 13.44 | 0.27 | 2.63 | 1.90 | 1.54 | 1.05 | 0.344 | [illegible] |
| **T3_D2_DP2** | lease_2 | 2000 | 13.26 | 15 | -42.2913 | 145.3475 | 25555.04 | 4.97 | 18.67 | 13.69 | 0.03 | 1.51 | 0.59 | 0.05 | 6.87 | 0.072 | [illegible] |
| **T3_D2_DP3** | lease_2 | 2000 | 13.26 | 25 | -42.2913 | 145.3475 | 36436.19 | 1.05 | 29.60 | 14.42 | 0.00 | 1.26 | 0.42 | 0.06 | 7.86 | 0.076 | [illegible] |
| **T3_D3_DP1** | lease_2 | 600 | 75 | 3 | -42.2916 | 145.3481 | 7114.97 | 7.38 | 5.25 | 13.38 | 0.34 | 2.59 | 1.92 | 0.86 | 1.03 | 0.32 | [illegible] |
| **T3_D3_DP2** | lease_2 | 1400 | 75 | 10 | -42.2916 | 145.3481 | 26112.00 | 6.21 | 21.00 | 13.52 | 0.00 | 1.53 | 0.47 | 1.88 | 2.02 | 0.273 | [illegible] |
| **T3_D3_DP3** | lease_2 | 2000 | 75 | 25 | -42.2916 | 145.3481 | 36400.86 | 1.28 | 29.60 | 14.37 | 0.00 | 1.28 | 0.40 | 0.04 | 7.35 | 0.081 | [illegible] |
| **T4_D0_DP1** | lease_2 | 600 | 0 | 0 | -42.2909 | 145.3481 | 5859.57 | 7.87 | 3.98 | 13.44 | 0.51 | 2.69 | 1.97 | 1.78 | 1.09 | 0.318 | [illegible] |
| **T4_D1_DP1** | lease_2 | 600 | 5.7 | 3 | -42.291 | 145.3481 | 4351.44 | 9.20 | 3.02 | 13.72 | 0.23 | 2.71 | 2.02 | 3.2 | 1.17 | 0.343 | [illegible] |
| **T4_D1_DP2** | lease_2 | 2000 | 5.7 | 15 | -42.291 | 145.3481 | 34323.75 | 3.98 | 28.02 | 13.98 | 0.00 | 1.28 | 0.42 | 0.06 | 7.47 | 0.101 | [illegible] |
| **T4_D1_DP3** | lease_2 | 2000 | 5.7 | 20 | -42.291 | 145.3481 | 35891.00 | 1.96 | 29.23 | 14.25 | 0.00 | 1.29 | 0.38 | 0.1 | 7.85 | 0.123 | [illegible] |
| **T4_D2_DP1** | lease_2 | 700 | 7.77 | 3 | -42.291 | 145.3481 | 4710.39 | 7.88 | 3.29 | 13.62 | 0.42 | 2.72 | 1.82 | 3.1 | 1.06 | 0.304 | [illegible] |
| **T4_D2_DP2** | lease_2 | 2000 | 7.77 | 15 | -42.291 | 145.3481 | 35176.71 | 1.65 | 28.67 | 14.14 | 0.00 | 1.29 | 0.48 | 0.08 | 6.91 | 0.104 | [illegible] |
| **T4_D2_DP3** | lease_2 | 2000 | 7.77 | 26 | -42.291 | 145.3481 | 36677.40 | 0.63 | 29.77 | 14.47 | 0.00 | 1.26 | 0.44 | 0.29 | 7.82 | 0.174 | [illegible] |
| **T4_D3_DP1** | lease_2 | 700 | 80.54 | 3 | -42.2914 | 145.3486 | 12194.40 | 5.79 | 9.50 | 13.51 | 0.36 | 2.56 | 1.77 | 1.9 | 1.24 | 0.335 | [illegible] |
| **T4_D3_DP2** | lease_2 | 2000 | 80.54 | 15 | -42.2914 | 145.3486 | 35174.50 | 3.10 | 28.65 | 14.15 | 0.00 | 1.28 | 0.38 | 0.06 | 6.94 | 0.102 | [illegible] |
| **T4_D3_DP3** | lease_2 | 2000 | 80.54 | 25 | -42.2914 | 145.3486 | 36568.33 | 1.15 | 29.69 | 14.45 | 0.00 | 1.27 | 0.45 | 0.1 | 7.7 | 0.106 | [illegible] |
| **T5_D0_DP1** | lease_2 | 500 | 0 | 0 | -42.2891 | 145.3517 | 5932.86 | 7.76 | 4.08 | 13.46 | 0.46 | 2.69 | 1.96 | 1.6 | 1.11 | 0.365 | [illegible] |
| **T5_D1_DP1** | lease_2 | 800 | 9.47 | 3 | -42.2891 | 145.3518 | 5555.19 | 7.19 | 3.94 | 13.77 | 0.02 | 2.62 | 1.89 | 2.31 | 1.19 | 0.35 | [illegible] |
| **T5_D1_DP2** | lease_2 | 2000 | 9.47 | 15 | -42.2891 | 145.3518 | 33720.00 | 1.47 | 27.34 | 14.19 | 0.00 | 1.28 | 0.37 | 1.08 | 4.04 | 0.162 | [illegible] |
| **T5_D1_DP3** | lease_2 | 2000 | 9.47 | 35 | -42.2891 | 145.3518 | 35801.40 | 0.45 | 28.97 | 14.49 | 0.00 | 1.24 | 0.61 | 0.14 | 6.01 | 0.254 | [illegible] |
| **T5_D2_DP1** | lease_2 | 600 | 16.13 | 3 | -42.2891 | 145.3518 | 8038.60 | 6.69 | 5.99 | 13.56 | 0.33 | 2.55 | 1.76 | 1.78 | 1.14 | 0.327 | [illegible] |
| **T5_D2_DP2** | lease_2 | 2000 | 16.13 | 15 | -42.2891 | 145.3518 | 34273.50 | 4.40 | 27.94 | 14.02 | 0.00 | 1.29 | 0.36 | 0.07 | 6.64 | 0.119 | [illegible] |

| sample_id | site | Volume filtered (ml) | Dist(m) | Depth(m) | lat | long | conductivity | oxygen | salinity | temperature | PAR | fluorescence | turbidity | NH4 | NOX | NO2 | PO4 |
|---|---|---|---|---|---|---|---|---|---|---|---|---|---|---|---|---|---|
| **T5_D2_DP3** | lease_2 | 2000 | 16.13 | 35 | -42.2891 | 145.3518 | 36860.00 | 0.68 | 29.93 | 14.47 | 0.00 | 1.24 | 0.53 | 0.16 | 7.67 | 0.244 | 0.3[illegible] |
| **T5_D3_DP1** | lease_2 | 500 | 84.25 | 3 | -42.2897 | 145.3523 | 5802.93 | 9.04 | 4.13 | 13.63 | 0.32 | 2.51 | 2.14 | 1.91 | 0.97 | 0.289 | 0.0[illegible] |
| **T5_D3_DP2** | lease_2 | 2000 | 84.25 | 15 | -42.2897 | 145.3523 | 34086.00 | 4.28 | 27.81 | 13.96 | 0.00 | 1.28 | 0.34 | 0.18 | 5.63 | 0.19 | 0.0[illegible] |
| **T5_D3_DP3** | lease_2 | 2000 | 84.25 | 35 | -42.2897 | 145.3523 | 36972.75 | 0.79 | 30.08 | 14.41 | 0.00 | 1.22 | 1.32 | 0.04 | 7.97 | 0.145 | 0.3[illegible] |
| **CA_DP1** | Control | 1100 | >1500 | 3 | -42.2855 | 145.369 | 14858.70 | 6.86 | 11.51 | 13.80 | 0.15 | 2.36 | 1.49 | 1.01 | 1.17 | 0.238 | 0.0[illegible] |
| **CA_DP2** | Control | 2000 | >1500 | 15 | -42.2855 | 145.369 | 34385.00 | 3.39 | 28.12 | 13.91 | 0.00 | 1.27 | 0.40 | 0.25 | 4.09 | 0.102 | 0.0[illegible] |
| **CA_DP3** | Control | 2000 | >1500 | 35 | -42.2855 | 145.369 | 34954.67 | 0.97 | 28.26 | 14.43 | 0.00 | 1.24 | 0.37 | 0.08 | 7.58 | 0.084 | 0.2[illegible] |
| **CB_DP1** | Control | 1600 | >1500 | 4 | -42.2418 | 145.3249 | 9477.53 | 9.67 | 6.98 | 13.52 | 0.01 | 2.25 | 1.09 | 0.96 | 1.8 | 0.292 | 0.0[illegible] |
| **CB_DP2** | Control | 2000 | >1500 | 15 | -42.2418 | 145.3249 | 35303.67 | 2.64 | 28.83 | 14.08 | 0.00 | 1.25 | 0.48 | 0.22 | 4 | 0.106 | 0.0[illegible] |
| **CB_DP3** | Control | 2000 | >1500 | 37 | -42.2418 | 145.3249 | 36900.75 | 1.37 | 30.04 | 14.38 | 0.00 | 1.21 | 0.52 | 0.1 | 7.75 | 0.118 | 0.2[illegible] |
| **CC_DP1** | Control | 600 | >1500 | 2 | -42.327 | 145.409 | 2920.53 | 10.61 | 2.06 | 12.43 | 2.41 | 3.03 | 3.37 | 0.42 | 0.9 | 0.325 | 0.0[illegible] |
| **CC_DP2** | Control | 800 | >1500 | 15 | -42.327 | 145.409 | 34060.40 | 3.84 | 27.78 | 13.98 | 0.00 | 1.28 | 0.34 | 0.2 | 6.13 | 0.104 | 0.1[illegible] |
| **CC_DP3** | Control | 800 | >1500 | 34 | -42.327 | 145.409 | 36931.83 | 0.59 | 29.98 | 14.50 | 0.00 | 1.27 | 0.39 | 0.13 | 7.52 | 0.136 | 0.2[illegible] |

Table S3 – Summary of nutrient concentrations (mean ± standard deviation, minimum and maximum) at each sampled site. Distance (D0-D4) and depth (DP1-DP3) categories are detailed in Table S1 and Table S2, respectively.

| site | Distance | Depth | $NH_4$ | | $NO_2$ | | NOx | | $PO_4$ | |
|---|---|---|---|---|---|---|---|---|---|---|
| | | | mean ± sd | max-min | mean±sd | max-min | mean±sd | max-min | mean±sd | max-min |
| **Lease_1** | D0 | DP1 | 1.31 ± 1.07 | 0.6 - 2.1 | 0.21 ± 0.03 | 0.2 - 0.2 | 0.95 ± 0.14 | 0.8 - 1 | 0.02 ± 0.01 | 0 - 0 |
| | D1 | DP1 | 0.48 ± 0.08 | 0.4 - 0.5 | 0.16 ± 0.07 | 0.1 - 0.2 | 1.38 ± 1.37 | 0.4 - 2.4 | 0 ± 0.03 | 0 - 0 |
| | D1 | DP2 | 0.08 ± 0.01 | 0.1 - 0.1 | 0.04 ± 0.05 | 0 - 0.1 | 6.38 ± 1.22 | 5.5 - 7.2 | 0.14 ± 0.01 | 0.1 - 0.1 |
| | D1 | DP3 | 0.92 ± 1.2 | 0.1 - 1.8 | 0.11 ± 0.13 | 0 - 0.2 | 7.71 ± 1.65 | 6.5 - 8.9 | 0.38 ± 0.04 | 0.3 - 0.4 |
| | D2 | DP1 | 1.23 ± 0.81 | 0.7 - 1.8 | 0.17 ± 0.15 | 0.1 - 0.3 | 2.24 ± 0.69 | 1.8 - 2.7 | 0.03 ± 0.01 | 0 - 0 |
| | D2 | DP2 | 0.17 ± 0.12 | 0.1 - 0.3 | 0.07 ± 0.04 | 0 - 0.1 | 6.6 ± 0.25 | 6.4 - 6.8 | 0.14 ± 0.01 | 0.1 - 0.1 |
| | D2 | DP3 | 0.37 ± 0.31 | 0.1 - 0.6 | 0.12 ± 0.12 | 0 - 0.2 | 7.3 ± 0.64 | 6.8 - 7.8 | 0.34 ± 0.04 | 0.3 - 0.4 |
| | D3 | DP1 | 1.07 ± 0.66 | 0.6 - 1.5 | 0.28 ± 0.03 | 0.3 - 0.3 | 1.69 ± 0.21 | 1.5 - 1.8 | 0 ± 0.01 | 0 - 0 |
| | D3 | DP2 | 0.25 ± 0.1 | 0.2 - 0.3 | 0.09 ± 0.01 | 0.1 - 0.1 | 6.66 ± 0.06 | 6.6 - 6.7 | 0.12 ± 0.01 | 0.1 - 0.1 |
| | D3 | DP3 | 0.71 ± 0.85 | 0.1 - 1.3 | 0.17 ± 0.02 | 0.2 - 0.2 | 7.59 ± 0.57 | 7.2 - 8 | 0.37 ± 0.1 | 0.3 - 0.4 |
| **Lease_** | D0 | DP1 | 1.66 ± 0.1 | 1.6 - 1.8 | 0.33 ± 0.04 | 0.3 - 0.4 | 1.08 ± 0.04 | 1 - 1.1 | 0.04 ± 0.01 | 0 - 0 |

| | | | | | | | | | | |
|---|---|---|---|---|---|---|---|---|---|---|
| | D1 | DP1 | 2.84 ± 0.47 | 2.3 - 3.2 | 0.33 ± 0.03 | 0.3 - 0.3 | 1.17 ± 0.02 | 1.2 - 1.2 | 0.06 ± 0.03 | 0 - 0.1 |
| | D1 | DP2 | 0.57 ± 0.51 | 0.1 - 1.1 | 0.12 ± 0.04 | 0.1 - 0.2 | 5.84 ± 1.72 | 4 - 7.5 | 0.13 ± 0.05 | 0.1 - 0.2 |
| | D1 | DP3 | 0.1 ± 0.04 | 0.1 - 0.1 | 0.16 ± 0.09 | 0.1 - 0.3 | 7.25 ± 1.07 | 6 - 7.9 | 0.26 ± 0.01 | 0.2 - 0.3 |
| | D2 | DP1 | 2.14 ± 0.84 | 1.5 - 3.1 | 0.32 ± 0.02 | 0.3 - 0.3 | 1.08 ± 0.05 | 1 - 1.1 | 0.04 ± 0.03 | 0 - 0.1 |
| | D2 | DP2 | 0.07 ± 0.02 | 0 - 0.1 | 0.1 ± 0.02 | 0.1 - 0.1 | 6.81 ± 0.15 | 6.6 - 6.9 | 0.13 ± 0.01 | 0.1 - 0.1 |
| | D2 | DP3 | 0.17 ± 0.12 | 0.1 - 0.3 | 0.16 ± 0.08 | 0.1 - 0.2 | 7.78 ± 0.1 | 7.7 - 7.9 | 0.31 ± 0.05 | 0.3 - 0.4 |
| | D3 | DP1 | 1.56 ± 0.6 | 0.9 - 1.9 | 0.31 ± 0.02 | 0.3 - 0.3 | 1.08 ± 0.14 | 1 - 1.2 | 0.04 ± 0.01 | 0 - 0 |
| | D3 | DP2 | 0.71 ± 1.02 | 0.1 - 1.9 | 0.19 ± 0.09 | 0.1 - 0.3 | 4.86 ± 2.55 | 2 - 6.9 | 0.07 ± 0.06 | 0 - 0.1 |
| | D3 | DP3 | 0.06 ± 0.03 | 0 - 0.1 | 0.11 ± 0.03 | 0.1 - 0.1 | 7.67 ± 0.31 | 7.3 - 8 | 0.24 ± 0.09 | 0.1 - 0.3 |
| **Control** | D4 | DP1 | 0.8 ± 0.33 | 0.4 - 1 | 0.28 ± 0.04 | 0.2 - 0.3 | 1.29 ± 0.46 | 0.9 - 1.8 | 0.02 ± 0.01 | 0 - 0 |
| | D4 | DP2 | 0.22 ± 0.03 | 0.2 - 0.2 | 0.1 ± 0 | 0.1 - 0.1 | 4.74 ± 1.2 | 4 - 6.1 | 0.09 ± 0.03 | 0.1 - 0.1 |
| | D4 | DP3 | 0.1 ± 0.03 | 0.1 - 0.1 | 0.11 ± 0.03 | 0.1 - 0.1 | 7.62 ± 0.12 | 7.5 - 7.8 | 0.27 ± 0.01 | 0.3 - 0.3 |

Table S4 – Summary of the other environmental variables (mean ± standard deviation, minimum and maximum) at each sampled site. Distance (D0-D4) and depth (DP1-DP3) categories are detailed in Table S1 and Table S2, respectively.

| **site** | **Dist** | **Depth** | **conductivity** | | **fluorescence** | | **oxygen** | | **PAR** | | **salinity** | | **temperature** | | **turbidity** | |
|---|---|---|---|---|---|---|---|---|---|---|---|---|---|---|---|---|
| | | | **mean ± sd** | **min-max** | **mean ± sd** | **min-max** | **mean ± sd** | **min-max** | **mean ± sd** | **min-max** | **mean ± sd** | **min-max** | **mean ± sd** | **min-max** | **mean ± sd** | **min-max** |
| | D0 | DP1 | 5658.89 ± 287.39 | 5455.7 - 5862.1 | 2.75 ± 0 | 2.7 - 2.7 | 8.59 ± 0.21 | 8.4 - 8.7 | 0.59 ± 0.03 | 0.6 - 0.6 | 4.01 ± 0.1 | 3.9 - 4.1 | 13.07 ± 0.11 | 13 - 13.2 | 2.12 ± 0.04 | 2.1 - 2.2 |
| | D1 | DP1 | 4310.08 ± 2652.15 | 2434.7 - 6185.4 | 2.94 ± 0.24 | 2.8 - 3.1 | 9.54 ± 0.25 | 9.4 - 9.7 | 2.28 ± 2.58 | 0.5 - 4.1 | 3.13 ± 2.03 | 1.7 - 4.6 | 12.56 ± 0.09 | 12.5 - 12.6 | 2.21 ± 0.52 | 1.8 - 2.6 |
| **Lease_1** | D1 | DP2 | 33551.83 ± 1132.08 | 32751.3 - 34352.3 | 1.3 ± 0.02 | 1.3 - 1.3 | 4.27 ± 0.89 | 3.6 - 4.9 | 0 ± 0 | 0 - 0 | 27.34 ± 0.91 | 26.7 - 28 | 13.94 ± 0.16 | 13.8 - 14.1 | 0.35 ± 0.01 | 0.3 - 0.4 |
| | D1 | DP3 | 36921.2 ± 45.54 | 36889 - 36953.4 | 1.27 ± 0.01 | 1.3 - 1.3 | 0.61 ± 0.03 | 0.6 - 0.6 | 0 ± 0 | 0 - 0 | 29.95 ± 0.04 | 29.9 - 30 | 14.51 ± 0 | 14.5 - 14.5 | 0.57 ± 0.25 | 0.4 - 0.8 |
| | D2 | DP1 | 8269.75 ± 1613.97 | 7128.5 - 9411 | 2.76 ± 0 | 2.8 - 2.8 | 7.94 ± 0.74 | 7.4 - 8.5 | 0.99 ± 0.59 | 0.6 - 1.4 | 6.3 ± 1.39 | 5.3 - 7.3 | 13.01 ± 0.22 | 12.9 - 13.2 | 2.04 ± 0.2 | 1.9 - 2.2 |
| | D2 | DP2 | 34376.12 ± 246.25 | 34202 - 34550.2 | 1.29 ± 0 | 1.3 - 1.3 | 3.21 ± 0.1 | 3.1 - 3.3 | 0 ± 0 | 0 - 0 | 28.01 ± 0.17 | 27.9 - 28.1 | 14.05 ± 0.07 | 14 - 14.1 | 0.34 ± 0.01 | 0.3 - 0.3 |

| | | | | | | | | | | | | | | | | |
|---|---|---|---|---|---|---|---|---|---|---|---|---|---|---|---|---|
| | D2 | DP3 | 36961.7 ± 8.06 | 36956 - 36967.4 | 1.28 ± 0.01 | 1.3 - 1.3 | 0.54 ± 0.03 | 0.5 - 0.6 | 0 ± 0 | 0 - 0 | 29.99 ± 0.01 | 30 - 30 | 14.51 ± 0 | 14.5 - 14.5 | 1.64 ± 1.26 | 0.7 - 2.5 |
| | D3 | DP1 | 8421.98 ± 1324.53 | 7485.4 - 9358.6 | 2.71 ± 0.05 | 2.7 - 2.7 | 7.37 ± 0.38 | 7.1 - 7.6 | 0.02 ± 0.01 | 0 - 0 | 6.36 ± 0.99 | 5.7 - 7.1 | 12.92 ± 0.03 | 12.9 - 12.9 | 1.84 ± 0.09 | 1.8 - 1.9 |
| | D3 | DP2 | 34888.12 ± 201.7 | 34745.5 - 35030.8 | 1.29 ± 0 | 1.3 - 1.3 | 2.82 ± 0.53 | 2.4 - 3.2 | 0 ± 0 | 0 - 0 | 28.43 ± 0.15 | 28.3 - 28.5 | 14.1 ± 0.05 | 14.1 - 14.1 | 0.35 ± 0 | 0.4 - 0.4 |
| | D3 | DP3 | 36935.42 ± 0.12 | 36935.3 - 36935.5 | 1.27 ± 0 | 1.3 - 1.3 | 0.59 ± 0.11 | 0.5 - 0.7 | 0 ± 0 | 0 - 0 | 29.98 ± 0.01 | 30 - 30 | 14.5 ± 0.02 | 14.5 - 14.5 | 1.16 ± 1.04 | 0.4 - 1.9 |
| **Lease_2** | D0 | DP1 | 5899.11 ± 36.99 | 5859.6 - 5932.9 | 2.69 ± 0.01 | 2.7 - 2.7 | 8.01 ± 0.35 | 7.8 - 8.4 | 0.49 ± 0.03 | 0.5 - 0.5 | 4.06 ± 0.07 | 4 - 4.1 | 13.39 ± 0.1 | 13.3 - 13.5 | 1.99 ± 0.05 | 2.0 - 2.0 |
| | D1 | DP1 | 5232.25 ± 771.79 | 4351.4 - 5790.1 | 2.66 ± 0.05 | 2.6 - 2.7 | 8.05 ± 1.04 | 7.2 - 9.2 | 0.17 ± 0.14 | 0 - 0.3 | 3.63 ± 0.54 | 3 - 3.9 | 13.7 ± 0.08 | 13.6 - 13.8 | 1.94 ± 0.07 | 1.9 - 2 |
| | D1 | DP2 | 30734.51 ± 5701.87 | 24159.8 - 34323.8 | 1.38 ± 0.17 | 1.3 - 1.6 | 3.6 ± 1.97 | 1.5 - 5.3 | 0.02 ± 0.03 | 0 - 0 | 24.42 ± 5.65 | 17.9 - 28 | 13.95 ± 0.25 | 13.7 - 14.2 | 0.49 ± 0.16 | 0.4 - 0.7 |
| | D1 | DP3 | 35993.41 ± 258.87 | 35801.4 - 36287.8 | 1.26 ± 0.03 | 1.2 - 1.3 | 1.18 ± 0.76 | 0.4 - 2 | 0 ± 0 | 0 - 0 | 29.26 ± 0.3 | 29 - 29.6 | 14.38 ± 0.12 | 14.2 - 14.5 | 0.47 ± 0.12 | 0.4 - 0.6 |
| | D2 | DP1 | 6651.44 ± 1731.86 | 4710.4 - 8038.6 | 2.64 ± 0.08 | 2.6 - 2.7 | 7.35 ± 0.6 | 6.7 - 7.9 | 0.34 ± 0.07 | 0.3 - 0.4 | 4.87 ± 1.41 | 3.3 - 6 | 13.54 ± 0.09 | 13.4 - 13.6 | 1.82 ± 0.07 | 1.8 - 1.9 |
| | D2 | DP2 | 31668.42 ± 5313.57 | 25555 - 35176.7 | 1.36 ± 0.13 | 1.3 - 1.5 | 3.68 ± 1.78 | 1.7 - 5 | 0.01 ± 0.01 | 0 - 0 | 25.09 ± 5.57 | 18.7 - 28.7 | 13.95 ± 0.23 | 13.7 - 14.1 | 0.48 ± 0.12 | 0.4 - 0.6 |
| | D2 | DP3 | 36657.86 ± 212.58 | 36436.2 - 36860 | 1.25 ± 0.01 | 1.2 - 1.3 | 0.79 ± 0.23 | 0.6 - 1.1 | 0 ± 0 | 0 - 0 | 29.77 ± 0.17 | 29.6 - 29.9 | 14.45 ± 0.03 | 14.4 - 14.5 | 0.46 ± 0.06 | 0.4 - 0.5 |
| | D3 | DP1 | 8370.76 ± 3375.72 | 5802.9 - 12194.4 | 2.55 ± 0.04 | 2.5 - 2.6 | 7.41 ± 1.62 | 5.8 - 9 | 0.34 ± 0.02 | 0.3 - 0.4 | 6.29 ± 2.84 | 4.1 - 9.5 | 13.51 ± 0.13 | 13.4 - 13.6 | 1.95 ± 0.19 | 1.8 - 2.1 |
| | D3 | DP2 | 31790.83 ± 4948.04 | 26112 - 35174.5 | 1.36 ± 0.14 | 1.3 - 1.5 | 4.53 ± 1.57 | 3.1 - 6.2 | 0 ± 0 | 0 - 0 | 25.82 ± 4.2 | 21 - 28.7 | 13.88 ± 0.33 | 13.5 - 14.2 | 0.4 ± 0.07 | 0.3 - 0.5 |
| | D3 | DP3 | 36647.31 ± 294.01 | 36400.9 - 36972.8 | 1.26 ± 0.03 | 1.2 - 1.3 | 1.07 ± 0.25 | 0.8 - 1.3 | 0 ± 0 | 0 - 0 | 29.79 ± 0.25 | 29.6 - 30.1 | 14.41 ± 0.04 | 14.4 - 14.4 | 0.72 ± 0.52 | 0.4 - 1.3 |

| | | | | | | | | | | | | | | | | |
|---|---|---|---|---|---|---|---|---|---|---|---|---|---|---|---|---|
| **Control** | D4 | DP1 | 9085.59 ± 5978.73 | 2920.5 - 14858.7 | 2.55 ± 0.42 | 2.3 - 3 | 9.05 ± 1.95 | 6.9 - 10.6 | 0.86 ± 1.34 | 0 - 2.4 | 6.85 ± 4.73 | 2.1 - 11.5 | 13.25 ± 0.73 | 12.4 - 13.8 | 1.98 ± 1.22 | 1.1 - 3.4 |
| | D4 | DP2 | 34583.02 ± 644.85 | 34060.4 - 35303.7 | 1.27 ± 0.02 | 1.3 - 1.3 | 3.29 ± 0.6 | 2.6 - 3.8 | 0 ± 0 | 0 - 0 | 28.24 ± 0.53 | 27.8 - 28.8 | 13.99 ± 0.09 | 13.9 - 14.1 | 0.41 ± 0.07 | 0.3 - 0.5 |
| | D4 | DP3 | 36262.42 ± 1132.65 | 34954.7 - 36931.8 | 1.24 ± 0.03 | 1.2 - 1.3 | 0.98 ± 0.39 | 0.6 - 1.4 | 0 ± 0 | 0 - 0 | 29.42 ± 1.01 | 28.3 - 30 | 14.44 ± 0.06 | 14.4 - 14.5 | 0.43 ± 0.08 | 0.4 - 0.5 |

Table S5 – Differentially abundant ASVs in ease 1

| Depth | Phylum | Class | Order | Family | Genus | #ASVs |
|---|---|---|---|---|---|---|
| bottom | Proteobacteria | Gammaproteobacteria | Enterobacterales | Psychromonadaceae | Psychromonas | 16 |
| bottom | Spirochaetota | Spirochaetia | Spirochaetales | Spirochaetaceae | Spirochaeta 2 | 10 |
| bottom | Campylobacterota | Campylobacteria | Campylobacterales | Arcobacteraceae | unknown | 5 |
| bottom | Fusobacteriota | Fusobacteriia | Fusobacteriales | Leptotrichiaceae | unknown | 5 |
| bottom | Bacteroidota | Bacteroidia | Bacteroidales | unknown | unknown | 4 |
| bottom | Fusobacteriota | Fusobacteriia | Fusobacteriales | Leptotrichiaceae | Hypnocyclicus | 4 |
| bottom | Bacteroidota | Bacteroidia | Bacteroidales | Prolixibacteraceae | Draconibacterium | 3 |
| bottom | Proteobacteria | Gammaproteobacteria | Steroidobacterales | Woeseiaceae | Woeseia | 3 |
| bottom | Bacteroidota | Bacteroidia | Bacteroidales | Bacteroidetes BD2-2 | unknown | 2 |
| bottom | Bacteroidota | Bacteroidia | Bacteroidales | Marinifilaceae | unknown | 2 |
| bottom | Bacteroidota | Bacteroidia | Bacteroidales | Marinilabiliaceae | unknown | 2 |
| bottom | Campylobacterota | Campylobacteria | Campylobacterales | Sulfurimonadaceae | Sulfurimonas | 2 |
| bottom | Desulfobacterota | Desulfovibrionia | Desulfovibrionales | Desulfovibrionaceae | Desulfovibrio | 2 |
| bottom | Firmicutes | Clostridia | Lachnospirales | Lachnospiraceae | unknown | 2 |
| bottom | Proteobacteria | Gammaproteobacteria | Pseudomonadales | Pseudomonadaceae | Pseudomonas | 2 |
| bottom | Verrucomicrobiota | Lentisphaeria | P.palmC41 | unknown | unknown | 2 |
| bottom | Verrucomicrobiota | Lentisphaeria | Victivallales | PRD18C08 | unknown | 2 |
| surface | Planctomycetota | Planctomycetes | Planctomycetales | Gimesiaceae | unknown | 2 |
| bottom | Acidobacteriota | Vicinamibacteria | Subgroup 17 | unknown | unknown | 1 |
| bottom | Actinobacteriota | Acidimicrobiia | Actinomarinales | unknown | unknown | 1 |
| bottom | Bacteroidota | Bacteroidia | Flavobacteriales | Flavobacteriaceae | unknown | 1 |

| Depth | Phylum | Class | Order | Family | Genus | #ASVs |
|---|---|---|---|---|---|---|
| bottom | Bacteroidota | Ignavibacteria | Ignavibacteriales | PHOS-HE36 | unknown | 1 |
| bottom | Campylobacterota | Campylobacteria | Campylobacterales | Sulfurospirillaceae | Sulfurospirillum | 1 |
| bottom | Campylobacterota | Campylobacteria | Campylobacterales | Sulfurovaceae | Sulfurovum | 1 |
| bottom | Desulfobacterota | Desulfobulbia | Desulfobulbales | Desulfocapsaceae | Desulforhopalus | 1 |
| bottom | Firmicutes | Bacilli | Erysipelotrichales | Erysipelotrichaceae | Erysipelothrix | 1 |
| bottom | Firmicutes | Bacilli | Izemoplasmatales | Izemoplasmataceae | Izimaplasma | 1 |
| bottom | Firmicutes | Clostridia | MAT-CR-H4-C10 | unknown | unknown | 1 |
| bottom | Firmicutes | Clostridia | Peptostreptococcales-Tissierellales | Fusibacteraceae | Fusibacter | 1 |
| bottom | Firmicutes | Clostridia | Peptostreptococcales-Tissierellales | unknown | unknown | 1 |
| bottom | Fusobacteriota | Fusobacteriia | Fusobacteriales | Fusobacteriaceae | unknown | 1 |
| bottom | Fusobacteriota | Fusobacteriia | Fusobacteriales | unknown | unknown | 1 |
| bottom | LCP-89 | unknown | unknown | unknown | unknown | 1 |
| bottom | Myxococcota | Polyangia | Polyangiales | Sandaracinaceae | unknown | 1 |
| bottom | Planctomycetota | Planctomycetes | Planctomycetales | Rubinisphaeraceae | Planctomicrobium | 1 |
| bottom | Planctomycetota | Planctomycetes | Planctomycetales | Rubinisphaeraceae | unknown | 1 |
| bottom | Proteobacteria | Alphaproteobacteria | Rhodobacterales | Rhodobacteraceae | Limibaculum | 1 |
| bottom | Proteobacteria | Gammaproteobacteria | Enterobacterales | Alteromonadaceae | Gayadomonas | 1 |
| bottom | Proteobacteria | Gammaproteobacteria | Enterobacterales | Kangiellaceae | unknown | 1 |
| bottom | Proteobacteria | Gammaproteobacteria | Enterobacterales | unknown | unknown | 1 |
| bottom | Proteobacteria | Gammaproteobacteria | Enterobacterales | Vibrionaceae | unknown | 1 |
| bottom | Proteobacteria | Gammaproteobacteria | Enterobacterales | Vibrionaceae | Vibrio | 1 |
| bottom | Proteobacteria | Gammaproteobacteria | Pseudomonadales | Halomonadaceae | Halomonas | 1 |
| bottom | Proteobacteria | Gammaproteobacteria | Thiomicrospirales | Thiomicrospiraceae | endosymbionts | 1 |
| bottom | Spirochaetota | Spirochaetia | Spirochaetales | Spirochaetaceae | GWE2-31-10 | 1 |
| bottom | Spirochaetota | Spirochaetia | Spirochaetales | Spirochaetaceae | unknown | 1 |
| middle | Proteobacteria | Gammaproteobacteria | Enterobacterales | Enterobacteriaceae | Escherichia-Shigella | 1 |
| middle | Proteobacteria | Gammaproteobacteria | Pseudomonadales | Halomonadaceae | Halomonas | 1 |
| middle | Proteobacteria | Gammaproteobacteria | Pseudomonadales | Pseudomonadaceae | Pseudomonas | 1 |
| surface | Bacteroidota | Bacteroidia | Flavobacteriales | Flavobacteriaceae | Flavobacterium | 1 |

| Depth | Phylum | Class | Order | Family | Genus | #ASVs |
|---|---|---|---|---|---|---|
| surface | Firmicutes | Bacilli | Erysipelotrichales | Erysipelotrichaceae | Erysipelothrix | 1 |
| surface | Patescibacteria | Gracilibacteria | Candidatus Peregrinibacteria | unknown | unknown | 1 |
| surface | Proteobacteria | Alphaproteobacteria | Rhizobiales | Beijerinckiaceae | Methylocystis | 1 |
| surface | Proteobacteria | Alphaproteobacteria | Rhizobiales | Rhizobiaceae | Mesorhizobium | 1 |
| surface | Proteobacteria | Gammaproteobacteria | Methylococcales | Methylomonadaceae | Milano-WF1B-03 | 1 |
| surface | Proteobacteria | Gammaproteobacteria | Pseudomonadales | Halomonadaceae | Halomonas | 1 |
| surface | Proteobacteria | Gammaproteobacteria | Pseudomonadales | Moraxellaceae | Acinetobacter | 1 |
| surface | Proteobacteria | Gammaproteobacteria | Pseudomonadales | Thioglobaceae | SUP05 cluster | 1 |

Table S6 – Differentially abundant ASVs in Lease 2

| Depth | Phylum | Class | Order | Family | Genus | #ASVs |
|---|---|---|---|---|---|---|
| surface | Proteobacteria | Gammaproteobacteria | Enterobacterales | Vibrionaceae | Aliivibrio | 4 |
| bottom | Bacteroidota | Bacteroidia | Flavobacteriales | Flavobacteriaceae | NS3a marine group | 1 |
| bottom | Bdellovibrionota | Bdellovibrionia | Bacteriovoracales | Bacteriovoracaceae | unknown | 1 |
| bottom | Desulfobacterota | Desulfuromonadia | PB19 | unknown | unknown | 1 |
| surface | Actinobacteriota | Actinobacteria | Frankiales | Sporichthyaceae | hgcI clade | 1 |
| surface | Actinobacteriota | Actinobacteria | Micrococcales | Microbacteriaceae | Aurantimicrobium | 1 |
| surface | Actinobacteriota | Actinobacteria | Micrococcales | Microbacteriaceae | Candidatus Limnoluna | 1 |
| surface | Bacteroidota | Bacteroidia | Cytophagales | Spirosomaceae | Taeseokella | 1 |
| surface | Proteobacteria | Alphaproteobacteria | Caulobacterales | Hyphomonadaceae | Hirschia | 1 |
| surface | Proteobacteria | Alphaproteobacteria | Rhizobiales | Rhizobiales Incertae Sedis | unknown | 1 |
| surface | Proteobacteria | Alphaproteobacteria | Sphingomonadales | Sphingomonadaceae | Rhizorhapis | 1 |
| surface | Proteobacteria | Gammaproteobacteria | Burkholderiales | Alcaligenaceae | GKS98 freshwater group | 1 |
| surface | Proteobacteria | Gammaproteobacteria | Burkholderiales | Comamonadaceae | Rhodoferax | 1 |
| surface | Proteobacteria | Gammaproteobacteria | Burkholderiales | Methylophilaceae | Methylotenera | 1 |

Table S7 – Differentially abundant ASVs in control sites

| Depth | Phylum | Class | Order | Family | Genus | #ASVs |
|---|---|---|---|---|---|---|
| bottom | Bacteroidota | Bacteroidia | Flavobacteriales | Flavobacteriaceae | Flavicella | 1 |
| bottom | Bacteroidota | Bacteroidia | Flavobacteriales | Flavobacteriaceae | NS2b marine group | 1 |
| bottom | Bdellovibrionota | Bdellovibrionia | Bdellovibrionales | Bdellovibrionaceae | OM27 clade | 1 |
| bottom | Myxococcota | Polyangia | Blfdi19 | unknown | unknown | 1 |
| bottom | Nitrospinota | Nitrospinia | Nitrospinales | Nitrospinaceae | Nitrospina | 1 |
| bottom | Proteobacteria | Alphaproteobacteria | Puniceispirillales | unknown | unknown | 1 |
| bottom | Proteobacteria | Alphaproteobacteria | Rhodobacterales | Rhodobacteraceae | Amylibacter | 1 |
| bottom | Proteobacteria | Alphaproteobacteria | Rhodobacterales | Rhodobacteraceae | Planktomarina | 1 |
| bottom | Proteobacteria | Gammaproteobacteria | Pseudomonadales | Porticoccaceae | SAR92 clade | 1 |
| bottom | Proteobacteria | Gammaproteobacteria | Pseudomonadales | Pseudohongiellaceae | Pseudohongiella | 1 |
| bottom | Proteobacteria | Gammaproteobacteria | Pseudomonadales | SAR86 clade | unknown | 1 |
| middle | Bacteroidota | Bacteroidia | Sphingobacteriales | LiUU-11-161 | unknown | 1 |
| middle | Proteobacteria | Alphaproteobacteria | Rhizobiales | Hyphomicrobiaceae | Filomicrobium | 1 |
| surface | Cyanobacteria | Cyanobacteriia | Chloroplast | unknown | unknown | 1 |
| surface | Proteobacteria | Alphaproteobacteria | Defluviicoccales | unknown | unknown | 1 |
| surface | Proteobacteria | Alphaproteobacteria | Rhodospirillales | Magnetospiraceae | Magnetospira | 1 |

Table S8 – Differentially abundant pathways in Lease 1

| Depth | Super.classes | description | ontology | pathway | n |
|---|---|---|---|---|---|
| bottom | Alcohol-Degradation | superpathway of glycerol degradation to 1,3-propanediol | glycerol-deg | alcohol-degradation | 1 |
| bottom | Alcohol-Degradation | superpathway of glycol metabolism and degradation | alcohol-degradatio | alcohol-degradation | 1 |
| bottom | Alcohol-Degradation | superpathway of N-acetylglucosamine, N-acetylmannosamine and N-acetylneuraminate degradation | amine-and-polyamine-degradation | alcohol-degradation | 1 |
| bottom | Alcohol-Degradation | superpathway of ornithine degradation | polyamine-degradation | polyamine-degradation | 1 |
| bottom | Alcohol-Degradation | superpathway of phenylethylamine degradation | amine-deg | alcohol-degradation | 1 |
| bottom | Amino-Acid-Biosynthesis | L-lysine biosynthesis II | lysine-syn | amino-acid-biosynthesis | 1 |

| Depth | Super.classes | description | ontology | pathway | n |
|---|---|---|---|---|---|
| bottom | Amino-Acid-Biosynthesis | L-methionine salvage cycle III | l-methionine-biosynthesis | amino-acid-biosynthesis | 1 |
| bottom | Amino-Acid-Biosynthesis | S-methyl-5-thio-α-D-ribose 1-phosphate degradation | l-methionine-biosynthesis | amino-acid-biosynthesis | 1 |
| bottom | Amino-Acid-Degradation | arginine, ornithine and proline interconversion | arginine-deg | amino-acid-degradation | 1 |
| bottom | Amino-Acid-Degradation | superpathway of L-arginine and L-ornithine degradation | arginine-deg | amino-acid-degradation | 1 |
| bottom | Amino-Acid-Degradation | superpathway of L-arginine, putrescine, and 4-aminobutanoate degradation | arginine-deg | amino-acid-degradation | 1 |
| bottom | Aromatic-compound-degradation | 4-methylcatechol degradation (ortho cleavage) | aromatic-compound-degradation | aromatic-compound-degradation | 1 |
| bottom | Aromatic-compound-degradation | phenylacetate degradation I (aerobic) | phenylacetate-degradation | aromatic-compound-degradation | 1 |
| bottom | Aromatic-compound-degradation | toluene degradation III (aerobic) (via p-cresol) | super-pathway | aromatic-compound-degradation | 1 |
| bottom | Carbohydrates-Degradation | fucose degradation | sugars-and-polysaccharides-degradation | carbohydrate-degradation | 1 |
| bottom | Carbohydrates-Degradation | glucose degradation (oxidative) | sugar-acids-deg | carbohydrate-degradation | 1 |
| bottom | Carbohydrates-Degradation | L-rhamnose degradation I | l-rhamnose-degradation | carbohydrate-degradation | 1 |
| bottom | Carbohydrates-Degradation | mannan degradation | polysaccharides-deg | carbohydrate-degradation | 1 |
| bottom | Cell-Structure/ Cell-Wall-Biosynthesis | polymyxin resistance | antibiotic-resistance | cell-structure/-cell-wall-biosynthesis | 1 |
| bottom | Cofactor-Biosynthesis | cob(II)yrinate a,c-diamide biosynthesis I (early cobalt insertion) | cobyrinate-diamide-biosynthesis | cofactor-biosynthesis | 1 |
| bottom | Cofactor-Biosynthesis | NAD salvage pathway II | nad-syn | cofactor-biosynthesis | 1 |
| bottom | Degradation | superpathway of methylglyoxal degradation | aldehyde-degradation | degradation | 1 |

| Depth | Super.classes | description | ontology | pathway | n |
|---|---|---|---|---|---|
| bottom | Fermentation | acetylene degradation | acetate-formation | fermentation | 1 |
| bottom | Fermentation | glycerol degradation to butanol | alcohol-biosynthesis | fermentation | 1 |
| bottom | Fermentation | hexitol fermentation to lactate, formate, ethanol and acetate | acetate-formation | fermentation | 1 |
| bottom | Fermentation | L-lysine fermentation to acetate and butanoate | acetate-formation | fermentation | 1 |
| bottom | Fermentation | pyruvate fermentation to acetone | pyruvate-degradation | fermentation | 1 |
| bottom | Fermentation | pyruvate fermentation to butanoate | acetyl-coa-butyrate | fermentation | 1 |
| bottom | Fermentation | superpathway of Clostridium acetobutylicum acidogenic fermentation | pyruvate-degradation | fermentation | 1 |
| bottom | Methanogenesis | methanogenesis from acetate | methanogenesis | methanogenesis | 1 |
| bottom | Polyamine-Biosynthesis | superpathway of polyamine biosynthesis III | amine-polyamine-biosynthesis | amine-polyamine-biosynthesis | 1 |
| bottom | Secondary-Metabolite-Biosynthesis | 4-deoxy-L-threo-hex-4-enopyranuronate degradation | sugar-derivatives-degradation | secondary-metabolite-biosynthesis | 1 |
| bottom | Secondary-Metabolite-Biosynthesis | isoprene biosynthesis II (engineered) | isoprenoids | secondary-metabolite-biosynthesis | 1 |
| bottom | Secondary-Metabolite-Biosynthesis | superpathway of (R,R)-butanediol biosynthesis | butanediol-biosynthesis | secondary-metabolite-biosynthesis | 1 |
| bottom | Secondary-Metabolite-Biosynthesis | superpathway of 2,3-butanediol biosynthesis | butanediol-biosynthesis | secondary-metabolite-biosynthesis | 1 |
| bottom | Secondary-Metabolite-Biosynthesis | superpathway of hexitol degradation (bacteria) | sugar-alcohols-deg | secondary-metabolite-biosynthesis | 1 |
| bottom | Sugars-And-Polysaccharides-Degradation | superpathway of N-acetylneuraminate degradation | carboxylates-deg | carbohydrate-degradation | 1 |

| Depth | Super.classes | description | ontology | pathway | n |
|---|---|---|---|---|---|
| bottom | Vitamin-Biosynthesis | adenosylcobalamin biosynthesis I (early cobalt insertion) | de-novo-adenosylcobalamin-biosynthesis | enzyme-cofactor-biosynthesis | 1 |
| bottom | Vitamin-Biosynthesis | adenosylcobalamin biosynthesis II (late cobalt incorporation) | de-novo-adenosylcobalamin-biosynthesis | vitamin-biosynthesis | 1 |
| bottom | Vitamin-Biosynthesis | biotin biosynthesis II | biotin-syn | vitamin-biosynthesis | 1 |
| bottom | Vitamin-Biosynthesis | thiazole biosynthesis I (E. coli) | thiazole-biosynthesis | vitamin-biosynthesis | 1 |
| middle | Alcohol-Degradation | allantoin degradation IV (anaerobic) | allantoin-degradation | alcohol-degradation | 1 |
| middle | Alcohol-Degradation | superpathway of N-acetylglucosamine, N-acetylmannosamine and N-acetylneuraminate degradation | amine-and-polyamine-degradation | alcohol-degradation | 1 |
| middle | Vitamin-Biosynthesis | thiazole biosynthesis II (Bacillus) | thiazole-biosynthesis | vitamin-biosynthesis | 1 |
| surface | Cell-Structure/ Cell-Wall-Biosynthesis | polymyxin resistance | antibiotic-resistance | cell-structure/-cell-wall-biosynthesis | 1 |
| surface | Degradation | vitamin B6 degradation | vitamin-b6-degradation | vitamin-degradation | 1 |

Table S9 – Differentially abundant pathways in control site

| Depth | Super.classes | description | ontology | pathway | n |
|---|---|---|---|---|---|
| bottom | Secondary-Metabolite-Biosynthesis | spirilloxanthin and 2,2'-diketo-spirilloxanthin biosynthesis | c40-carotenoids-biosynthesis | secondary-metabolite-biosynthesis | 1 |
| middle | lipid-biosynthesis | superpathway of fatty acids biosynthesis (E. coli) | fatty-acid-and-lipid-biosynthesis | lipid-biosynthesis | 1 |
| middle | lipid-biosynthesis | superpathway of mycolate biosynthesis | fatty-acid-and-lipid-biosynthesis | lipid-biosynthesis | 1 |

Table S10 – The 20 most important ASVs in predicting the environmental drivers of the bacterial community.

| ASV | Importance | Kingdom | Phylum | Class | Order | Family | Genus | env_par |
|---|---|---|---|---|---|---|---|---|
| **b_asv_1104** | 6.64 | Bacteria | Actinobacteriota | Actinobacteria | Frankiales | Sporichthyaceae | hgcI clade | NH4 |
| **b_asv_123** | 5.33 | Bacteria | Proteobacteria | Alphaproteobacteria | Kiloniellales | Kiloniellaceae | unknown | NH4 |
| **b_asv_1105** | 5.07 | Bacteria | Proteobacteria | Gammaproteobacteria | Burkholderiales | Comamonadaceae | Rhodoferax | NH4 |

| ASV | Importance | Kingdom | Phylum | Class | Order | Family | Genus | env_par |
|---|---|---|---|---|---|---|---|---|
| **b_asv_1492** | 4.96 | Bacteria | Proteobacteria | Alphaproteobacteria | Rhizobiales | Rhizobiales Incertae Sedis | unknown | NH4 |
| **b_asv_739** | 4.52 | Bacteria | Proteobacteria | Alphaproteobacteria | Caulobacterales | Hyphomonadaceae | Hirschia | NH4 |
| **b_asv_164** | 4.44 | Bacteria | Proteobacteria | Gammaproteobacteria | Pseudomonadales | Pseudohongiellaceae | Pseudohongiella | NH4 |
| **b_asv_18** | 4.00 | Bacteria | Proteobacteria | Alphaproteobacteria | SAR11 clade | Clade III | unknown | NH4 |
| **b_asv_23** | 3.88 | Bacteria | Proteobacteria | Gammaproteobacteria | Burkholderiales | Comamonadaceae | RS62 marine group | NH4 |
| **b_asv_30** | 3.76 | Bacteria | Bacteroidota | Bacteroidia | Cytophagales | Spirosomaceae | Taeseokella | NH4 |
| **b_asv_171** | 3.69 | Bacteria | Proteobacteria | Alphaproteobacteria | SAR11 clade | Clade I | Clade Ib | NH4 |
| **b_asv_539** | 3.65 | Bacteria | Proteobacteria | Alphaproteobacteria | Rhodospirillales | Magnetospiraceae | unknown | NH4 |
| **b_asv_115** | 3.61 | Bacteria | Proteobacteria | Alphaproteobacteria | Rhodobacterales | Rhodobacteraceae | unknown | NH4 |
| **b_asv_1327** | 3.57 | Bacteria | Bacteroidota | Bacteroidia | Flavobacteriales | Flavobacteriaceae | Flavobacterium | NH4 |
| **b_asv_151** | 3.53 | Bacteria | Proteobacteria | Alphaproteobacteria | Puniceispirillales | unknown | unknown | NH4 |
| **b_asv_283** | 3.52 | Bacteria | Proteobacteria | Alphaproteobacteria | Rhizobiales | Hyphomicrobiaceae | Filomicrobium | NH4 |
| **b_asv_331** | 3.46 | Bacteria | Proteobacteria | Gammaproteobacteria | Pseudomonadales | Porticoccaceae | C1-B045 | NH4 |
| **b_asv_672** | 3.45 | Bacteria | Bacteroidota | Bacteroidia | Flavobacteriales | Flavobacteriaceae | NS3a marine group | NH4 |
| **b_asv_832** | 3.32 | Bacteria | Proteobacteria | Gammaproteobacteria | Burkholderiales | Methylophilaceae | Methylotenera | NH4 |
| **b_asv_299** | 3.31 | Bacteria | Bacteroidota | Bacteroidia | Flavobacteriales | Crocinitomicaceae | Fluviicola | NH4 |
| **b_asv_9** | 3.29 | Bacteria | Proteobacteria | Alphaproteobacteria | SAR11 clade | Clade III | unknown | NH4 |
| **b_asv_327** | 4.80 | Bacteria | Bdellovibrionota | Oligoflexia | Oligoflexales | Oligoflexaceae | unknown | NOX |
| **b_asv_1** | 4.20 | Bacteria | Proteobacteria | Gammaproteobacteria | Pseudomonadales | Thioglobaceae | Candidatus Thioglobus | NOX |
| **b_asv_447** | 4.05 | Bacteria | Bdellovibrionota | Bdellovibrionia | Bdellovibrionales | Bdellovibrionaceae | OM27 clade | NOX |
| **b_asv_11** | 4.02 | Bacteria | Proteobacteria | Gammaproteobacteria | Burkholderiales | Comamonadaceae | RS62 marine group | NOX |
| **b_asv_115** | 3.93 | Bacteria | Proteobacteria | Alphaproteobacteria | Rhodobacterales | Rhodobacteraceae | unknown | NOX |
| **b_asv_23** | 3.66 | Bacteria | Proteobacteria | Gammaproteobacteria | Burkholderiales | Comamonadaceae | RS62 marine group | NOX |
| **b_asv_147** | 3.60 | Bacteria | Bacteroidota | Bacteroidia | Sphingobacteriales | LiUU-11-161 | unknown | NOX |

| ASV | Importance | Kingdom | Phylum | Class | Order | Family | Genus | env_par |
|---|---|---|---|---|---|---|---|---|
| **b_asv_13** | 3.57 | Bacteria | Planctomycetota | Planctomycetes | Planctomycetales | Gimesiaceae | unknown | NOX |
| **b_asv_102** | 3.57 | Bacteria | Bdellovibrionota | unknown | unknown | unknown | unknown | NOX |
| **b_asv_180** | 3.55 | Bacteria | Dadabacteria | Dadabacteriia | Dadabacteriales | unknown | unknown | NOX |
| **b_asv_27** | 3.48 | Bacteria | Patescibacteria | Gracilibacteria | Candidatus Peregrinibacteria | unknown | unknown | NOX |
| **b_asv_55** | 3.36 | Bacteria | Proteobacteria | Alphaproteobacteria | Rhodobacterales | Rhodobacteraceae | Amylibacter | NOX |
| **b_asv_20** | 3.36 | Bacteria | Bacteroidota | Bacteroidia | Flavobacteriales | Flavobacteriaceae | NS3a marine group | NOX |
| **b_asv_149** | 3.33 | Bacteria | Proteobacteria | Alphaproteobacteria | Rhodobacterales | Rhodobacteraceae | unknown | NOX |
| **b_asv_944** | 3.29 | Bacteria | Proteobacteria | Alphaproteobacteria | Rickettsiales | unknown | unknown | NOX |
| **b_asv_319** | 3.22 | Bacteria | Cyanobacteria | Cyanobacteriia | Chloroplast | unknown | unknown | NOX |
| **b_asv_30** | 3.21 | Bacteria | Bacteroidota | Bacteroidia | Cytophagales | Spirosomaceae | Taeseokella | NOX |
| **b_asv_146** | 3.17 | Bacteria | Cyanobacteria | Cyanobacteriia | Chloroplast | unknown | unknown | NOX |
| **b_asv_194** | 3.14 | Bacteria | Proteobacteria | Alphaproteobacteria | Rhodobacterales | Rhodobacteraceae | Ascidiaceihabitans | NOX |
| **b_asv_2** | 3.11 | Bacteria | Campylobacterota | Campylobacteria | Campylobacterales | Arcobacteraceae | unknown | NOX |
| **b_asv_230** | 12.96 | Bacteria | Proteobacteria | Gammaproteobacteria | Burkholderiales | Methylophilaceae | OM43 clade | turbidity |
| **b_asv_724** | 10.21 | Bacteria | Planctomycetota | Planctomycetes | Planctomycetales | Rubinisphaeraceae | Planctomicrobium | turbidity |
| **b_asv_1275** | 6.83 | Bacteria | Proteobacteria | Alphaproteobacteria | Sphingomonadales | Sphingomonadaceae | Rhizorhapis | turbidity |
| **b_asv_546** | 5.30 | Bacteria | Proteobacteria | Gammaproteobacteria | Pseudomonadales | SAR86 clade | unknown | turbidity |
| **b_asv_691** | 5.27 | Bacteria | Bacteroidota | Bacteroidia | Flavobacteriales | Flavobacteriaceae | NS2b marine group | turbidity |
| **b_asv_55** | 4.50 | Bacteria | Proteobacteria | Alphaproteobacteria | Rhodobacterales | Rhodobacteraceae | Amylibacter | turbidity |
| **b_asv_327** | 3.78 | Bacteria | Bdellovibrionota | Oligoflexia | Oligoflexales | Oligoflexaceae | unknown | turbidity |
| **b_asv_426** | 3.52 | Bacteria | Myxococcota | Polyangia | Polyangiales | Sandaracinaceae | Sandaracinus | turbidity |
| **b_asv_4** | 2.97 | Bacteria | Proteobacteria | Alphaproteobacteria | SAR11 clade | Clade I | Clade Ia | turbidity |
| **b_asv_180** | 2.70 | Bacteria | Dadabacteria | Dadabacteriia | Dadabacteriales | unknown | unknown | turbidity |
| **b_asv_2** | 2.59 | Bacteria | Campylobacterota | Campylobacteria | Campylobacterales | Arcobacteraceae | unknown | turbidity |
| **b_asv_739** | 2.35 | Bacteria | Proteobacteria | Alphaproteobacteria | Caulobacterales | Hyphomonadaceae | Hirschia | turbidity |
| **b_asv_218** | 2.33 | Bacteria | Bacteroidota | Bacteroidia | Flavobacteriales | Flavobacteriaceae | Flavicella | turbidity |

| ASV | Importance | Kingdom | Phylum | Class | Order | Family | Genus | env_par |
|---|---|---|---|---|---|---|---|---|
| **b_asv_1127** | 2.31 | Bacteria | Proteobacteria | Gammaproteobacteria | Pseudomonadales | Moraxellaceae | Acinetobacter | turbidity |
| **b_asv_11** | 2.24 | Bacteria | Proteobacteria | Gammaproteobacteria | Burkholderiales | Comamonadaceae | RS62 marine group | turbidity |
| **b_asv_194** | 2.22 | Bacteria | Proteobacteria | Alphaproteobacteria | Rhodobacterales | Rhodobacteraceae | Ascidiaceihabitans | turbidity |
| **b_asv_115** | 2.01 | Bacteria | Proteobacteria | Alphaproteobacteria | Rhodobacterales | Rhodobacteraceae | unknown | turbidity |
| **b_asv_171** | 1.88 | Bacteria | Proteobacteria | Alphaproteobacteria | SAR11 clade | Clade I | Clade Ib | turbidity |
| **b_asv_462** | 1.79 | Bacteria | Proteobacteria | Alphaproteobacteria | Rhodobacterales | Rhodobacteraceae | Planktomarina | turbidity |
| **b_asv_865** | 1.69 | Bacteria | Proteobacteria | Gammaproteobacteria | Enterobacterales | Vibrionaceae | Aliivibrio | turbidity |

Table S11 – The 20 most important pathways in predicting the environmental drivers of the bacterial community.

| ASV | Importance | description | pathway | ontology | Super.classes | env_par |
|---|---|---|---|---|---|---|
| **PWY-6891** | 8.64 | thiazole biosynthesis II (Bacillus) | vitamin-biosynthesis | thiazole-biosynthesis | Vitamin-Biosynthesis | NOX |
| **PWY-6895** | 7.69 | superpathway of thiamin diphosphate biosynthesis II | vitamin-biosynthesis | super-pathway | Vitamin-Biosynthesis | NOX |
| **METH-ACETATE-PWY** | 7.50 | methanogenesis from acetate | methanogenesis | methanogenesis | Methanogenesis | NOX |
| **PWY-5860** | 6.86 | superpathway of demethylmenaquinol-6 biosynthesis I | vitamin-biosynthesis | demethylmenaquinol-6-biosynthesis | Vitamin-Biosynthesis | NOX |
| **PWY-5100** | 6.68 | pyruvate fermentation to acetate and lactate II | fermentation | pyruvate-acetate-fermentation | Fermentation | NOX |
| **PWY-7315** | 5.04 | dTDP-N-acetylthomosamine biosynthesis | carbohydrate-biosynthesis | dtdp-sugar-biosynthesis | Carbohydrates-Biosynthesis | NOX |
| **GLUCUROCAT-PWY** | 4.97 | superpathway of β-D-glucuronide and D-glucuronate degradation | sugars-and-acids-degradation | d-glucuronate-degradation | Sugars-And-Acids-Degradation | NOX |
| **PWY-6185** | 4.58 | 4-methylcatechol degradation (ortho cleavage) | aromatic-compound-degradation | aromatic-compound-degradation | Aromatic-compound-degradation | NOX |
| **PWY-5181** | 4.45 | toluene degradation III (aerobic) (via p-cresol) | aromatic-compound-degradation | super-pathway | Aromatic-compound-degradation | NOX |

| ASV | Importance | description | pathway | ontology | Super.classes | env_par |
|---|---|---|---|---|---|---|
| **PWY-5531** | 4.27 | chlorophyllide a biosynthesis II (anaerobic) | chlorophyll-biosynthesis | chlorophyllide-a-biosynthesis | Chlorophyll-Biosynthesis | NOX |
| **PWY-7007** | 4.23 | methyl ketone biosynthesis | energy-metabolism | energy-metabolism | Energy-Metabolism | NOX |
| **HOMOSER-METSYN-PWY** | 4.01 | L-methionine biosynthesis I | amino-acid-biosynthesis | methionine-de-novo-biosynthesis | Amino-Acid-Biosynthesis | NOX |
| **ORNDEG-PWY** | 3.93 | superpathway of ornithine degradation | polyamine-degradation | polyamine-degradation | Alcohol-Degradation | NOX |
| **PWY-6892** | 3.83 | thiazole biosynthesis I (E. coli) | vitamin-biosynthesis | thiazole-biosynthesis | Vitamin-Biosynthesis | NOX |
| **PWY-7159** | 3.77 | chlorophyllide a biosynthesis III (aerobic, light independent) | chlorophyll-biosynthesis | chlorophyllide-a-biosynthesis | Chlorophyll-Biosynthesis | NOX |
| **CHLOROPHYLL-SYN** | 3.57 | chlorophyllide a biosynthesis I (aerobic, light-dependent) | chlorophyll-biosynthesis | chlorophyllide-a-biosynthesis | Chlorophyll-Biosynthesis | NOX |
| **PWY0-321** | 3.21 | phenylacetate degradation I (aerobic) | aromatic-compound-degradation | phenylacetate-degradation | Aromatic-compound-degradation | NOX |
| **ASPASN-PWY** | 3.11 | superpathway of L-aspartate and L-asparagine biosynthesis | amino-acid-biosynthesis | amino-acid-biosynthesis | Amino-Acid-Biosynthesis | NOX |
| **PWY-5347** | 2.97 | superpathway of L-methionine biosynthesis (transsulfuration) | amino-acid-biosynthesis | methionine-de-novo-biosynthesis | Amino-Acid-Biosynthesis | NOX |
| **PWY-7187** | 2.97 | pyrimidine deoxyribonucleotides de novo biosynthesis II | nucleotide-biosynthesis | pyrimid-deoxyribonucleot-de-novo-biosyn | Nucleotide-Biosynthesis | NOX |
| **GALACT-GLUCUROCAT-PWY** | 9.71 | superpathway of hexuronide and hexuronate degradation | generalized-reactions | super-pathway | Generalized-Reactions | turbidity |
| **P381-PWY** | 7.90 | adenosylcobalamin biosynthesis II (late cobalt incorporation) | vitamin-biosynthesis | de-novo-adenosylcobalamin-biosynthesis | Vitamin-Biosynthesis | turbidity |
| **P161-PWY** | 5.79 | acetylene degradation | fermentation | acetate-formation | Fermentation | turbidity |

| ASV | Importance | description | pathway | ontology | Super.classes | env_par |
|---|---|---|---|---|---|---|
| **GLUCOSE1PMETAB-PWY** | 5.52 | glucose and glucose-1-phosphate degradation | carbohydrate-degradation | sugars-and-polysaccharides-degradation | Carbohydrates-Degradation | turbidity |
| **NAD-BIOSYNTHESIS-II** | 5.39 | NAD salvage pathway II | cofactor-biosynthesis | nad-syn | Cofactor-Biosynthesis | turbidity |
| **PWY-7377** | 5.01 | cob(II)yrinate a,c-diamide biosynthesis I (early cobalt insertion) | cofactor-biosynthesis | cobyrinate-diamide-biosynthesis | Cofactor-Biosynthesis | turbidity |
| **PWY-5507** | 4.98 | adenosylcobalamin biosynthesis I (early cobalt insertion) | enzyme-cofactor-biosynthesis | de-novo-adenosylcobalamin-biosynthesis | Vitamin-Biosynthesis | turbidity |
| **PWY-5384** | 4.17 | sucrose degradation IV (sucrose phosphorylase) | carbohydrate-degradation | sucrose-deg | Sugars-And-Polysaccharides-Degradation | turbidity |
| **PWY-6565** | 4.06 | superpathway of polyamine biosynthesis III | amine-polyamine-biosynthesis | amine-polyamine-biosynthesis | Polyamine-Biosynthesis | turbidity |
| **PWY-5181** | 3.87 | toluene degradation III (aerobic) (via p-cresol) | aromatic-compound-degradation | super-pathway | Aromatic-compound-degradation | turbidity |
| **P125-PWY** | 3.73 | superpathway of (R,R)-butanediol biosynthesis | secondary-metabolite-biosynthesis | butanediol-biosynthesis | Secondary-Metabolite-Biosynthesis | turbidity |
| **GLUCUROCAT-PWY** | 3.64 | superpathway of β-D-glucuronide and D-glucuronate degradation | sugars-and-acids-degradation | d-glucuronate-degradation | Sugars-And-Acids-Degradation | turbidity |
| **PWY-6396** | 3.51 | superpathway of 2,3-butanediol biosynthesis | secondary-metabolite-biosynthesis | butanediol-biosynthesis | Secondary-Metabolite-Biosynthesis | turbidity |
| **PWY-621** | 3.47 | sucrose degradation III (sucrose invertase) | carbohydrate-degradation | sucrose-deg | Carbohydrates-Degradation | turbidity |
| **PWY0-321** | 3.43 | phenylacetate degradation I (aerobic) | aromatic-compound-degradation | phenylacetate-degradation | Aromatic-compound-degradation | turbidity |

| ASV | Importance | description | pathway | ontology | Super.classes | env_par |
|---|---|---|---|---|---|---|
| **PWY-6891** | 3.29 | thiazole biosynthesis II (Bacillus) | vitamin-biosynthesis | thiazole-biosynthesis | Vitamin-Biosynthesis | turbidity |
| **PWY-6895** | 3.07 | superpathway of thiamin diphosphate biosynthesis II | vitamin-biosynthesis | super-pathway | Vitamin-Biosynthesis | turbidity |
| **ASPASN-PWY** | 3.01 | superpathway of L-aspartate and L-asparagine biosynthesis | amino-acid-biosynthesis | amino-acid-biosynthesis | Amino-Acid-Biosynthesis | turbidity |
| **PWY-7254** | 2.98 | TCA cycle VII (acetate-producers) | energy-metabolism | tca-variants | Energy-Metabolism | turbidity |
| **METH-ACETATE-PWY** | 2.91 | methanogenesis from acetate | methanogenesis | methanogenesis | Methanogenesis | turbidity |
